# Proteomic Investigation of Neurotrophic *trans*-Banglene Reveals Potential Link to Iron Homeostasis

**DOI:** 10.1101/2024.09.04.611284

**Authors:** Piyumi B. Wijesiri Gunawardana, Khyati Gohil, Kyung-Mee Moon, Leonard J. Foster, Florence J. Williams

## Abstract

In an effort to gain insight into cellular systems impacted by neurotrophic *trans*-banglene (*t-*BG), global proteomic profiling and Western blot analyses were employed. Expression level changes in response to *t*-BG treatment were compared to those observed with nerve growth factor (NGF), a natural neurotrophic protein and functional analog to *t*-BG. Findings from these studies did not point to direct interception of NGF/TrkA signaling by *t-*BG. Instead, significant alterations in iron-binding and iron-regulating proteins were observed. Intracellular iron measurements by FerroOrange indicate lower ferrous (Fe^2+^) iron levels in *t*-BG treated cells but not in NGF treated cells. These results highlight a potential connection between iron regulation and neurotrophic activity.

## INTRODUCTION

Neurodegenerative diseases remain an area where the causative disease mechanisms, with few exceptions, are poorly understood and which are very resistant to modern therapeutic advances. Current FDA-approved medications result in modest improvements in quality of life measures and no increase in life expectancy [1]. In this context, insight into cellular elements that dictate neuronal survival and growth versus degradation or death (the shared feature of all neurodegenerative disorders), is valuable. Molecular agents that modulate these responses can be uniquely beneficial in such inquiries because of their ease of administration, frequent robustness in biological media, and their potential for pharmacokinetic optimization or derivatization in the hands of a medicinal chemist. Further, investigation of the mechanism of action for small molecule natural products can lead to unexpected insights, as was the case in the serendipitous discovery of the mTOR protein (mammalian target of rapamycin), a central controller of cell metabolism [2]. *t*-BG is a small molecule neurotrophic agent that is stable in serum and blood [3]. This is a distinction from neurotrophin proteins (the body’s natural neurotrophic agents), which have short half-lives in serum (< 1 minute).

*t*-BG is readily accessible (2 or 3 synthetic steps from inexpensive materials, depending on route preference) [4], and it exhibits neurotrophic activity in PC-12 cells (neuronal model cell line), primary rat hippocampal cells, and mouse models. *t*-BG has been shown to be orally bioavailable in mice and to cross the blood-brain barrier, and has been demonstrated to increase neurogenesis in a surgical neurodegenerative mouse model (olfactory bulbectomy) [5]. Despite these compelling phenotypic results, a cellular target and mechanism of action remain unknown. We aim to close this knowledge gap.

Identifying mechanism of action with only phenotypic response data is challenging because of the many signaling pathways that could result in similar phenotypes. To provide early insight into which cellular systems are impacted by *t*-BG, we used global proteomic profiling.

We performed a global proteomic analysis of *t*-BG’s effects in PC-12 cells and compared it to the impact of natural NGF, a neurotrophin protein which causes similar phenotypic changes in PC-12 cells. To our surprise, while we gained insight into classic pro-growth, pro-differentiation, and pro-survival pathways, we also observed changes in iron-binding proteins in the cell that were unanticipated. Since iron dyshomeostasis is an increasingly recognized element of neurodegenerative disease, particularly in early stages of Alzheimer’s and Parkinson’s disease [6,7], the potential connection between neurotrophic activity and altered iron homeostasis is quite notable [8-12].

## RESULTS AND DISCUSSION

### Proteomics Results

Our global proteomic profile of PC-12 responses compared *t*-BG treatment and NGF treatment to a control (DMSO alone, which was the solvent used to administer *t*-BG and was added to the NGF treatment samples). Looking at a heatmap of hierarchical clustering of NGF and *t*-BG treatment responses (Figure 1(a)), along with Volcano plots showing the visualization of significant protein expression changes in red dots (>2 fold, p<0.05, Figure 1(b)), we observe both significant similarity between NGF and *t*-BG treatment, and some key regions of difference. Out of 3176 proteins investigated, 236 proteins were identified showing statistically significant changes (> 2 fold, p < 0.05) in at least one of the treatment groups (Figure 1 (c)) ((see supplementary information, which include Benjamini-Hochberg values listed as q-values).

**Figure 1:**
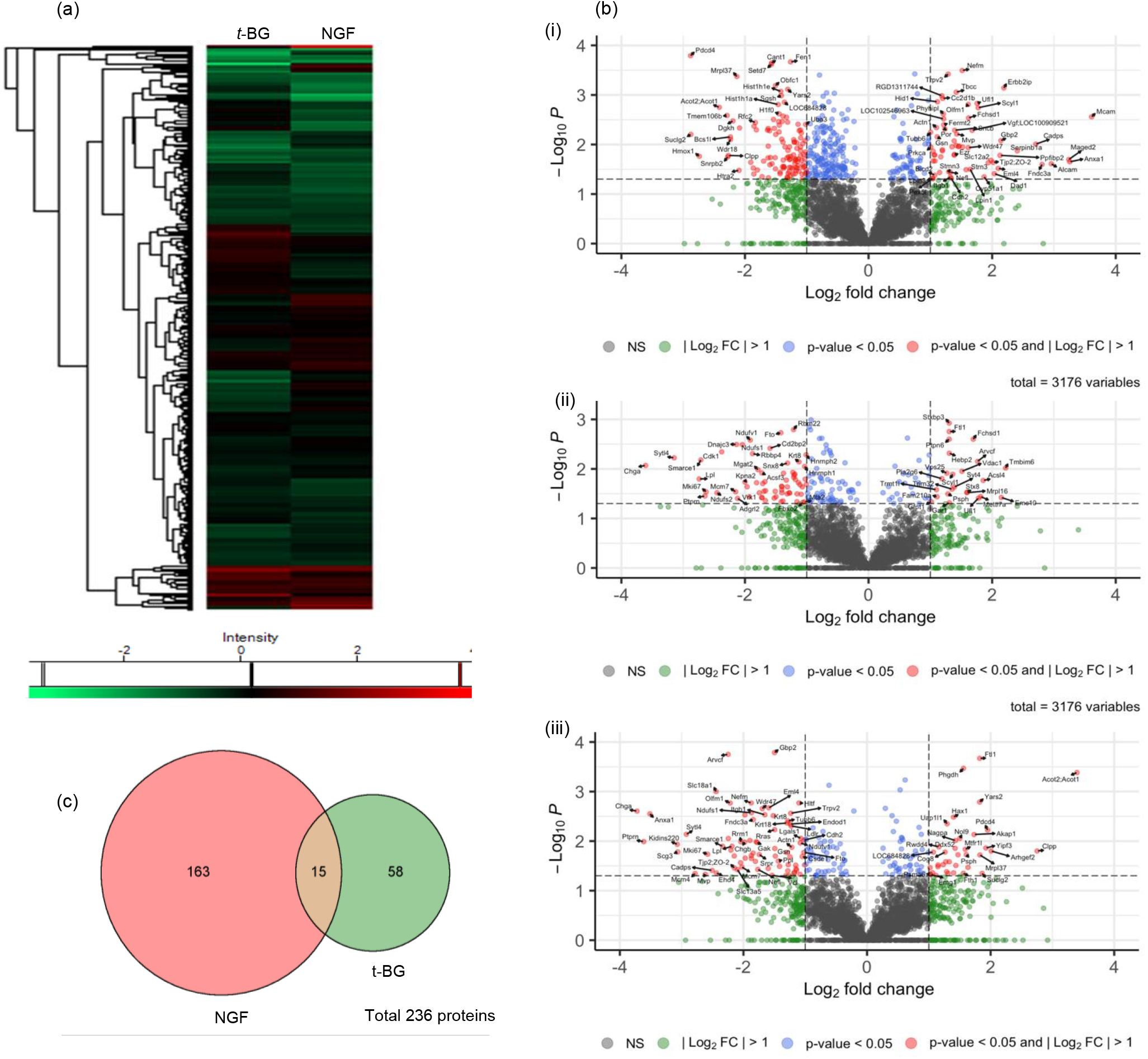
Differential protein expression analysis of proteomics data derived from label-free shotgun proteomics (Perseus was used to create the heat plot) (a) Dendrogram heatmap of protein expression for log_2_ fold change (LFC) of NGF and *t*-BG treatment (b) Volcano plots of proteins identified by global proteomic analysis (EnhancedVolcano was used to create volcano plots) (i) NGF vs control (ii) *t*-BG vs control (iii) *t*-BG vs NGF (c) Venn diagram of statistically significant proteins identified by global proteomic analysis (>2 fold, p<0.05) of NGF and *t*-BG treatment (SRplots were used to create the Venn diagram). Enlarged versions of the Volcano plots are provided in the supplemental information.

NGF treatment resulted in significant protein level changes in more of these proteins (178) as compared to *t*-BG treatment (73) (Figure 1(c)), with 15 altered in both treatment regimes. Of the significantly altered proteins for both NGF and *t*-BG treatments, those that were involved in replication, chromatin structure, transcription, rRNA and tRNA processing, mitochondrial respiratory chain complex I proteins and nucleosome assembly proteins were generally downregulated (see SI). Given that NGF and *t*-BG are pro-growth and pro-neuritogenesis/differentiation effectors, downregulation of these targets is likely arising from a feedback response following a major growth and/or differentiation event. These samples were taken after 48 hours of treatment, after most of the differentiation, cell body area growth, and proliferation has occurred.

To gain insight into signal transduction pathways that are affected by *t*-BG treatment, we used DAVID 6.8 (Database for Annotation, Visualization and Integrated Discovery) to analyze the altered proteins with respect to annotated KEGG pathways (See SI). Ordering by count and removing entries with p > 0.05, we see that the top four KEGG pathways identified involve neurodegenerative diseases (see Table 1).

**Table 1:** KEGG pathway analysis by DAVID knowledgebase for altered proteins with *t*-BG treatment.

| Entry | KEGG Term | Count | p-Value | FDR |
| --- | --- | --- | --- | --- |
| 1 | Amyotrophic Lateral Sclerosis | 13 | 0.000498 | 0.0458 |
| 2 | Alzheimer's Disease | 11 | 0.00795 | 0.133 |
| 3 | Parkinson's Disease | 10 | 0.00213 | 0.0782 |
| 4 | Prion Disease | 9 | 0.00787 | 0.133 |
| 5 | Spliceosome | 8 | 0.0126 | 0.165 |

For the top entry, Amyotrophic Lateral Sclerosis, nearly a third of proteins identified from our altered protein list are iron-binding proteins (Ndufs1, Ndufs2, Ndufv1, Cat). The remaining proteins involve the proteasome (Psmc3, Psmc4, Psmc5), nuclear transport (Nxf1, Nup54, Nup88), phospholipid binding (Anxa11), tubulin (Tubb5), and ion transport (Vdac1). KEGG pathway analysis of proteins altered with NGF treatment identified similar affected neurodegenerative pathways.

We next used DAVID analysis to look at the differences between *t*-BG and NGF, namely looking at which proteins are altered in *t*-BG as compared to NGF treatment. Because of the similar outcomes for *t*-BG and NGF treatment groups when using KEGG pathway analysis, for this assessment we used a wide array of annotation categories and used DAVID annotation clustering to look for related features identified from different database (category) assessments. Using all default selection criteria, the top hit/annotation cluster focused on cellular processes associated with or utilizing iron (See Table 2). This was unexpected and led us to consider the frequency with which we see iron-binding proteins in our data sets, and the identity and role of those proteins.

**Table 2:** Annotation cluster 1 from DAVID knowledgebase analysis for *t*-BG treatment as compared to NGF treatment.

| Annotation Cluster 1 - Enrichment Score: 2.75 |  |  |  |
| --- | --- | --- | --- |
| Category | Term | p-Value | FDR |
| GOTERM-BP Direct | Cellular Iron Homeostasis | 0.000587 | 0.137 |
| INTERPRO | Ferritin_CS | 0.000622 | 0.0165 |
| GOTERM-BP-Direct | Intracellular sequestering of iron | 0.000778 | 0.137 |
| UP-SEQ-Feature | Ferritin-like diiron | 0.000989 | 0.0500 |
| INTERPRO | Ferritin | 0.001082 | 0.0165 |
| INTERPRO | Ferritin_DPS_dom | 0.001082 | 0.0165 |
| INTERPRO | Ferritin-like | 0.001171 | 0.0165 |
| INTERPRO | Ferritin-like_SF | 0.001458 | 0.0190 |
| GOTERM-MF Direct | Ferric iron binding | 0.00258 | 0.119 |
| GOTERM-MF Direct | Ferrous iron binding | 0.0041 | 0.119 |
| GOTERM-BP Direct | Iron ion transport | 0.00534 | 0.564 |
| UP-KW-Ligand | Iron | 0.0476 | 0.301 |

We next focused on specific protein changes; we closely evaluated two groups: NGF/TrkA related proteins and iron-binding and storage proteins, given the DAVID results.

NGF/Trk signal transduction is responsible for the cell’s neurotrophic response to NGF [13]. Due to the similarity of function between *t*-BG and NGF, it was important to look for evidence of *t*-BG intercepting similar NGF-mediated neurotrophic processes (See Figure 2).

**Figure 2:**
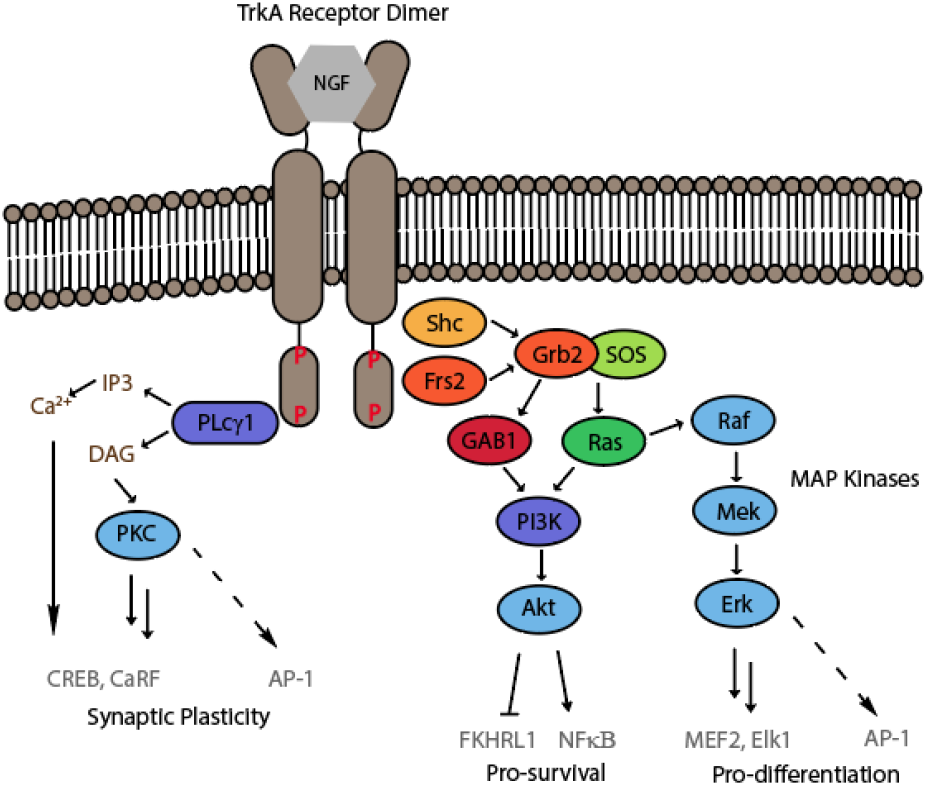
TrkA/NGF signaling [14]

Overall, the majority of proteins in the NGF/TrkA signaling pathway, for both NGF and *t*-BG treatment groups, showed limited expression level changes. These limited expression level changes are in line with the primary mechanism of NGF/Trk signal transduction occurring through post-translational modification (e.g. phosphorylation) rather than impacts on expression levels [13]. Therefore, it is difficult with this data set to determine whether *t*-BG is intercepting and/or promoting NGF/Trk signal transduction directly. However, we do find evidence that Trk signaling is impacted by *t*-BG treatment.

In our dataset, proteins in the TrkA/NGF pathway, or which are intimately connected to that pathway, that were significantly altered by either NGF or *t*-BG are: protein kinase C α (Pkcα), protein interacting with C kinase 1 (Pick1), Diacylglycerol kinase eta (Dgkh), tripartite motif 32 (Trim32), Protein kinase R (PKR), Src homology region 2 domain-containing phosphatase-1 (SHP-1)), N-cadherin, activated leucocyte cell adhesion molecule (Alcam), β-synuclein, and murine serpin family B member 1 (Serpinb1a), which are highlighted in Figure 3.

**Figure 3:**
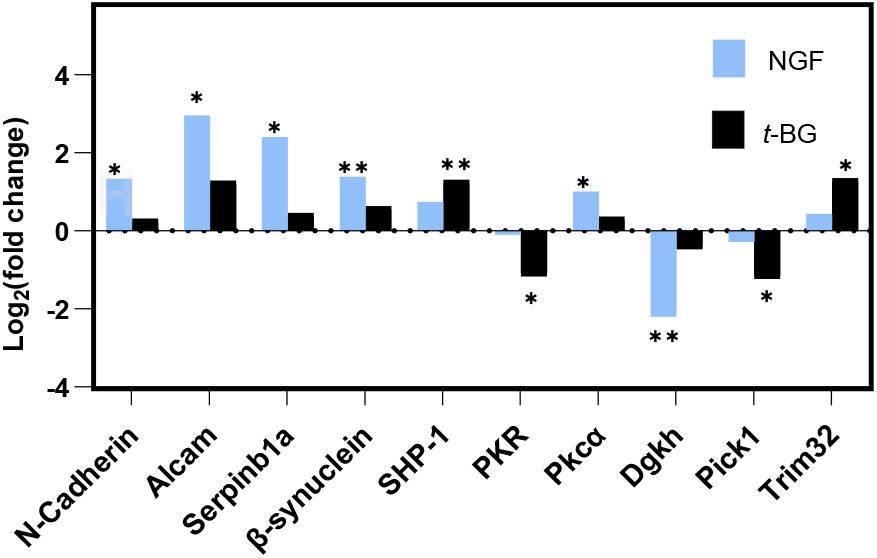
NGF/Trk signaling related expression of significant protein fold change (>2 fold, p<0.05) compared to control *p<0.05, **p<0.01, ***p<0.001.

#### NGF-Altered TrkA Signaling Proteins

N-cadherin is increased in expression upon NGF treatment, which is interesting. While N-cadherin is not known to directly activate NGF/Trk signaling, it plays a crucial role in synapse formation via the Cdh2/Iqgap1/Erk-2 pathway and in neuronal differentiation through PI3K/Akt/β-catenin signaling independent of Wnt [15-17].

The observed elevated levels of activated leukocyte cell adhesion molecule (Alcam) with NGF treatment is part of the cell’s feedback responses to NGF. Alcam potentiates NGF activity by enhancing TrkA autophosphorylation, thereby enhancing TrkA/NGF signaling effects [18]. Additionally, Alcam is co-transported with low-affinity nerve growth factor receptor (p75NTR) towards the cell body in motor neuron axons [18]. p75NTR has an integral impact on TrkA/NGF signaling and can be augmenting or pro-apoptotic in effect [19, 20].

NGF treatment further resulted in upregulation of murine serpin family B member 1 (Serpinb1A), a homologue of human Serpinb1, and β-synuclein. Both of these proteins regulate key drivers of TrkA/NGF signal transduction. Serpinb1A exerts an anti-apoptotic effect by regulating the activity of Erk, p38 and PI3k [21]. β-synuclein is a presynaptic protein which exhibits neuroprotection by upregulating phosphorylated Akt [22]. Importantly, β-synuclein has been suggested as an inhibitor to α-synuclein aggregation, which results in Lewy bodies in Parkinson’s disease [22].

There was also an increase in the level of Pkcα in the NGF treated samples. Pkcα activation involves the generation of diacylgycerol (DAG), a compound that activates downstream effectors, including other Pkc isoforms, to maintain neuronal synaptic plasticity [23]. Related, Dgkh is downregulated in this sample group and is a major controller of DAG in the cell. Dgkh is a critical component of the Ras/Raf/Mek/Erk signaling pathway for cell proliferation, differentiation and survival, as it enhances the heterodimerization of b-Raf and c-Raf and subsequent activation of Mek and Erk [24].

#### t-BG Altered TrkA Signaling Proteins

Though the observed increase in Cdh2, Alcam, Serpinb1a and β-synuclein are statistically insignificant for *t*-BG treatment samples, SHP-1 is notably upregulated. SHP-1 is a phosphotyrosine phosphatase that negatively regulates TrkA, and the inhibition of SHP-1 results in activation of Akt [25]. Downregulation of PKR also affects Akt activity. PKR is a transcription factor that acts as a positive regulator for Akt by increasing Akt phosphorylation [26]. Similarly, though the observed increase in Pkcα is modest and statistically insignificant for *t*-BG treatment samples, Pick1, a Pkcα-modulating protein, is downregulated. Pick1 protein complexes and activates Pkcα to mediate Pkc-regulated synaptic transmission [27]. Trim32 is significantly upregulated in *t*-BG treated samples; Trim32 is known to bind with Pkcζ and inhibit neuronal differentiation [28].

Collectively, these responses to *t*-BG treatment mitigate the action of NGF/TrkA signal transduction. These changes may be a negative feedback response to the differentiation-inducing effects of *t*-BG.

#### Altered Iron-Binding Proteins

Looking further at the proteomic data, an unexpected trend emerged – several proteins involved in iron storage, or which utilize and bind to iron (e.g. iron-sulfur cluster proteins, heme proteins) were altered in expression level (See Figure 4). Most dramatically upregulated with *t*-BG treatment is ferritin light chain (Ftl), which, along with ferritin heavy chain, plays a central role in iron storage in the cytosol and in modulating iron homeostasis. Unlike ferritin heavy chain, ferritin light chain’s only known role is the storage of Fe^3+^; it is not thought to be involved in the reduction of Fe^3+^ to Fe^2+^.

**Figure 4:**
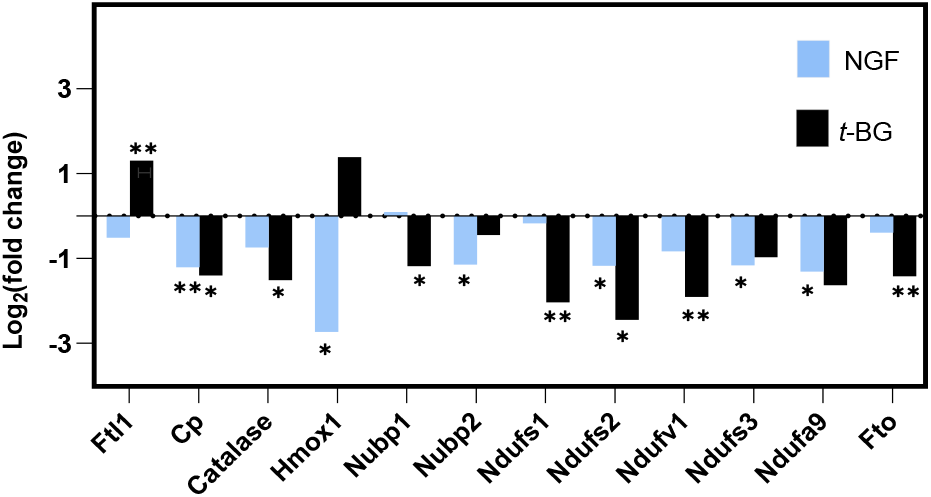
Iron homeostasis related expression of significant protein fold change (>2 fold, p<0.05) compared to control *p<0.05, **p<0.01.

Other proteins that either regulate iron oxidation levels or the utilization of iron in the cytoplasm are altered in these treatment profiles. Ceruloplasmin (Cp) and catalase are antioxidant enzymes that oxidize Fe^2+^, converting it to Fe^3+^. Both of these enzymes are downregulated with *t*-BG treatment, and Cp is downregulated upon NGF treatment. Heme oxygenase-1 (Hmox1) converts heme into biliverdin, releasing free ferrous iron and carbon monoxide. Iron-Sulfur Cluster Assembly Factor 1 and 2 (Nubp1 and Nubp2 respectively) play an integral role in the machinery of cytosolic iron-sulfur cluster assembly [29]. All three enzymes (Hmox1, Nubp1, and Nubp2) were downregulated in NGF treatment, but not changed in a statistically significant manner in *t*-BG treatment. In contrast, a non-heme iron-containing enzyme, Alpha-ketoglutarate-dependent dioxygenase (Fto) was downregulated in only *t*-BG treatment.

Mitochondrial iron-binding proteins are observed to be downregulated in both treatment regimes, including NADH: ubiquinone oxidoreductase core subunit S1 (Ndufs1), NADH: ubiquinone oxidoreductase core subunit S2 (Ndufs2) (also significantly downregulated with NGF treatment), NADH: ubiquinone oxidoreductase core subunit S 3 (Ndufs3), NADH: ubiquinone oxidoreductase core subunit A9 (Ndufa9) and NADH: ubiquinone oxidoreductase core subunit V1 (Ndufv1). These proteins are core subunits of mitochondrial complex I and are responsible for maintaining proper redox balance in the cell and proper oxidative respiration (See Figure 4) [30].

The protein level changes that are significant in *t*-BG treatment as compared to NGF treatment (the differences between the groups) are shown in Figure 5. Of note, alpha ketoglutarate dependent dioxygenase (Fto) has < 2 fold change. However, significant differences in Ferritin (light chain, ftl1), Heme oxygenase 1 (Hmox1), BolA Family Member 2 (Bola2), NUBP Iron-Sulfur Cluster AssemblyFactor 1 (Nubp1), Ubiquinone Oxidoreductase Core Subunit S1 (Ndufs1), and NADH dehydrogenase [ubiquinone] flavoprotein 1 (Ndufv1) all suggest that *t*-BG alters iron-binding proteins (and possibly iron homeostasis) to a greater extent than NGF alone.

**Figure 5:**
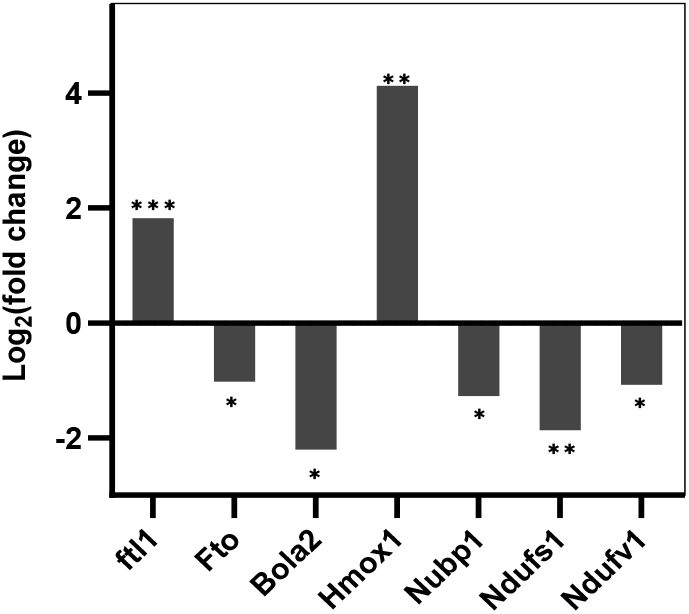
Significant protein expression changes *t*-BG compared to NGF treatment *p<0.05, **p<0.01.

### Western Blot Results

Given these proteomic results, we sought to look for the impacts of protein expression at a shorter time point (2 hours) to compare to our 48-hour data, and to expand the investigation to include both the study of the active enantiomer of *t*-BG alone [(–)-*t*-BG], and a combined *t*-BG/NGF treatment. The latter has previously been shown to cause augmented PC-12 differentiation as compared to NGF or *t*-BG alone [14]. Further, Western blots would allow for identification of subtle (<2 fold change) alterations in protein expression levels with increased statistical confidence.

We first examined one type of driver kinase (Akt, Erk and Pkc) for each of three key pathways triggered by NGF/Trk signal transduction. At 48 hours, Akt and Erk Western blots were consistent with global proteomic results; no expression level changes were observed for control, NGF treatment alone or racemic *t*-BG (See Figure 6a and 6b). Akt expression was increased only with (–)*t*-BG, while Erk expression was decreased with combined *t*-BG and NGF treatment. At 2 hours, no treatment caused statistically significant changes in Akt or Erk expression levels.

**Figure 6:**
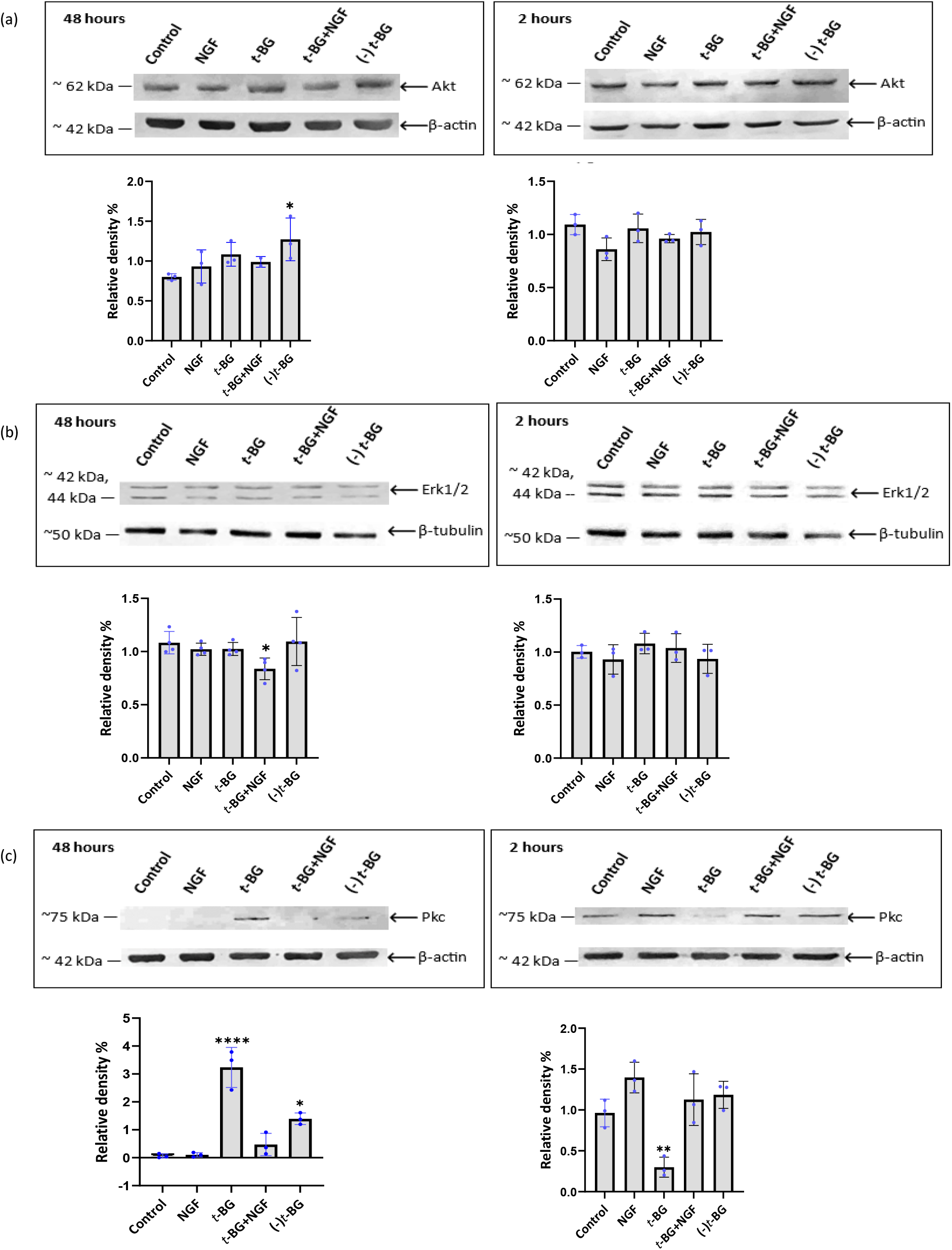
Western blot analysis and quantification. β-actin or β-tubulin was used as a loading reference. Each treatment contains 0.6% DMSO: Control - no additive, NGF (10 ng/mL), *t*-BG (30 μM), *t*-BG (30 μM) + NGF (10 ng/mL), (–)*t*-BG (15 μM). ^*^p<0.05, ^**^p<0.01, ^***^p<0.001, ^****^p<0.0001 (a) Protein loading: Akt (20 μg) (b) Erk1/2 (20 μg) (c) Pkc (30 μg)

Pkc expression levels were the most altered in this group, which might be expected with the observed expression level increase in Pkcα with NGF treatment in the global proteomics analysis. Here, with a pan-Pkc antibody, Western blots reveal effects quite distinct from that result (See Figure 6(c)). At 48 hours, NGF-induced changes in Pkc levels were negligible, though some expression level changes may be lost due to the weak bands observed for the control and the NGF bands. However, *t*-BG, both racemic and enantioenriched, showed significant increases in Pkc expression levels at 48 hours. At 2 hours, curiously, (±) *t*-BG resulted in decreased Pkc expression, suggesting that the (+)*t*-BG enantiomer may have some effect on Pkc levels. The origin of the inconsistencies between our global proteomics and Western blot results is unclear, but it should be noted that the Western blot data is measuring all Pkc isoforms.

We also investigated downstream transcription factors which are activated as a result of Akt, Erk, and Pkc kinase activity in the NGF/Trk pathway. At 48 hours, NGF treatment did not affect NFκB levels, which is downstream of Akt and Pkc [31,32], but *t*-BG treatment (racemic and enantiopure) caused a modest increase (See Figure 7(a)). At 2 hours, a modest decrease of NFκB was observed for both racemic and enantiopure t-BG.

**Figure 7:**
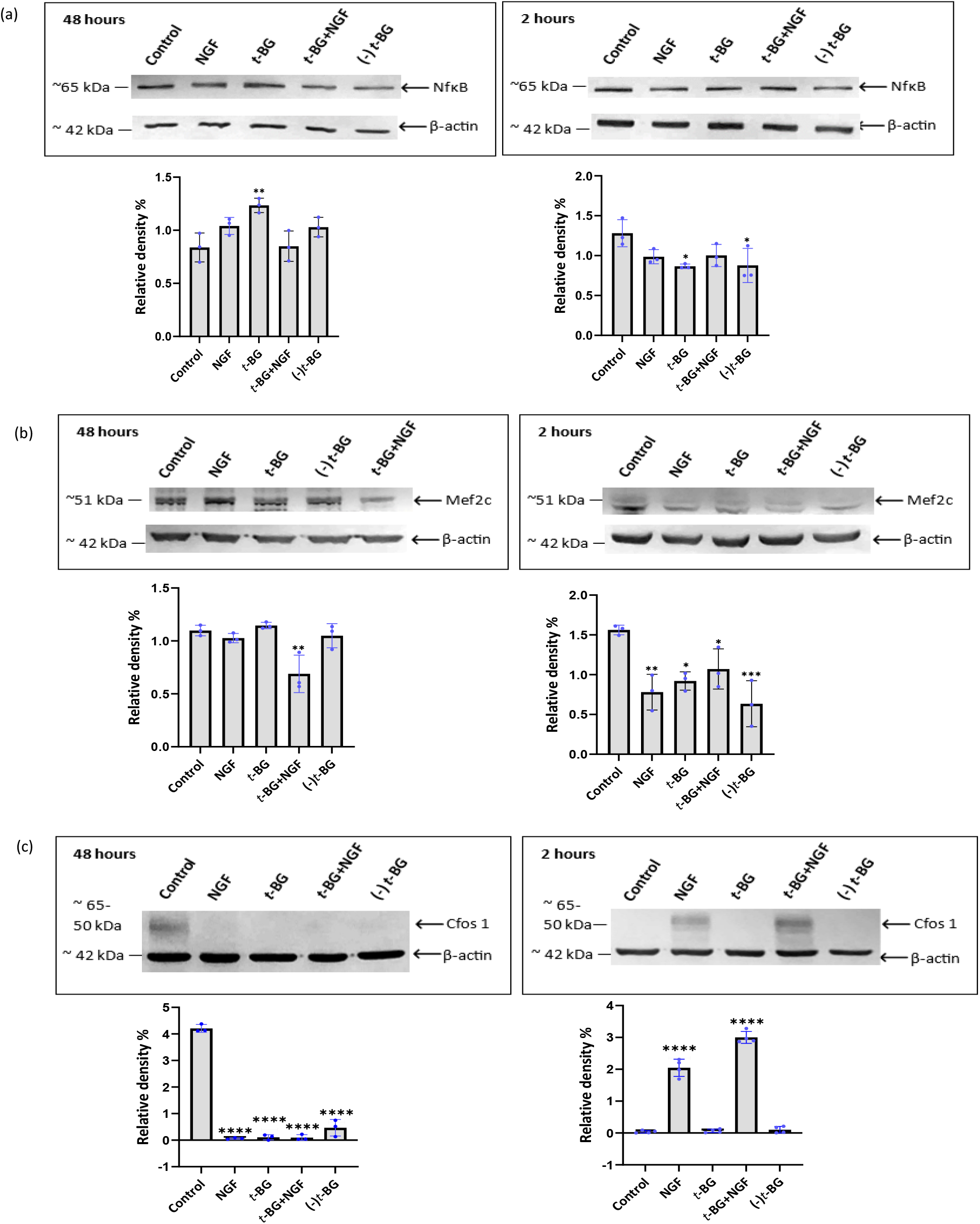
Western blot analysis and quantification. β-actin or β-tubulin was used as a loading reference. Each treatment contains 0.6% DMSO: Control - no additive, NGF (10 ng/mL), *t*-BG (30 μM), *t*-BG (30 μM) + NGF (10 ng/mL), (–)*t*-BG (15 μM). ^*^p<0.05, ^**^p<0.01, ^***^p<0.001, ^****^p<0.0001 (a) Protein loading: NFκB (7.5 μg), (b) Mef2C (30 μg) (c) Cfos 1 (20 μg)

In NGF/Trk signaling, once Erk is translocated into the nucleus, it reacts with nuclear targets such as E twenty-six Like Tyrosine Kinase 1 (Elk1) and myocyte enhancer factor 2 (Mef2). Mef2 members consists of Mef2A, Mef2B, Mef2C, and Mef2D. Since Mef2C is highly abundant in the central nervous system compared to other members, we looked at expression levels of Mef2C (See Figure 7(b)). According to our Western blot data, the combined *t*-BG/NGF treatment slightly reduces Mef2C protein expression levels after 48 hours, which is consistent with a slightly reduced Erk1/2 expression observed for the same treatment profile at 48 hours. Although it is generally believed that Erk5 is primarily involved in inducing Mef2C protein expression, Ling and coworkers have reported that the inhibition of Erk1/2 decreased Mef2C expression levels in embryonic stem cells [33]. Interestingly, Mef2C protein expression levels with NGF, *t*-BG, *t*-BG + NGF and (–)*t*-BG treatments were all found to be lowered after 2 hours, despite no statistical change in Erk1/2 expression for all treatment regimens at this time.

Since Erk signaling is ascribed to be the primary driver of NGF-induced differentiation, the consistent downregulation of Mef2C is surprising. However, it should be noted that overexpression of Mef2C in hippocampal neural stem cells was shown to inhibit both neurogenesis and differentiation [34]. Therefore, Mef2C regulation may be complex in neurotrophic response contexts.

The final transcription factor investigated is c-Fos 1, which is a component of AP-1, and is downstream of Pkc in the NGF/Trk cascade (See Figure 7(c)). Here, Western blots detect both c-Fos 1 (∼60 kDa) and sumoylated c-Fos 1 (∼95 kDa, See Supplemental Info) [35]. Sumoylated c-Fos 1 levels remain unchanged for all treatment profiles at 2 hours and 48 hours. However, all treatments cause dramatic decreases (∼ 4.5 fold) in c-Fos 1 at 48 hours. c-Fos 1 is also strongly increased at 2 hours with both NGF (∼ 2 fold) and NGF + *t*-BG (∼ 3 fold) treatments, and unaffected by *t*-BG alone.

The distinct increase in c-Fos 1 at 2 hours is consistent with increased Pkc activity induced by NGF; that *t*-BG does not elicit a similar effect suggests that it may not intercept the same Pkc-activating pathways as NGF.

Next, we looked at the iron-binding/storage proteins involved in iron homeostasis and oxidative stress, given the trends observed from global proteomics data. Ferritin is a central protein in iron homeostasis because it binds free cytosolic iron (often referred to as the labile iron pool) and stores it for later use. Ferritin light chain (ftl) expression was dramatically increased at 48 hours (up to 3.3-fold) with *t*-BG and slightly decreased with NGF (See Figure 8(a)). This is consistent with global proteomics results in the effect, although not the magnitude, of *t*-BG on ftl1 expression level change. At 2 hours, no change in ftl expression was observed for all treatments.

**Figure 8:**
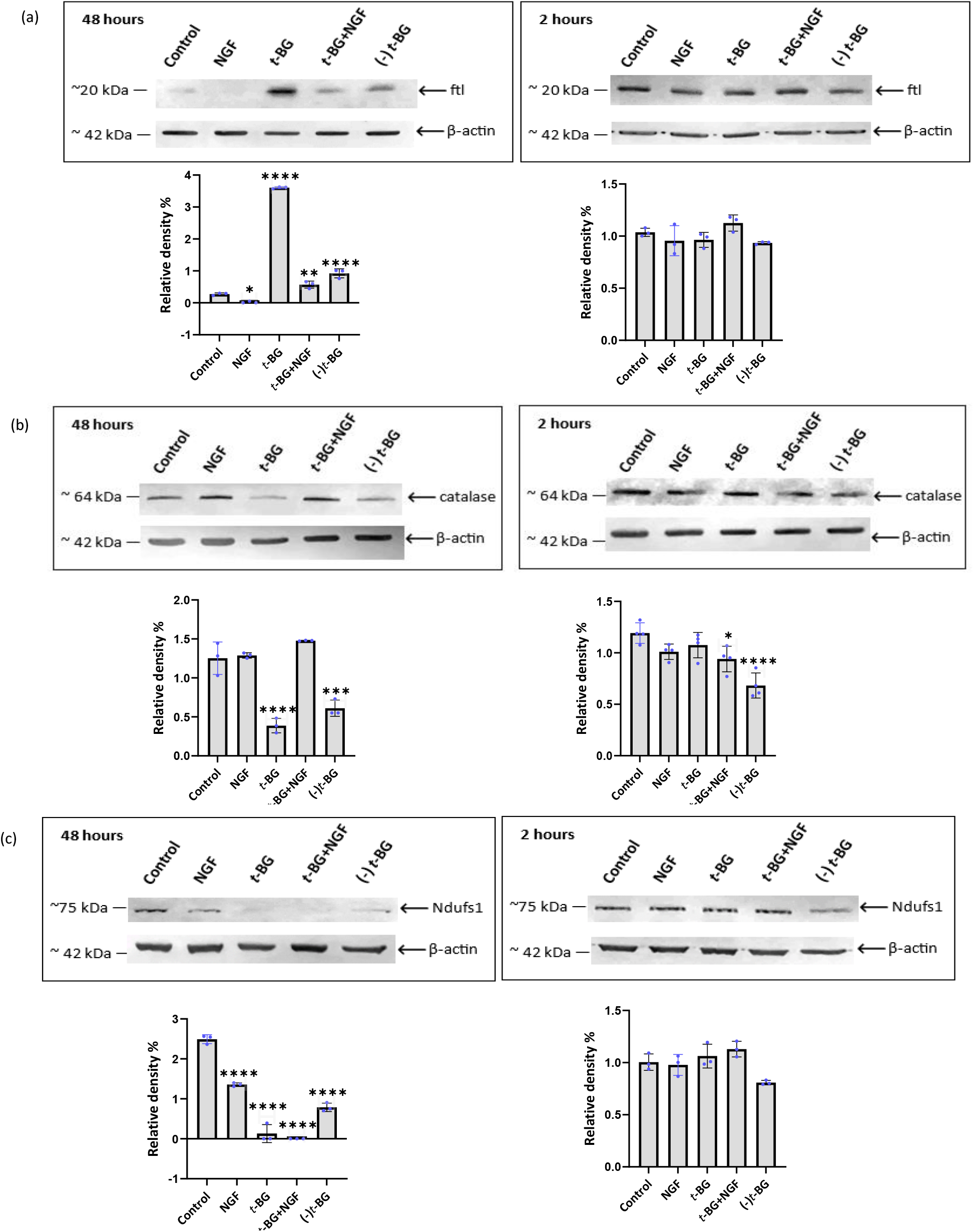
Western blot analysis and quantification. β-actin or β-tubulin was used as a loading reference. Each treatment contains 0.6% DMSO: Control - no additive, NGF (10 ng/mL), *t*-BG (30 μM), *t*-BG (30 μM) + NGF (10 ng/mL), (–) *t*-BG (15 μM). *p<0.05, **p<0.01, ***p<0.001, ****p<0.0001 (A). Protein loading: (a) Ferritin (10 μg) (b) Catalase (10 μg) (c) Ndufs1 (30 μg)

In contrast, catalase protein expression levels were downregulated at 48 hours with all *t*-BG treatments, as well as NGF treatment. This is also consistent with global proteomics results, which demonstrated a 1.5 fold decrease in catalase expression with *t*-BG treatment (See Figure 8(b)). Decreases at 2 hours are observed for enantiopure *t*-BG only and for combined *t*-BG/NGF treatments.

Our Western blot data shows that Ndufs1 protein, part of mitochondrial complex I, was significantly downregulated in all the treatments compared to the control at 48 hours, with no statistically significant change with all treatments at 2 hours (See Figure 8(c). The significant downregulation of Ndufs1 expression with *t*-BG treatment is consistent with global proteomic data, which shows a 2-fold decrease in Ndufs1 expression. The decrease in expression level from NGF treatment is more modest, also consistent with the global proteomics results.

### Intracellular iron measurement

To connect iron-binding protein level changes to intracellular iron levels, we measured intracellular ferrous iron using a FerroOrange fluorescent assay. The assay results indicated that Fe^2+^ levels decreased markedly in all *t*-BG treatment groups (Figure 9). The degree of change may also be correlated with the relative increase in ftl protein, as measured by Western blot and shown in Figure 8a.

**Figure 9:**
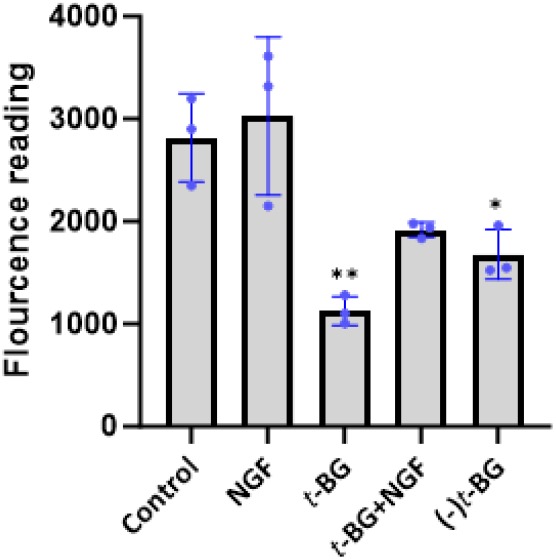
Cellular Fe^2+^ measurements using FerroOrange. A higher fluorescence reading is indicative of a higher intracellular concentration of Fe^2+^.

### Discussion and Conclusion

As expected, most of our Western blot analyses for 48-hour treatment profiles match the general profile of the global proteomics data, including Akt, Erk, ferritin, Ndufs1, and catalase. Notably, Western blot analyses identified subtle (<2 fold) changes in ferritin and Ndufs1 with NGF treatment at 48 hours in Western blot quantification.

While many pro-growth factors are upregulated in NGF treatment samples (Alcam, Serpinb1A, β-synuclein, c-Fos 1, Pkcα), in *t*-BG treated samples, factors which negatively regulate these pro-growth responses were upregulated (SHP-1, Trim32), and c-Fos 1, a major indicator of pro-growth responses, was notably not affected at 2 hours of treatment. These are important distinctions between the cell effects and signaling responses to *t*-BG as compared to NGF; we see no obvious evidence that *t*-BG intercepts NGF/TrkA signaling.

For Pkc expression levels, Western blots differed distinctly from global proteomics results. All *t*-BG treatments caused marked increases in Pkc at 48 hours by Western blot, while global proteomics showed no increases. These Western blot measurements were conducted with a pan-Pkc antibody; it has been previously reported that PC-12 cells contain α, δ, ε and ζ isoforms out of 11 possible Pkc isoforms, including conventional (α, βI, βII, γ), novel (δ, ε, η, θ, μ), and atypical (ζ, ι).[36,37] Generally, Pkc activity is associated with synaptic plasticity in neuronal cells [38, 39]. Therefore, the increase in Pkc expression may be related to some of the neuritogenic effects of *t*-BG. While NGF is known to activate Pkc, no increase in expression level was observed in our Western blot assays, despite the increase in Pkcα observed in global proteomic analysis.

Further notable protein expression changes identified through Western blots were those of transcription factors Mef2C and c-Fos 1. In particular, c-Fos is categorized as an immediate and early expression gene that activates rapidly in response to stimulus [40]. The remarkable transient upregulation of c-Fos 1 protein expression levels with NGF and combined NGF + *t*-BG treatments at 2 hours, and downregulation at 48 hours with all treatments, is consistent with responses reported by Milbrandt and colleagues. They have shown that NGF rapidly increased *c-fos* mRNA levels in PC-12 cells at 15-45 mins, followed by a rapid decrease after 120 mins [41], due to self-repression of mRNA expression by c-Fos protein [40]. That this initial increase at 2 hours is not observed in *t*-BG treatments in the absence of NGF suggests that *t*-BG likely does not intercept/activate this part of the NGF/TrkA signaling pathway.

Less anticipated at the outset of our study was the impact on both NGF and *t*-BG on iron-binding and iron storage proteins. It has been previously reported that NGF promotes hippocampal and cortical neuronal survival against iron catalyzed oxidative damage [42], and that NGF increases the uptake of iron in PC-12 cells [43], but otherwise there is little known about the connection between neurotrophic agents and iron homeostasis. According to our global proteomics analysis, twelve different iron binding/storage proteins were impacted significantly by NGF and/or *t*-BG treatment, suggesting these treatments may have an impact on iron regulation and utilization in the cell. Iron homeostasis is controlled through iron uptake, compartmentalization, storage, and excretion [44]. Iron is transported into cells via transferrin-bound and non-transferrin bound iron uptake pathways, which are summarized in Figure 10 [43].

**Figure 10:**
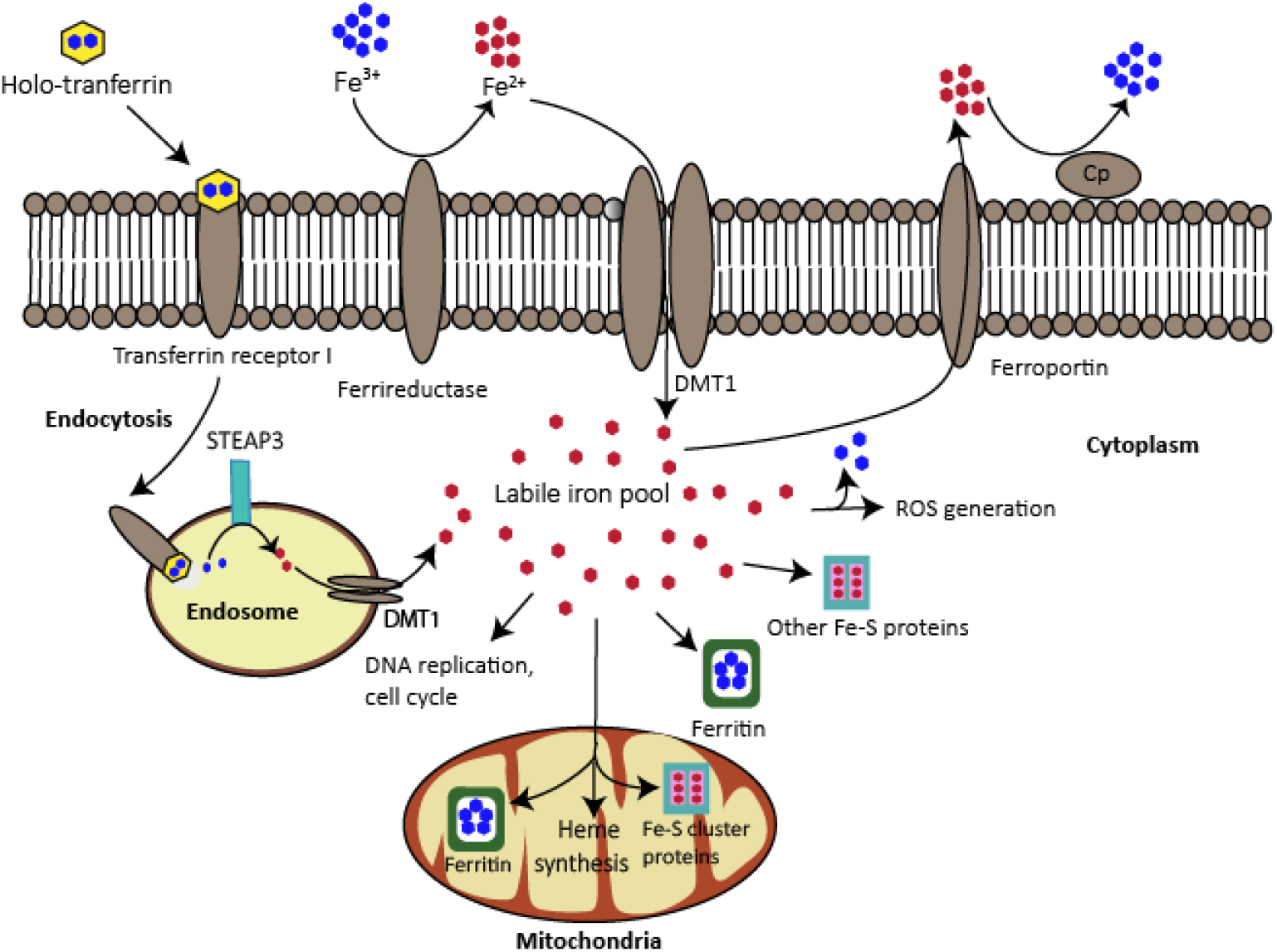
Cellular iron homeostasis. In the transferrin bound pathway, holo-transferrin (Fe^3+-^bound transferrin) binds to transferrin receptor 1 [43]. This complex internalizes into the cell via endocytosis. The acidic environment in the endosome releases iron from transferrin and six-transmembrane epithelial antigen of the prostate 3 (Steap3) metalloreductase reduces iron to give Fe^2+^ [45]. The subsequent release of Fe^2+^ into the cytoplasm (to become part of the labile iron pool) is facilitated by divalent metal transporter 1 (DMT1) [46]. In the non-transferrin-bound iron uptake pathway, Fe^3+^ outside the cell is reduced to Fe^2+^ by ferrireductase. DMT1 then transports Fe^2+^into the cytosol by DMT1. Fe is excreted from the cell as Fe^2+^ by ferroportin with coordination of ceruloplasmin.

Fe^2+^ is important for multiple cellular processes, including DNA replication, oxygen transport, ATP synthesis/respiration, and redox balance [47]. Free Fe^2+^ in the cytosol also generates reactive oxygen species (ROS) via the Fenton reaction. These ROSs cause oxidative stress when iron levels are high, most notably by damaging lipids in the cell membrane, and can ultimately lead to cell death (ferroptosis). Excess Fe^2+^ is stored as Fe^3+^ bound to ferritin in both the cytoplasm and mitochondria, effectively regulating the levels of free (labile) iron.

Mwanjewe and co-workers have reported that NGF-induces increased iron uptake in PC-12 cells predominately through a non-transferrin bound pathway and also increases nitric oxide synthase [43]. Nitric oxide represses the translation of ferritin mRNA by activating iron regulatory factor proteins that play a central role in iron metabolism [48]. This may be why we observe a modest downregulation of ferritin expression levels with NGF treatment at 48 hours by Western blot.

In contrast, we see significant increases in ferritin light chain expression levels from *t*-BG treatment in both global proteomics and Western blot quantification, which could impact free cytosolic iron levels. To gain insight into this connection, we evaluated ferrous iron levels and observed decreased Fe^2+^ levels in all *t*-BG treatment groups.

The downregulation of mitochondrial enzymes in response to *t*-BG observed in both global proteomics results and from Western blot assays (Figure 4, Figure 8: Western blots for catalase, Ndufs1) could arise due to a decreased need for reductive activity. If there is a lower concentration of iron in the cytosol due to an increase in ferritin, then there may also be lower oxidative pressure in the cell. Alternatively, the sequestering of free cytosolic iron by ferritin may result in insufficient free iron concentrations in the mitochondria. If true, the biosynthesis of many oxidoreductases and related iron-containing enzymes would be hindered by the lack of available mitochondrial iron.

In either case, the known neurotrophic impact of *t*-BG coupled with an impact on iron homeostasis is significant when considering current understanding of many neurodegenerative diseases. For various protein aggregates in neurodegenerative contexts (Aβ, Lewy bodies/α-synuclein, tau tangles) high iron concentrations are co-localized [8], indicating binding and association with these aggregates. In APP mice and in SH-SY5Y cells, increased iron concentrations result in increased Aβ [49,50], postulated to occur through a conformational change of amyloid protein caused by Fe^3+^ along with an iron-triggered increase in β-secretase, which is implicated in Aβ production [51, 52]. Both Fe^2+^ and Fe^3+^ have been shown to accelerate the aggregation [53], hyperphosphorylation [54], and nitration of tau [55], and these tau tangles have also been shown to promote Fe^3+^ reduction to Fe^2+,^ resulting in increased oxidative stress in the cell [11]. There is also growing evidence of a connection between ferroptotic pathways and neurodegenerative disease [12, 56]. As such, treatments which alter iron homeostasis levels in neuronal cells may have critical roles in slowing the development and/or progression of neurodegenerative disease.

Given that neurodegenerative mechanisms appear intimately connected to iron levels in brain tissues, perhaps it should be unsurprising to observe a connection between iron-binding proteins and neurotrophic responses. Our proteomic results highlight this relationship along with new TrkA-related cell responses to both NGF and *t*-BG. Further investigation into the mechanism of *t*-BG will be informed by these insights and may ultimately establish important connections between neurotrophic activity and iron management in neuronal cells.

## MATERIAL AND METHODS

### Organic synthesis procedure

Step 1 and 2 of BG synthesis was conducted as reported in Khyati *et al* (2022) (Scheme 1) [4].

**Scheme 1:**
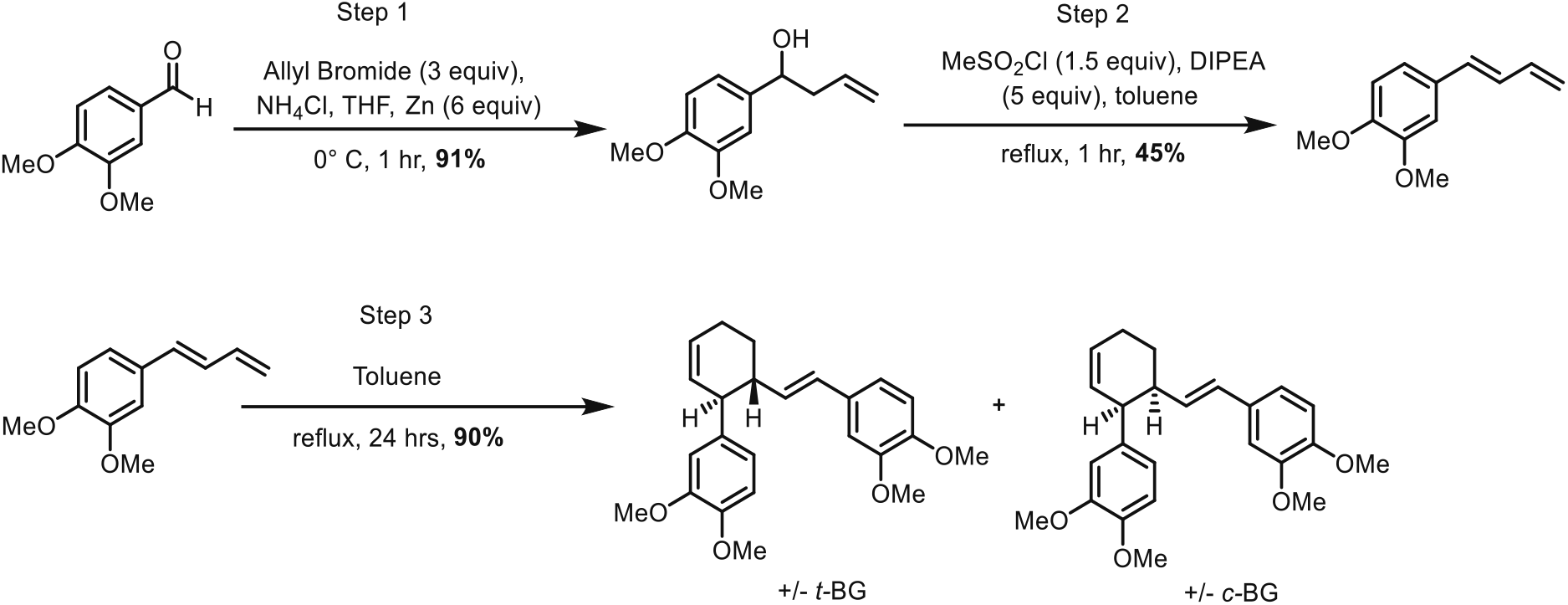
Banglene synthesis

Step 3 was slightly modified as follows: in a flame dried 5 mL RBF, the diene (0.90 g. 4.73 mmol) was dissolved in dry toluene (2 mL). The reaction was heated to reflux for 24 hours under N_2_. The crude mixture was concentrated in vacuo, and the resulting residue was purified by flash column chromatography followed by preparative HPLC.

The enantiomers of ±*t*-BG were separated commercially by Lotus Separations. See Gohil, K.; Kazmi, M. Z. H.; Williams, F. J. *Org. Biomol. Chem*., **2022**, *20*, 2187 for details regarding enantiopurity measurements and optical rotation for these separated enantiomer samples.[4] See supplemental information for further information on the synthesis, isolation, and characterization of *t*-BG.

### PC-12 cell culture

PC-12 cell line (CRL-1721) was obtained from ATCC. PC-12 cells were maintained in growth medium containing Dulbecco’s modified Eagle’s medium (DMEM) (GibcoTM LS11965092) for global proteomics analysis and Roswell Park Memorial Institute (RPMI) (Gibco™ 11875093) medium for Western blot analysis, horse serum (HS) (Gibco™ 16050122), fetal bovine serum (FBS) (Gibco™ 16000069) and Pen-Strep (Gibco™ 15140122). Passage 4 at 70% confluency were taken for protein extractions. Hematocytometer was used to count the cells stained in trypan blue dye.

### Global proteomics profiling

Passage 4 PC-12 cells (at 70% confluency) were seeded in the collagen-IV coated T-175 flasks (Corning® BioCoat® 354528) at a density of 2×10^4^ cells/cm^2^ and cultured in DMEM medium containing 5% HS, 5% FBS and 1% Pen-Strep for 24 hours, then the medium was changed to DMEM containing 2% HS, 1% FBS and a given treatment was added-30 μM (±) t-BG or NGF (10 ng/mL) with 0.6% DMSO or 0.6% DMSO. 4 flasks of cells were grown per treatment condition. The cells were cultured for a further 48 hours, following which the cell pellets were collected and flash frozen in liquid nitrogen. Cell pellets were lysed in 4% SDS 100mM tris pH 6.8 and total protein concentration was estimated with BCA assay. Approximately 20 micrograms of total protein was reduced and alkylated with dithiothreitol and iodoacetamide and loaded onto 10% SDS-PAGE [57]. Each sample’s lane was separated into 5 fractionations and the proteins were digested out of the gels using MS-grade trypsin [58]. Resulting peptides were cleaned with C-18 STop And Go Extraction (STAGE) tips using 40% (v/v) acetonitrile in 0.1% (v/v) formic acid as the elution buffer [59]. Peptide concentration was estimated with NanoDrop One (Thermo Scientific) to load equal amounts of total peptides on Impact II Qtof (Bruker Daltonics) coupled to easy nLC 1200 (Thermo Scientific) using a in-house constructed column set up consisting a fritted 2-cm-long trap column made with 100-μm-inner diameter fused silica and 5 μm Aqua C-18 beads (Phenomenex, cat# 04A-4331) packing material, and a 30-50 cm-long analytical column with an integrated spray tip (6-8 μm-diameter opening, pulled on a P-2000 laser puller from Sutter Instruments) made with 75-μm-inner diameter fused silica and 1.9 μm-diameter Reprosil-Pur C-18-AQ beads (Dr. Maisch, cat# r119.aq.0003) packing material. The column was heated to 50°C with another in-house constructed column heater and was used for 60 min sample separation using the acquisition settings [60]. Four biological replicates of the treatment sets were processed, and each sample was analyzed twice for duplicate technical replicates. The samples were randomized before injection.

The resulting data were searched on MaxQuant version 1.6.7.0 using Uniprot’s rat proteome and common contaminants [61]. Label-free quantitation, and match-between-run options were enabled using fixed modification of carbamidomethylation of cysteines, and variable modifications of oxidation of methionines and acetylation of protein N-termini. Specific proteolytic cleavages after arginine and lysine with up to 2 missed cleavages were set. The data were filtered for 1% false discovery at protein, peptide and PSM levels using revert decoy search mode. The mass spectrometry proteomics data have been deposited to the ProteomeXchange Consortium via the PRIDE partner repository with the dataset identifier PXD053648 [62]. The proteomic data was analyzed used Perseus software platform [63]. Volcano plots were made using EnhancedVolcano [64].

### Proteomic analysis

Perseus software was used to analyze the proteomic data. Label-free quantification (LFQ) intensity values from two technical replicates (e.g., DMSO 1a and 1b) for each biological replicate (DMSO 1) were combined. The combining process followed this approach: if both technical replicates had intensity values, the values were averaged. However, if only one technical replicate had a reported intensity value, that value was used. This data was then analyzed using Perseus software. First, annotations were added for context. Next, proteins identified only by site, contaminants, and those identified in reverse were excluded for better data quality. Finally, the data underwent a Log_2_ transformation and a 2-sample Student’s t-test was performed to compare two groups. The Benjamini-Hochberg test was employed to calculate the false discovery rate (FDR) and account for multiple testing.

### Protein extraction for Western blots

#### Collagen coating

225 cm^2^ Corning cell culture flasks (Sigma CLS431080) were coated with 6 μg/cm^2^ of collagen from human placenta Bornstein and traub type IV (Sigma C5533) in hanks’ balanced salt solution ((-) Ca^2^+, (-) Mg^2^+, no phenol) (HBSS)(Gibco™14180046). The flasks were stored at 2-8 °C overnight. Then, excess collagen was removed and rinsed with autoclaved distilled water (20 mL).

#### Protein extraction procedure

Passage 4 PC-12 cells (at 70% confluency) were seeded in the collagen-IV coated T-225 flasks in RPMI medium containing 10% HS, 5% FBS and 1% Pen-Strep for 24 hours 37 °C. Then, growth media were removed and differentiation media containing Reduced Serum Medium Opti-MEM™ (Gibco™ 31985070), 1% HI and 0.5% FBS) were added with treatments. The final concentration of the DMSO in each treatment was 0.6%. The flasks were then incubated for treatmnet hours at 37 °C. Then, the cells were scraped into pre-weighed falcon tubes (50 mL). The cells-filled pre-weighed falcon tubes -were centrifuged at 4000 rpm for 15 mins. Then, the supernatant was discarded, followed by rinsing the cells with 5 mL of cold phosphate buffered saline (PBS) (Gibco™10010023). The cells were centrifuged at 4000 rpm for 10 mins at 4 °C and the supernatant was discarded. The rinsing step was repeated. 1 mL of RIPA lysis buffer (sigma-Aldrich R0278) containing cOmplete™, mini EDTA-free protease inhibitor cocktail (Roche 11836170001) and phosphatase inhibitor – phosSTOP (Sigma Aldrich PHOSS-RO) was added to 42 mg of cell pellet. Cells were homogenized for 30 mins. Then, the cell suspension was centrifuged at 14000 rpm for 15 mins at 0 °C. The supernatants were aliquoted and stored at -78 °C. The concentration of the protein samples was calculated by using DC protein assay kit II (Biorad 500-0112).

### Sodium dodecyl sulfate – polyacrylamide electrophoresis (SDS-PAGE) procedure

#### Reagents

Resolving gel buffer (1.5 M tris-HCl (pH = 8.8), 0.4% SDS), stacking gel buffer (0.5 M tris-HCl (pH = 6.8), 0.4% SDS), running buffer: 10x Tris/Glycine/SDS (biorad 1610732), 4x Laemmli Sample Buffer (biorad 1610747), β-mercaptoethanol, N,N,N’,N’-tetramethylethylenediamine (TEMED) (sigma T9281), Precision Plus Protein™ All Blue Prestained Protein Standards (biorad 1610373), ammonium persulfate (APS), 30% Acrylamide/Bis Solution, 29:1 (biorad 1610156), and isopropanol were used.

#### Procedure

Glass plates were assembled in the casting stand. The SDS-PAGE gel was made using resolving gel mixture (3.635 mL distilled water, 2.185 mL resolving gel buffer, 2.925 mL 30% Acrylamide/Bis Solution, 50 μL APS and 7.5 μL TEMED) and stacking gel mixture (2.925 mL distilled water, 1.25 mL stacking gel buffer, 0.825 mL 30% Acrylamide/Bis Solution, 50 μL APS and 7.5 μL TEMED). The gel was placed in the buffer chamber containing running buffer (Tris/Glycine/SDS). Different protein treatment samples and precision plus protein™ all blue prestained protein ladder were loaded into each well after denaturing with 4x Laemmli Sample Buffer and β-mercaptoethanol. The electrophoresis was run for 1:10-1:45 hours at 200 V.

### Western blot procedure

#### Reagents and antibodies

×10 TBST (24.23 g Tris base, pH 7.4, 87.66 g NaCl, 10 mL tween 20 in 1 L of water), ×10 transfer buffer without HPLC grade methanol (30.3 tris base, 144 g of glycine in 1 L of water), ×1 transfer buffer with HPLC grade methanol (100 mL of 10 transfer buffer without HPLC grade methanol, 800 mL of distilled water, 200 mL methanol), nonfat dry milk, BSA, PKC Pan Polyclonal Antibody (Invitrogen™ PA5120593), Anti-NF-kB p65 antibody (ab16502), Recombinant Anti-c-fos 1 antibody [EPR20769] (ab214672), Anti-Ferritin Light Chain antibody (ab69090), Ndufs1 Rabbit, Polyclonal, Novus Biologicals™ (NBP15652020), Anti-Catalase antibody - Peroxisome Marker (ab52477), ERK1/ERK2 Rabbit anti-Human, Mouse, Rat, Polyclonal (Invitrogen™ 44-654G), AKT Pan Rabbit anti-Human, Mouse, Rat, Polyclonal, (Invitrogen™ PA5-77855), beta Actin Polyclonal Antibody (Invitrogen™ PA116889), beta Tubulin Polyclonal Antibody, (Invitrogen™ PA516863), Phospho-AKT1 (Thr308) Recombinant Polyclonal Antibody (B18HCLC) (Invitrogen™ 710122), Anti-MEF2C antibody [EPR19089-202] - ChIP Grade (ab211493), Goat Anti-Rabbit IgG (H+L), HRP, Secondary Antibody (Promega PRW4011), and 1-Step™ Ultra TMB-Blotting Solution (Invitrogen™ 37574) were used.

#### Procedure

After SDS-PAGE, the gel was transferred into 5 mL of ×1 transfer buffer for 2 mins. Transfer buffer-soaked gel was then stacked with a nitrocellulose membrane into a gel holder cassette. The cassette was placed in the ×1 transfer buffer filled chamber. The voltage was applied at 50 V for 2 hours. Then, the nitrocellulose membrane was transferred and rinsed with 3×5 mL of TBST in a 500 mL beaker. The membrane was blocked using BSA/skim milk for 2 hours at room temperature and incubated with the primary antibody at 4 °C overnight. Then, the membrane was rinsed with TBST (5×5mL) at room temperature, followed by treating with secondary antibody for 1 hour. Finally, the membrane was treated with 1-Step™ Ultra TMB-Blotting Solution to expose the bands after rinsing with TBST (5×5mL) (See SI for images of all Western blots). Fiji Image J was used to quantify the western blots [65]. Graphpad prism 10 was used to plot the graphs for western blots.

### Intracellular iron measurement

#### Collagen coating

SPL life sciences cell culture dishes (cat# 20060) were coated with 6 μg/cm^2^ of collagen from human placenta Bornstein and traub type IV (Sigma C5533) in hanks’ balanced salt solution ((-) Ca^2^+, (-) Mg^2^+, no phenol) (HBSS)(Gibco™14180046). The dishes were stored at 2-8 °C overnight. Then, excess collagen was removed and rinsed with autoclaved distilled water (6 mL).

#### Assay procedure

Passage 4 PC-12 cells (at 70% confluency) were seeded in the collagen-IV coated cell culture dishes in 5 mL of RPMI medium containing 10% HS, 5% FBS and 1% Pen-Strep for 24 hours at 37 °C. Then, growth media were removed and 5 mL of differentiation media containing Reduced Serum Medium Opti-MEM™ (Gibco™ 31985070), 1% HI and 0.5% FBS) were added with treatments. The treatments were done in three replicas. The final concentration of the DMSO in each treatment was 0.6%. The dishes were then incubated for 48 hours at 37 °C. Then, the cells were scraped into 15 mL falcon tubes. The cells-filled falcon tubes were centrifuged at 4000 rpm for 10 mins. Then, the supernatant was discarded, followed by rinsing the cells with 15 mL of HBSS. The cells were centrifuged at 4000 rpm for 10 mins and the supernatant was discarded. The rinsing step was repeated. 0.2 *μ*L of 1 mmol/L ferroOrange dye (Dojindo laboratories item code: F374-10) was added each falcon tube and incubated the falcon tubes at 37 °C for 30 mins. The fluorescent reading was taken at 543 nm/580 nm.

## Author Contributions

The manuscript was written by Piyumi Wijesiri Gunawardana and Florence J Williams, with the aid of feedback from all other authors. For global proteomic analyses, Dr. Khyati Gohil prepared the biological samples, while Kyung-Mee Moon prepared the technical replicates and performed the mass spectrometric analyses. Prof. Leonard Foster oversaw the implementation of global proteomic profiling and the statistical analysis of the data. Piyumi Wijesiri Gunawardana performed all Western blots. All authors have given approval to the final version of the manuscript.

## Data availability

The Supporting Information is available free of charge on BioRxiv.

## ACKNOWLEDGEMENT

This work was supported by the Canada Research Coordinating Committee NFRFE-2018-00223 and University of Iowa. The mass spectrometry infrastructure was supported by the Canada Foundation for Innovation and the BC Knowledge Development Fund. Proteomics operations were supported by Genome Canada/Genome BC project (264PRO)

## REFERNECES

(1) Pardo-Moreno, T.; González-Acedo, A.; Rivas-Domínguez, A.; García-Morales, V.; García-Cozar, F. J.; Ramos-Rodríguez, J. J.; Melguizo-Rodríguez, L., Therapeutic Approach to Alzheimer’s Disease: Current Treatments and New Perspectives. Pharmaceutics 2022, 14 (6), 1117.

(2) Tee, A. R., The Target of Rapamycin and Mechanisms of Cell Growth. International Journal of Molecular Sciences 2018, 19 (3), 880.

(3) Chu, J.; Suh, D. H.; Lee, G.; Han, A. R.; Chae, S. W.; Lee, H. J.; Seo, E. K.; Lim, H. J., Synthesis and biological activity of optically active phenylbutenoid dimers. J Nat Prod 2011, 74 (8), 1817–21.

(4) Gohil, K.; Kazmi, M. Z. H.; Williams, F. J., Structure-activity relationship and bioactivity studies of neurotrophic trans-banglene. Organic & Biomolecular Chemistry 2022, 20 (11), 2187–2193.

(5) Matsui, N.; Kido, Y.; Okada, H.; Kubo, M.; Nakai, M.; Fukuishi, N.; Fukuyama, Y.; Akagi, M., Phenylbutenoid dimers isolated from Zingiber purpureum exert neurotrophic effects on cultured neurons and enhance hippocampal neurogenesis in olfactory bulbectomized mice. Neurosci Lett 2012, 513 (1), 72–7.

(6) van Bergen, J. M. G.; Li, X.; Hua, J.; Schreiner, S. J.; Steininger, S. C.; Quevenco, F. C.; Wyss, M.; Gietl, A. F.; Treyer, V.; Leh, S. E.; Buck, F.; Nitsch, R. M.; Pruessmann, K. P.; van Zijl, P. C. M.; Hock, C.; Unschuld, P. G., Colocalization of cerebral iron with Amyloid beta in Mild Cognitive Impairment. Scientific Reports 2016, 6 (1), 35514.

(7) Ma, L.; Gholam Azad, M.; Dharmasivam, M.; Richardson, V.; Quinn, R. J.; Feng, Y.; Pountney, D. L.; Tonissen, K. F.; Mellick, G. D.; Yanatori, I.; Richardson, D. R., Parkinson’s disease: Alterations in iron and redox biology as a key to unlock therapeutic strategies. Redox Biology 2021, 41, 101896.

(8) Ndayisaba, A.; Kaindlstorfer, C.; Wenning, G. K., Iron in Neurodegeneration – Cause or Consequence? Frontiers in Neuroscience 2019, 13.

(9) Ayton, S.; Wang, Y.; Diouf, I.; Schneider, J. A.; Brockman, J.; Morris, M. C.; Bush, A. I., Brain iron is associated with accelerated cognitive decline in people with Alzheimer pathology. Molecular Psychiatry 2020, 25 (11), 2932–2941.

(10) Peng, Y.; Chang, X.; Lang, M., Iron Homeostasis Disorder and Alzheimer’s Disease. International Journal of Molecular Sciences 2021, 22 (22), 12442.

(11) Ward, R. J.; Zucca, F. A.; Duyn, J. H.; Crichton, R. R.; Zecca, L., The role of iron in brain ageing and neurodegenerative disorders. The Lancet Neurology 2014, 13 (10), 1045–1060.

(12) Reichert, C. O.; de Freitas, F. A.; Sampaio-Silva, J.; Rokita-Rosa, L.; Barros, P. L.; Levy, D.; Bydlowski, S. P., Ferroptosis Mechanisms Involved in Neurodegenerative Diseases. Int J Mol Sci 2020, 21 (22).

(13) Huang, E. J.; Reichardt, L. F., Trk receptors: roles in neuronal signal transduction. Annu Rev Biochem 2003, 72, 609–42.

(14) Gohil, K.; Kazmi, M. Z. H.; Williams, F. J., Structure-activity relationship and bioactivity studies of neurotrophic trans-banglene. Org Biomol Chem 2022, 20 (11), 2187–2193.

(15) Schrick, C.; Fischer, A.; Srivastava, D. P.; Tronson, N. C.; Penzes, P.; Radulovic, J., N-cadherin regulates cytoskeletally associated IQGAP1/ERK signaling and memory formation. Neuron 2007, 55 (5), 786–98.

(16) Loh, C. Y.; Chai, J. Y.; Tang, T. F.; Wong, W. F.; Sethi, G.; Shanmugam, M. K.; Chong, P. P.; Looi, C. Y., The E-Cadherin and N-Cadherin Switch in Epithelial-to-Mesenchymal Transition: Signaling, Therapeutic Implications, and Challenges. Cells 2019, 8 (10).

(17) Zhang, J.; Shemezis, J. R.; McQuinn, E. R.; Wang, J.; Sverdlov, M.; Chenn, A., AKT activation by N-cadherin regulates beta-catenin signaling and neuronal differentiation during cortical development. Neural Dev 2013, 8, 7.

(18) Wade, A.; Thomas, C.; Kalmar, B.; Terenzio, M.; Garin, J.; Greensmith, L.; Schiavo, G., Activated leukocyte cell adhesion molecule modulates neurotrophin signaling. J Neurochem 2012, 121 (4), 575–86.

(19) Conroy, J. N.; Coulson, E. J., High-affinity TrkA and p75 neurotrophin receptor complexes: A twisted affair. Journal of Biological Chemistry 2022, 298 (3).

(20) Roux, P. P.; Barker, P. A., Neurotrophin signaling through the p75 neurotrophin receptor. Prog Neurobiol 2002, 67 (3), 203–33.

(21) Seaborn, T.; Ravni, A.; Au, R.; Chow, B. K.; Fournier, A.; Wurtz, O.; Vaudry, H.; Eiden, L. E.; Vaudry, D., Induction of serpinb1a by PACAP or NGF is required for PC12 cells survival after serum withdrawal. J Neurochem 2014, 131 (1), 21–32.

(22) Hashimoto, M.; Bar-on, P.; Ho, G.; Takenouchi, T.; Rockenstein, E.; Crews, L.; Masliah, E., β-Synuclein Regulates Akt Activity in Neuronal Cells: A POSSIBLE MECHANISM FOR NEUROPROTECTION IN PARKINSON′S DISEASE*. Journal of Biological Chemistry 2004, 279 (22), 23622–23629.

(23) Black, A. R.; Black, J. D., The complexities of PKCα signaling in cancer. Adv Biol Regul 2021, 80, 100769.

(24) Yasuda, S.; Kai, M.; Imai, S.; Takeishi, K.; Taketomi, A.; Toyota, M.; Kanoh, H.; Sakane, F., Diacylglycerol kinase eta augments C-Raf activity and B-Raf/C-Raf heterodimerization. J Biol Chem 2009, 284 (43), 29559–70.

(25) Marsh, H. N.; Dubreuil, C. I.; Quevedo, C.; Lee, A.; Majdan, M.; Walsh, G. S.; Hausdorff, S.; Said, F. A.; Zoueva, O.; Kozlowski, M.; Siminovitch, K.; Neel, B. G.; Miller, F. D.; Kaplan, D. R., SHP-1 negatively regulates neuronal survival by functioning as a TrkA phosphatase. J Cell Biol 2003, 163 (5), 999–1010.

(26) Du, H. X.; Wang, H.; Ma, X. P.; Chen, H.; Dai, A. B.; Zhu, K. X., Eukaryotic translation initiation factor 2α kinase 2 in pancreatic cancer: An approach towards managing clinical prognosis and molecular immunological characterization. Oncol Lett 2023, 26 (5), 478.

(27) Staudinger, J.; Zhou, J.; Burgess, R.; Elledge, S. J.; Olson, E. N., PICK1: a perinuclear binding protein and substrate for protein kinase C isolated by the yeast two-hybrid system. J Cell Biol 1995, 128 (3), 263–71.

(28) Hillje, A.-L.; Worlitzer, M. M. A.; Palm, T.; Schwamborn, J. C., Neural Stem Cells Maintain Their Stemness through Protein Kinase C ?-Mediated Inhibition of TRIM32. Stem Cells 2011, 29 (9), 1437–1447.

(29) Stehling, O.; Netz, D. J.; Niggemeyer, B.; Rösser, R.; Eisenstein, R. S.; Puccio, H.; Pierik, A. J.; Lill, R., Human Nbp35 is essential for both cytosolic iron-sulfur protein assembly and iron homeostasis. Mol Cell Biol 2008, 28 (17), 5517–28.

(30) Mimaki, M.; Wang, X.; McKenzie, M.; Thorburn, D. R.; Ryan, M. T., Understanding mitochondrial complex I assembly in health and disease. Biochimica et Biophysica Acta (BBA) - Bioenergetics 2012, 1817 (6), 851–862.

(31) Shirakawa, F.; Mizel, S. B., In vitro activation and nuclear translocation of NF-kappa B catalyzed by cyclic AMP-dependent protein kinase and protein kinase C. Mol Cell Biol 1989, 9 (6), 2424–30.

(32) Kane, L. P.; Shapiro, V. S.; Stokoe, D.; Weiss, A., Induction of NF-kappaB by the Akt/PKB kinase. Curr Biol 1999, 9 (11), 601–4.

(33) Ling, X.; Yao, D.; Kang, L.; Zhou, J.; Zhou, Y.; Dong, H.; Zhang, K.; Zhang, L.; Chen, H., Involment of RAS/ERK1/2 signaling and MEF2C in miR-155-3p inhibition-triggered cardiomyocyte differentiation of embryonic stem cell. Oncotarget 2017, 8 (48), 84403–84416.

(34) Basu, S.; Ro, E. J.; Liu, Z.; Kim, H.; Bennett, A.; Kang, S.; Suh, H., The <em>Mef2c</em> Gene Dose-Dependently Controls Hippocampal Neurogenesis and the Expression of Autism-Like Behaviors. The Journal of Neuroscience 2024, 44 (5), e1058232023.

(35) Tempé, D.; Vives, E.; Brockly, F.; Brooks, H.; De Rossi, S.; Piechaczyk, M.; Bossis, G., SUMOylation of the inducible (c-Fos:c-Jun)/AP-1 transcription complex occurs on target promoters to limit transcriptional activation. Oncogene 2014, 33 (7), 921–927.

(36) Soh, J. W.; Lee, E. H.; Prywes, R.; Weinstein, I. B., Novel roles of specific isoforms of protein kinase C in activation of the c-fos serum response element. Mol Cell Biol 1999, 19 (2), 1313–24.

(37) Zheng, W. H.; Fink, D. W., Jr.; Guroff, G., Role of protein kinase C alpha in nerve growth factor-induced arachidonic acid release from PC12 cells. J Neurochem 1996, 66 (5), 1868–75.

(38) Routtenberg, A., Protein kinase C activation leading to protein F1 phosphorylation may regulate synaptic plasticity by presynaptic terminal growth. Behavioral and Neural Biology 1985, 44 (2), 186–200.

(39) Ramakers, G. M. J.; Pasinelli, P.; Hens, J. J. H.; Gispen, W. H.; De Graan, P. N. E., Protein kinase C in synaptic plasticity: Changes in the in situ phosphorylation state of identified pre- and postsynaptic substrates. Progress in Neuro-Psychopharmacology and Biological Psychiatry 1997, 21 (3), 455–486.

(40) Lara Aparicio, S. Y.; Laureani Fierro, Á.d.J.; Aranda Abreu, G. E.; Toledo Cárdenas, R.; García Hernández, L. I.; Coria Ávila, G. A.; Rojas Durán, F.; Aguilar, M. E. H.; Manzo Denes, J.; Chi-Castañeda, L. D.; Pérez Estudillo, C.A., Current Opinion on the Use of c-Fos in Neuroscience. NeuroSci 2022, 3 (4), 687–702.

(41) Milbrandt, J., Nerve growth factor rapidly induces c-fos mRNA in PC12 rat pheochromocytoma cells. Proc Natl Acad Sci U S A 1986, 83 (13), 4789–93.

(42) Zhang, Y.; Tatsuno, T.; Carney, J. M.; Mattson, M. P., Basic FGF, NGF, and IGFs Protect Hippocampal and Cortical Neurons against Iron-Induced Degeneration. Journal of Cerebral Blood Flow & Metabolism 1993, 13 (3), 378–388.

(43) Mwanjewe, J.; Hui, B. K.; Coughlin, M. D.; Grover, A. K., Treatment of PC12 cells with nerve growth factor increases iron uptake. Biochem J 2001, 357 (Pt 3), 881–6.

(44) Chifman, J.; Laubenbacher, R.; Torti, S. V., A systems biology approach to iron metabolism. Adv Exp Med Biol 2014, 844, 201–25.

(45) Ohgami, R. S.; Campagna, D. R.; Greer, E. L.; Antiochos, B.; McDonald, A.; Chen, J.; Sharp, J. J.; Fujiwara, Y.; Barker, J. E.; Fleming, M. D., Identification of a ferrireductase required for efficient transferrin-dependent iron uptake in erythroid cells. Nature Genetics 2005, 37 (11), 1264–1269.

(46) Fleming, M. D.; Romano, M. A.; Su, M. A.; Garrick, L. M.; Garrick, M. D.; Andrews, N. C., NNramp2 is mutated in the anemic Belgrade (b) rat: Evidence of a role for Nramp2 in endosomal iron transport. Proceedings of the National Academy of Sciences 1998, 95 (3), 1148–1153.

(47) Abbaspour, N.; Hurrell, R.; Kelishadi, R., Review on iron and its importance for human health. J Res Med Sci 2014, 19 (2), 164–74.

(48) Weiss, G.; Goossen, B.; Doppler, W.; Fuchs, D.; Pantopoulos, K.; Werner-Felmayer, G.; Wachter, H.; Hentze, M. W., Translational regulation via iron-responsive elements by the nitric oxide/NO-synthase pathway. The EMBO Journal 1993, 12 (9), 3651–3657.

(49) Leskovjan, A. C.; Kretlow, A.; Lanzirotti, A.; Barrea, R.; Vogt, S.; Miller, L. M., Increased brain iron coincides with early plaque formation in a mouse model of Alzheimer’s disease. NeuroImage 2011, 55 (1), 32–38.

(50) Galante, D.; Cavallo, E.; Perico, A.; D’Arrigo, C., Effect of ferric citrate on amyloid-beta peptides behavior. Biopolymers 2018, 109 (6), e23224.

(51) Telling, N. D.; Everett, J.; Collingwood, J. F.; Dobson, J.; van der Laan, G.; Gallagher, J. J.; Wang, J.; Hitchcock, A. P., Iron Biochemistry is Correlated with Amyloid Plaque Morphology in an Established Mouse Model of Alzheimer’s Disease. Cell Chemical Biology 2017, 24 (10), 1205-1215.e3.

(52) Silvestri, L.; Camaschella, C., A potential pathogenetic role of iron in Alzheimer’s disease. Journal of Cellular and Molecular Medicine 2008, 12 (5a), 1548–1550.

(53) Ahmadi, S.; Ebralidze, I. I.; She, Z.; Kraatz, H.-B., Electrochemical studies of tau protein-iron interactions—Potential implications for Alzheimer’s Disease. Electrochimica Acta 2017, 236, 384–393.

(54) Xie, L.; Zheng, W.; Xin, N.; Xie, J.-W.; Wang, T.; Wang, Z.-Y., Ebselen inhibits iron-induced tau phosphorylation by attenuating DMT1 up-regulation and cellular iron uptake. Neurochemistry International 2012, 61 (3), 334–340.

(55) Nakamura, T.; Lipton, S. A., Redox modulation by S-nitrosylation contributes to protein misfolding, mitochondrial dynamics, and neuronal synaptic damage in neurodegenerative diseases. Cell Death & Differentiation 2011, 18 (9), 1478–1486.

(56) Greenough, M. A.; Lane, D. J. R.; Balez, R.; Anastacio, H. T. D.; Zeng, Z.; Ganio, K.; McDevitt, C. A.; Acevedo, K.; Belaidi, A. A.; Koistinaho, J.; Ooi, L.; Ayton, S.; Bush, A. I., Selective ferroptosis vulnerability due to familial Alzheimer’s disease presenilin mutations. Cell Death & Differentiation 2022, 29 (11), 2123–2136.

(57) Foster, L. J.; De Hoog, C. L.; Mann, M., Unbiased quantitative proteomics of lipid rafts reveals high specificity for signaling factors. Proc Natl Acad Sci U S A 2003, 100 (10), 5813–8.

(58) Shevchenko, A.; Wilm, M.; Vorm, O.; Mann, M., Mass spectrometric sequencing of proteins silver-stained polyacrylamide gels. Anal Chem 1996, 68 (5), 850–8.

(59) Ishihama, Y.; Rappsilber, J.; Andersen, J. S.; Mann, M., Microcolumns with self-assembled particle frits for proteomics. J Chromatogr A 2002, 979 (1-2), 233–9.

(60) Kerr, C. H.; Skinnider, M. A.; Andrews, D. D. T.; Madero, A. M.; Chan, Q. W. T.; Stacey, R. G.; Stoynov, N.; Jan, E.; Foster, L. J., Dynamic rewiring of the human interactome by interferon signaling. Genome Biol 2020, 21 (1), 140.

(61) Tyanova, S.; Temu, T.; Cox, J., The MaxQuant computational platform for mass spectrometry-based shotgun proteomics. Nat Protoc 2016, 11 (12), 2301–2319.

(62) Perez-Riverol, Y.; Bai, J.; Bandla, C.; García-Seisdedos, D.; Hewapathirana, S.; Kamatchinathan, S.; Kundu, D. J.; Prakash, A.; Frericks-Zipper, A.; Eisenacher, M.; Walzer, M.; Wang, S.; Brazma, A.; Vizcaíno, J. A., The PRIDE database resources in 2022: a hub for mass spectrometry-based proteomics evidences. Nucleic Acids Res 2022, 50 (D1), D543–d552.

(63) Tyanova, S.; Temu, T.; Sinitcyn, P.; Carlson, A.; Hein, M. Y.; Geiger, T.; Mann, M.; Cox, J., The Perseus computational platform for comprehensive analysis of (prote)omics data. Nat Methods 2016, 13 (9), 731–40.

(64) Blighe, K., S Rana, and M Lewis, EnhancedVolcano: Publication-ready volcano plots with enhanced colouring and labeling. 2018.

(65) Stael, S.; Miller, L. P.; Fernández-Fernández Á, D.; Van Breusegem, F., Detection of Damage-Activated Metacaspase Activity by Western Blot in Plants. Methods Mol Biol 2022, 2447, 127–137.

