## Supplementary Information for "Proteomic Investigation of Neurotrophic *trans*-Banglene Reveals Potential Link to Iron Homeostasis"

Electronic Supplementary Information (ESI)

Piyumi B. Wijesiri Gunawardana,[a] Khyati Gohil,[b†] Kyung-Mee Moon,[c] Leonard J. Foster,[c] Florence J. Williams[a*]

[a] Department of Chemistry, University of Iowa, Iowa City, IA, USA
[b] Department of Chemistry, University of Alberta, Edmonton, AB, Canada
c Department of Biochemistry & Molecular Biology, University of British Colombia, Vancouver, Canada


**Table of Contents**


**1.** General information ..................... SI2

**2.** Global proteomics graphs and volcano plots ..................... SI3

**3.** David 6.8 analysis protocol ..................... SI9

**4.** Western blot quantification protocol ..................... SI10

**5.** Western blots for 48-hour treatment ..................... SI11

**6.** Western blots for 2-hour treatment ..................... SI33

**7.** FerroOrange assay ..................... S156

8. NMR spectra ..................... SI57

### 1. General information

HPLC purification was performed using an Agilent 1260 preparatory system, with a C8 column: gradient 60% acetonitrile (ACN) to 100% ACN over 14 min, 20 mL/min, RT= 9.9 min (±*t*-BG) and 10 min ((±*c*-BG). (PrepHT, 21.2x150mm, 7µm particle size)

Nuclear magnetic resonance (NMR) spectra were obtained from one of the following Varian spectrometers: DD2 MR 400MHz.

NMR spectra chemical shifts (δ) are reported in ppm and are referenced to residual protonated solvent ($^{1}H$) and deuterated solvent ($^{13}C$) chemical shifts. Coupling constants (*J*) are reported in Hertz (Hz). The following abbreviations are used: s = singlet, d = doublet, t = triplet, q = quartet, dd = doublet of doublets, dq = doublet of quartets, ddd = doublet of doublet of doublets, tdd= triplet of doublet of doublets, m = multiplet.

### 2. Global proteomics

Proteomics analysis by Perseus software is available to see in three spreadsheets, named as tBG_NGF 50% annotated p value and $log_2$foldchange, as tBG_DMSO 50% annotated p value and $log_2$foldchange, and as NGF_DMSO 50% annotated p value and $log_2$foldchange. Each of the spreadsheets contains following columns.

DMSO replicate 1, 2,3,4 = 4 technical replicates of DMSO
NGF replicate 1, 2,3,4 = 4 technical replicates of NGF
*t*BG replicate 1, 2,3,4 = 4 technical replicates of *t*BG
GOBP name = Gene Ontology Biological Process
GOMF name= Gene Ontology Molecular Function
GOCC name= Gene Ontology Cellular component
GOBP slim name
GOCC slim name
KEGG name= Kyoto Encyclopedia of Genes and Genomes
Pfam = Protein families database information
GSEA = Gene Set Enrichment Analysis
Keywords = keywords for each gene
Log2foldchange, p value, t test and q value tBG_NGF
Log2foldchange, p value, t test and q value tBG_DMSO
Log2foldchange, p value, t test and q value NGF_DMSO
Gene names

The following graphs depict the downregulation of general cellular processes, such as replication, chromatin structure, transcription, rRNA and tRNA processing, mitochondrial respiratory chain complex I proteins and nucleosome assembly with NGF and *t*-BG treatments.

(A)

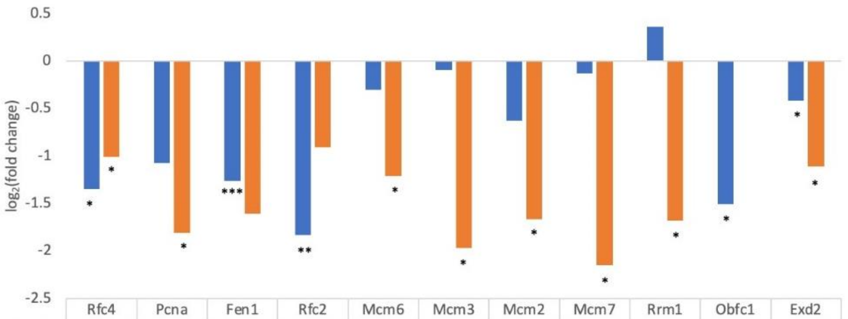


| | Rfc4 | Pcna | Fen1 | Rfc2 | Mcm6 | Mcm3 | Mcm2 | Mcm7 | Rrm1 | Obfc1 | Exd2 |
|---|---|---|---|---|---|---|---|---|---|---|---|
| NGF | -1.35065 | -1.07273 | -1.26223 | -1.83255 | -0.30181 | -0.09359 | -0.62966 | -0.12554 | 0.3628 | -1.50709 | -0.41594 |
| t-BG | -1.0057 | -1.81175 | -1.60926 | -0.91056 | -1.21074 | -1.96517 | -1.6661 | -2.15092 | -1.67756 | 0 | -1.10992 |

(B)

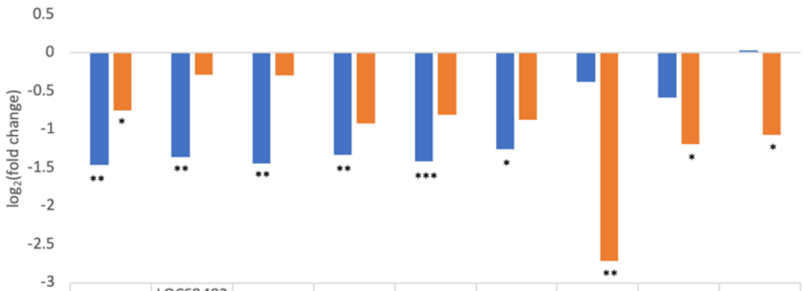


| | Hist1h1a | LOC684828 | H1f0 | H2afy2 | Hist1h1e | Anp32e | Smarce1 | Morf4l2 | Hltf |
|---|---|---|---|---|---|---|---|---|---|
| NGF | -1.46226 | -1.3607 | -1.44778 | -1.32817 | -1.41968 | -1.25405 | -0.376929 | -0.587239 | 0.0285196 |
| t-BG | -0.7552 | -0.287239 | -0.299763 | -0.918885 | -0.813499 | -0.874076 | -2.71446 | -1.19143 | -1.07241 |

Figure 1: Global proteomic analysis with significant protein expression fold change (>2-fold, $p<0.05$) compared to control (A) Cellular replication (B) Chromatin structure *$p<0.05$, **$p<0.01$, ***$p<0.001$ (Y axis: $\log_2$fold change and X axis: gene name with $\log_2$fold change value for respective treatment)

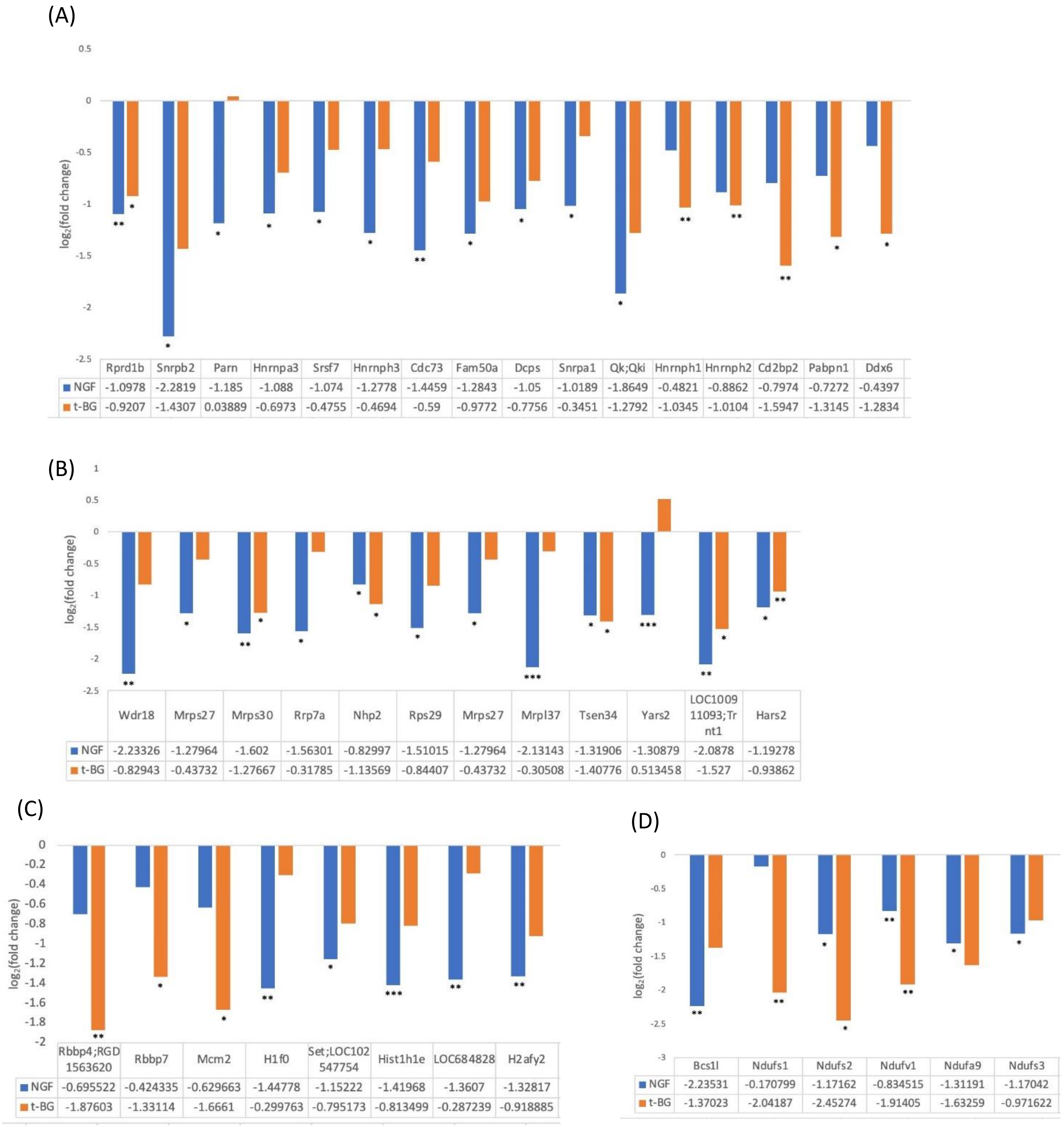


(A)

| | Rprd1b | Snrpb2 | Parn | Hnrnpa3 | Srsf7 | Hnrnph3 | Cdc73 | Fam50a | Dcps | Snrpa1 | Qk;Qki | Hnrnph1 | Hnrnph2 | Cd2bp2 | Pabpn1 | Ddx6 |
|---|---|---|---|---|---|---|---|---|---|---|---|---|---|---|---|---|
| NGF | -1.0978 | -2.2819 | -1.185 | -1.088 | -1.074 | -1.2778 | -1.4459 | -1.2843 | -1.05 | -1.0189 | -1.8649 | -0.4821 | -0.8862 | -0.7974 | -0.7272 | -0.4397 |
| t-BG | -0.9207 | -1.4307 | 0.03889 | -0.6973 | -0.4755 | -0.4694 | -0.59 | -0.9772 | -0.7756 | -0.3451 | -1.2792 | -1.0345 | -1.0104 | -1.5947 | -1.3145 | -1.2834 |

(B)

| | Wdr18 | Mrps27 | Mrps30 | Rrp7a | Nhp2 | Rps29 | Mrps27 | Mrpl37 | Tsen34 | Yars2 | LOC100911093;Trnt1 | Hars2 |
|---|---|---|---|---|---|---|---|---|---|---|---|---|
| NGF | -2.23326 | -1.27964 | -1.602 | -1.56301 | -0.82997 | -1.51015 | -1.27964 | -2.13143 | -1.31906 | -1.30879 | -2.0878 | -1.19278 |
| t-BG | -0.82943 | -0.43732 | -1.27667 | -0.31785 | -1.13569 | -0.84407 | -0.43732 | -0.30508 | -1.40776 | 0.513458 | -1.527 | -0.93862 |

(C)

| | Rbbp4;RGD1563620 | Rbbp7 | Mcm2 | H1f0 | Set;LOC102547754 | Hist1h1e | LOC684828 | H2afy2 |
|---|---|---|---|---|---|---|---|---|
| NGF | -0.695522 | -0.424335 | -0.629663 | -1.44778 | -1.15222 | -1.41968 | -1.3607 | -1.32817 |
| t-BG | -1.87603 | -1.33114 | -1.6661 | -0.299763 | -0.795173 | -0.813499 | -0.287239 | -0.918885 |

(D)

| | Bcs1l | Ndufs1 | Ndufs2 | Ndufv1 | Ndufa9 | Ndufs3 |
|---|---|---|---|---|---|---|
| NGF | -2.23531 | -0.170799 | -1.17162 | -0.834515 | -1.31191 | -1.17042 |
| t-BG | -1.37023 | -2.04187 | -2.45274 | -1.91405 | -1.63259 | -0.971622 |

Figure 1: Global proteomic analysis with significant protein expression fold change (>2-fold, $p<0.05$) compared to control (A) Transcription related (B) rRNA and tRNA processing (C) Nucleosome assembly (D) Mitochondrial respiratory complex 1 *$p<0.05$, **$p<0.01$, ***$p<0.001$ (Y axis: $log_2$fold change and X axis: gene name with $log_2$fold change value for respective treatment)

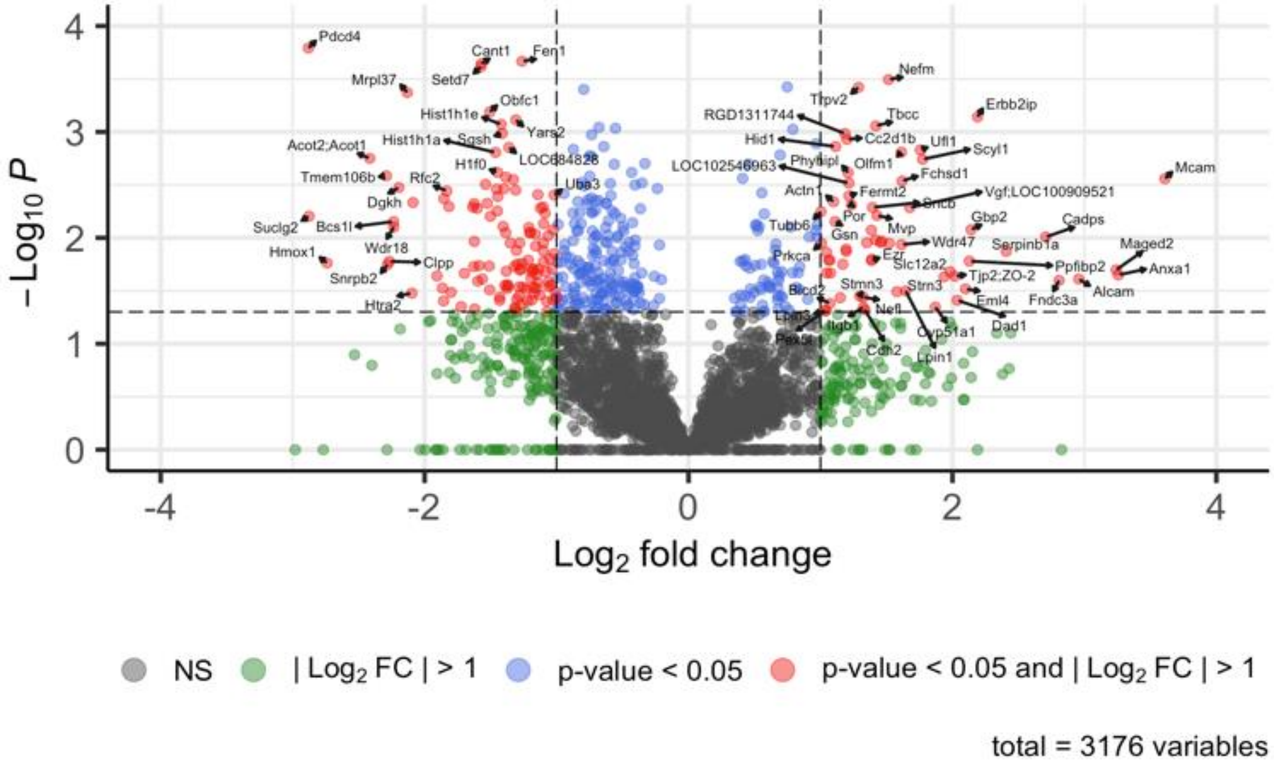


Figure 2: Volcano plots of proteins identified by global proteomic analysis: NGF vs control

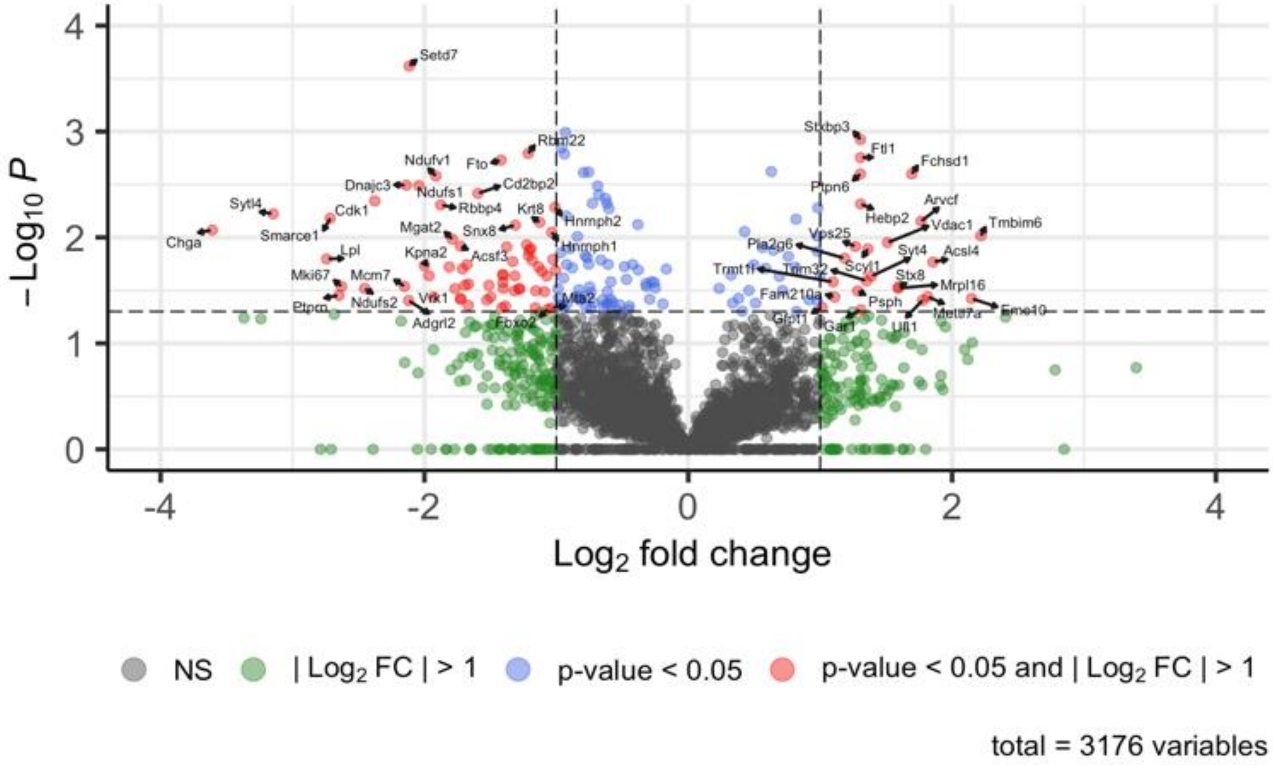


Figure 3: Volcano plots of proteins identified by global proteomic analysis: *t*-BG vs control

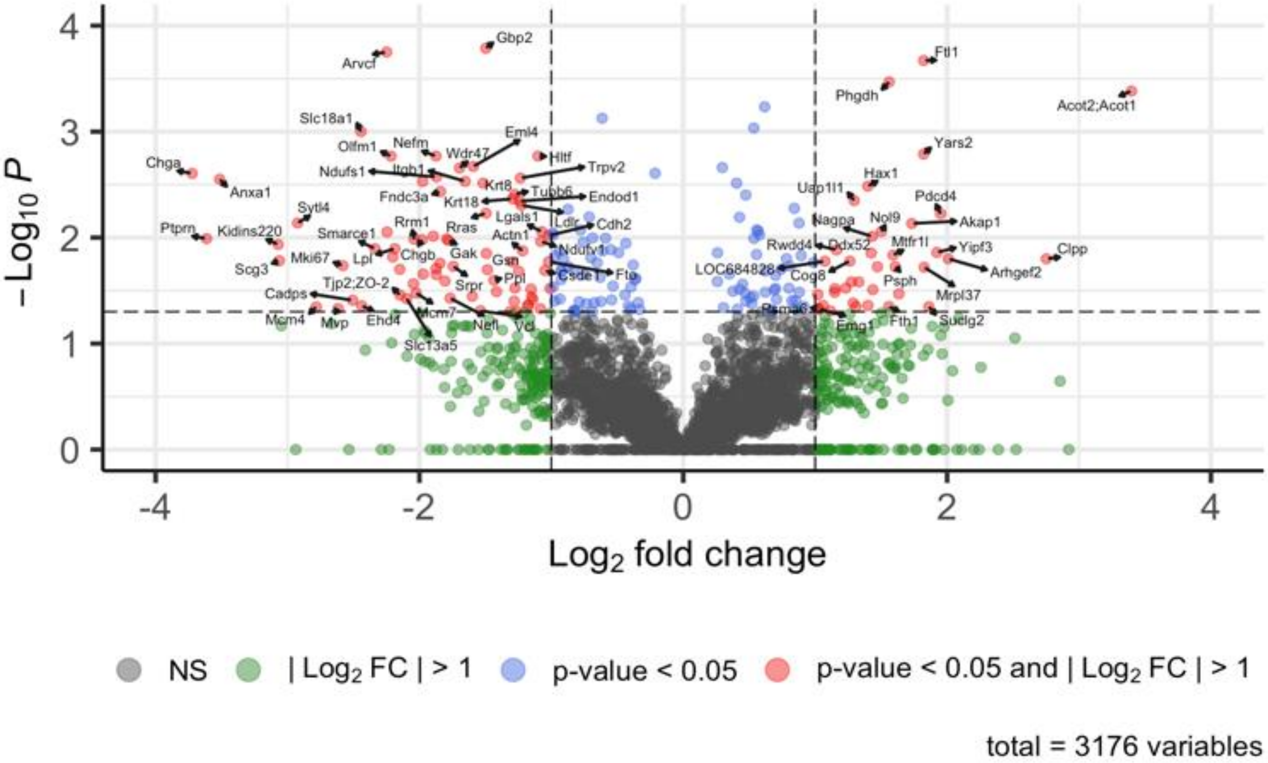


Figure 4: Volcano plots of proteins identified by global proteomic analysis: *t*-BG vs NGF

### 3. DAVID 6.8 Analysis Protocol

All proteins identified as having statistically significant expression level changes ($p<0.05$) from *trans*-banglene treatment as compared to DMSO control were uploaded as a list into DAVID 6.8 (https://david.ncifcrf.gov/). Following submission, the KEGG pathway was selected, and an Annotation Chart was generated. Sorting by count and then eliminating $p>0.05$ results (adjusted Fisher-Exact) generated the Table 1 in the manuscript. The full results can be found in "DAVID analyses" excel spreadsheet under the tab "KEGG pathway chart *t*BDvD." For all of the top hits listed in Table 1 of the manuscript, protein lists associated with these categories can be found in the same spreadsheet under the tab "Protein hits".

For comparison of the effects of *trans*-banglene as compared to NGF, all proteins identified as having statistically significant expression level changes ($p<0.05$) from trans-banglene treatment as compared to NGF treatment were submitted as a list into DAVID 6.8. Following submission, the default analysis category selections were used (UP_KW Biological Processes, Cellular Component, Molecular Function, PTM, UP_SEQ_Feature, GO_Term BP Direct, CC Direct, MF Direct, UP_KW_Ligand, KEGG_Pathway, Interpro, PIR_Superfamily, Smart, UP_KW_Domain). Then, Functional Annotation Clustering was done. The classification stringency was set to the default "Medium" settings (Similarity Term Overlap = 3, Threshold Similarity = 0.50, Initial Group Membership = 3, Final Group Membership = 3, Multiple Linkage Threshold = 0.50, EASE = 1.0). The resulting clusters generated are listed in the David Analyses1 spreadsheet under "tBDvN Annotation Clusters" tab.

Both the Annotation Chart and Annotation Clusters results in the David Analyses spreadsheet include adjusted p-values, Benjamini values, Bonferroni values, and FDR values.

### 4. <u>Western blot quantification protocol[1]</u>

Fiji ImageJ software was used. Following steps were implemented.

1. The western blot file was opened using *File ------> open ------> JPG file*
2. The rotation of the image was adjusted using *Image ------> transform ------> rotate*
3. The brightness was adjusted using *Image ------> adjust ------> Brightness/contrast------> Auto*
4. The image was converted to 32 bit using *Image ------> Type------> 32 bits*
5. The rectangle tool on ImageJ toolbar was selected to draw a rectangle around the bands in the first lane
6. The rectangle was saved as .roi file using *File------> Save as ------> Selection*
7. The rectangle on the first lane was highlighted using *Analyze------> Gels ------> Select First Lane*
8. The rectangle was dragged to the next lane and highlight the rectangle using *Analyze------> Gels ------> Select Next Lane*
9. Repeat step 8 for each subsequent lanes on the gel
10. Draw the profile plot using *Analyze------> Gels ------> Plot Lanes*
11. Select the straight line tool on ImageJ toolbar and draw a line across the base of the peak in the profile plot
12. Select wand tool on ImageJ toolbar and click in the enclosed peaks to get the measurements in the Results window (The values are represented in the tables as “Band area”.
13. Then, go to *Analyze------> Gels ------> Label Peaks* (The values are represented in the tables as “Area%”.
14. Relative density% was calculated using area% for the protein / area% for the control protein

5. <u>Western blots for 48-hour treatment</u>

***Akt Western blot replica for 48-hour treatment***

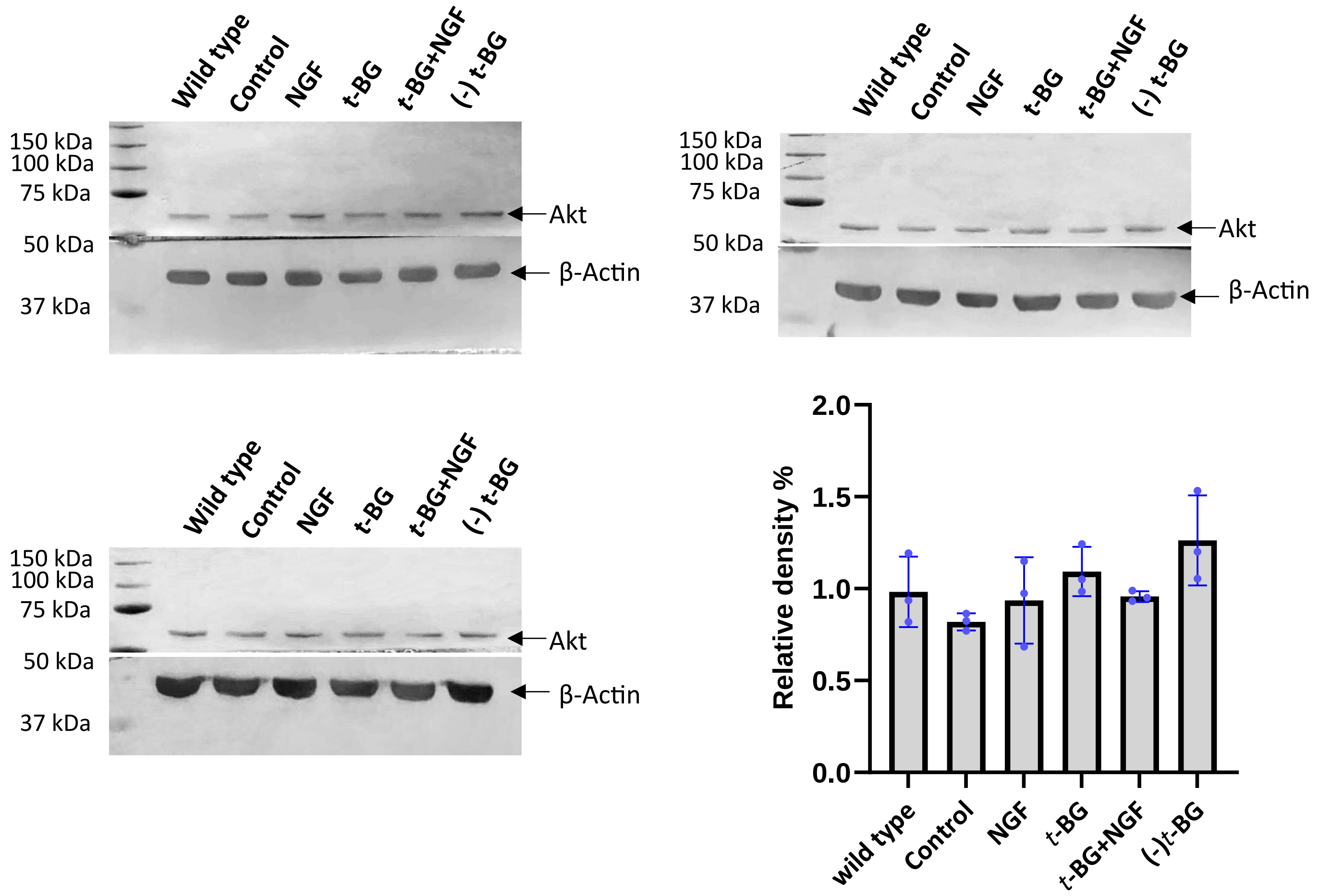

**Table 1: Akt calculations for the replica *without* wild type for 48-hour treatment**

| *Sample name* | *Akt band area* | *Actin band area* | *Akt area%* | *Actin area%* | *Relative density% (Akt area%/actin area%)* | |
|---|---|---|---|---|---|---|
| *Control* | 22086.69 | 43448.81 | 18.646 | 22.269 | 0.837307 | Replicate 1 |
| *NGF-treated* | 17542.4 | 40894.01 | 14.81 | 20.96 | 0.706584 | |
| *t-BG-treated* | 28340.73 | 46075.86 | 23.926 | 23.616 | 1.013127 | |
| *t-BG+NGF-treated* | 21509.4 | 34165.47 | 18.159 | 17.511 | 1.037005 | |
| *(-) t-BG* | 28973.27 | 30521.76 | 24.46 | 15.644 | 1.563539 | |
| *Control* | 22099.81 | 46209.86 | 15.223 | 20.121 | 0.756573 | Replicate 2 |
| *NGF-treated* | 32124.23 | 45446.76 | 22.128 | 19.788 | 1.118253 | |
| *t-BG-treated* | 26302.63 | 42258.9 | 18.118 | 18.4 | 0.984674 | |
| *t-BG+NGF-treated* | 25655.68 | 45039.48 | 17.673 | 19.611 | 0.901178 | |
| *(-) t-BG* | 38989.93 | 50708.7 | 26.858 | 22.08 | 1.216395 | |
| *Control* | 15799.06 | 50507.68 | 16.625 | 20.539 | 0.809436 | Replicate 3 |
| *NGF-treated* | 20060.91 | 53422.66 | 21.11 | 21.724 | 0.971736 | |
| *t-BG-treated* | 21760 | 44830.53 | 22.898 | 18.23 | 1.256061 | |
| *t-BG+NGF-treated* | 14762.89 | 40579.69 | 15.535 | 16.501 | 0.941458 | |
| *(-) t-BG* | 22646.83 | 56575.9 | 23.831 | 23.006 | 1.03586 | |

**Table 2: Akt calculations for the replica *with* wild type for 48-hour treatment**

| *Sample name* | *Akt band area* | *Actin band area* | *Akt area%* | *Actin area%* | *Relative density% (Akt area%/actin area%)* | |
|---|---|---|---|---|---|---|
| *Wild type* | 28802 | 45159.51 | 17.173 | 20.474 | 1.19222 | Replicate 1 |
| *Control* | 20167.03 | 48923.76 | 18.605 | 14.336 | 0.770546 | |
| *NGF-treated* | 16600.69 | 45406.84 | 17.267 | 11.801 | 0.683442 | |
| *t-BG-treated* | 26888.19 | 51067.69 | 19.42 | 19.114 | 0.984243 | |
| *t-BG+NGF-treated* | 20255.57 | 38293.71 | 14.562 | 14.399 | 0.988806 | |
| *(-) t-BG* | 27959.56 | 34110.83 | 12.972 | 19.876 | 1.532223 | |
| *Wild type* | 23047.34 | 46413.06 | 16.773 | 13.732 | 0.818697 | Replicate 2 |
| *Control* | 22903.48 | 45818.15 | 16.558 | 13.646 | 0.824133 | |
| *NGF-treated* | 31779.99 | 45566.29 | 16.467 | 18.935 | 1.149876 | |
| *t-BG-treated* | 25948.38 | 40685.48 | 14.703 | 15.46 | 1.051486 | |
| *t-BG+NGF-treated* | 25525.43 | 45154.73 | 16.319 | 15.208 | 0.93192 | |
| *(-) t-BG* | 38635.69 | 53070.11 | 19.179 | 23.019 | 1.200219 | |
| *Wild type* | 20760.2 | 58094.58 | 19.162 | 17.937 | 0.936071 | Replicate 3 |
| *Control* | 16599.2 | 50344.68 | 16.605 | 14.342 | 0.863716 | |
| *NGF-treated* | 19844.66 | 53358.48 | 17.599 | 17.146 | 0.97426 | |
| *t-BG-treated* | 21283.22 | 44841.41 | 14.79 | 18.388 | 1.243272 | |
| *t-BG+NGF-treated* | 14768.3 | 40697.27 | 13.423 | 12.76 | 0.950607 | |
| *(-) t-BG* | 22486.71 | 55846.9 | 18.42 | 19.428 | 1.054723 | |

*Catalase Western blot replica for 48-hour treatment*

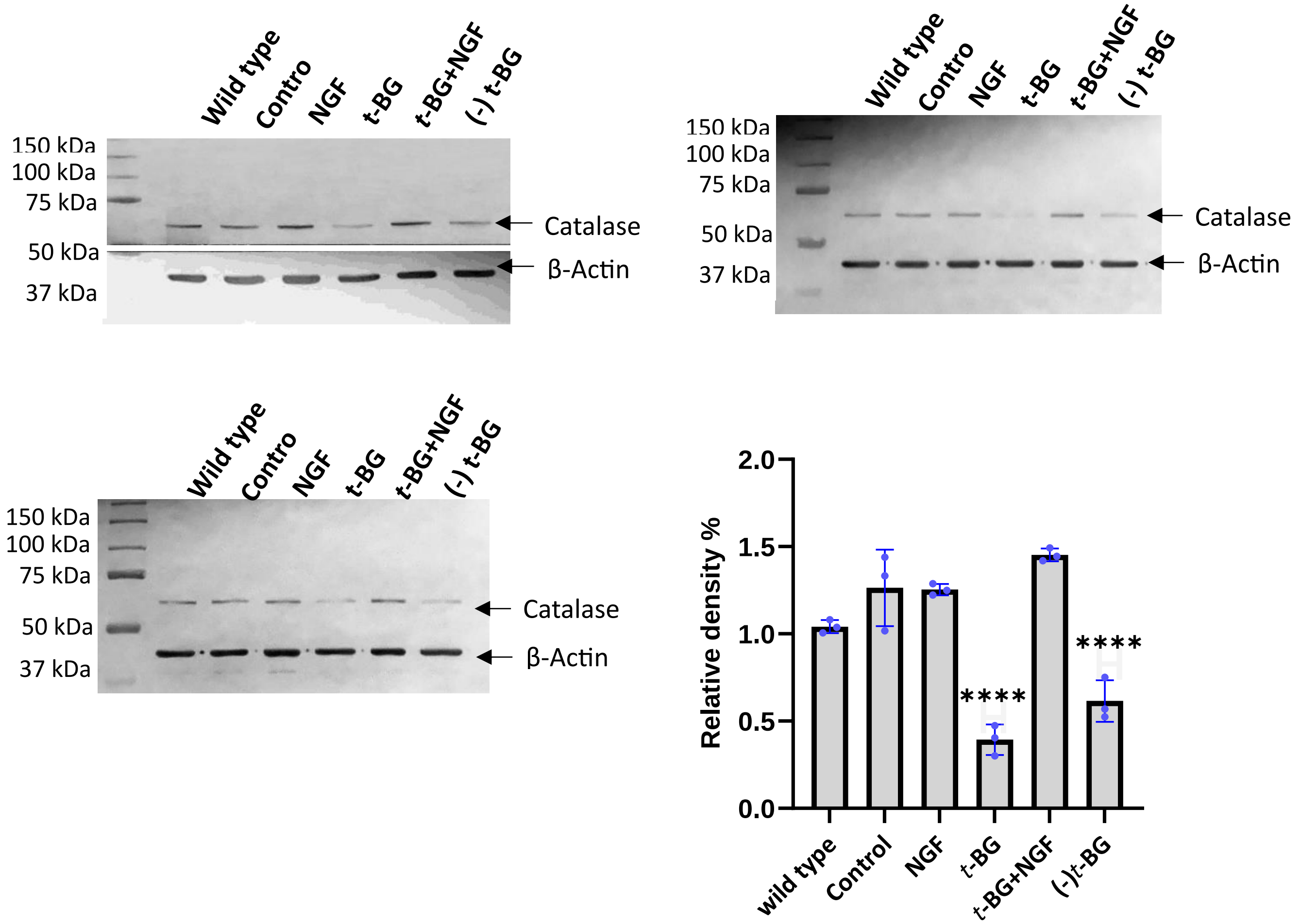

**Table 3: Catalase calculations for the replica *without* wild type for 48-hour treatment**

| *Sample name* | *Catalase band area* | *Actin band area* | *Catalase area%* | *Actin area%* | *Relative density% (Catalase area%/actin area%)* | |
|---|---|---|---|---|---|---|
| *Control* | 3081.841 | 13175.73 | 26.471 | 18.327 | 1.444372 | Replicate 1 |
| *NGF-treated* | 3021.255 | 14143.61 | 25.95 | 19.673 | 1.319067 | |
| *t-BG-treated* | 760.527 | 15832.97 | 6.532 | 22.023 | 0.296599 | |
| *t-BG+NGF-treated* | 3527.962 | 14700.88 | 30.302 | 20.449 | 1.485833 | |
| *(-) t-BG* | 1250.941 | 14038.64 | 10.745 | 19.527 | 0.550264 | |
| *Control* | 2843.79 | 12476.64 | 22.091 | 17.219 | 1.282943 | Replicate 2 |
| *NGF-treated* | 3517.569 | 15816.75 | 27.325 | 21.829 | 1.251775 | |
| *t-BG-treated* | 1182.719 | 13797.18 | 9.187 | 19.041 | 0.482485 | |
| *t-BG+NGF-treated* | 3762.669 | 14377.42 | 29.228 | 19.842 | 1.473037 | |
| *(-) t-BG* | 1566.548 | 15990.58 | 12.169 | 22.069 | 0.551407 | |
| *Control* | 11302.02 | 30525.04 | 19.26 | 18.685 | 1.030773 | Replicate 3 |
| *NGF-treated* | 15782.34 | 33958.87 | 26.895 | 20.787 | 1.293837 | |
| *t-BG-treated* | 4408.468 | 31867.35 | 7.513 | 19.507 | 0.385144 | |
| *t-BG+NGF-treated* | 18959.26 | 35696.4 | 32.309 | 21.851 | 1.478605 | |
| *(-) t-BG* | 8229.267 | 31318.5 | 14.024 | 19.171 | 0.731522 | |

**Table 4: Catalase calculations for the replica *with* wild type for 48-hour treatment**

| *Sample name* | *Catalase band area* | *Actin band area* | *Catalase area%* | *Actin area%* | *Relative density% (Catalase area%/actin area%)* | |
|---|---|---|---|---|---|---|
| *Wild type* | 2471.548 | 14701.22 | 17.505 | 16.874 | 1.037395 | Replicate 1 |
| *Control* | 3087.548 | 13232.61 | 21.868 | 15.189 | 1.439726 | |
| *NGF-treated* | 2996.962 | 14799.29 | 21.227 | 16.987 | 1.249603 | |
| *t-BG-treated* | 776.527 | 15897.15 | 5.5 | 18.247 | 0.301419 | |
| *t-BG+NGF-treated* | 3486.962 | 14423.35 | 24.697 | 16.555 | 1.491815 | |
| *(-) t-BG* | 1299.184 | 14068.47 | 9.202 | 16.148 | 0.569854 | |
| *Wild type* | 2628.134 | 13281.37 | 16.761 | 15.503 | 1.081146 | Replicate 2 |
| *Control* | 3018.79 | 12377.93 | 19.253 | 14.448 | 1.332572 | |
| *NGF-treated* | 3512.861 | 15689.63 | 22.404 | 18.314 | 1.223326 | |
| *t-BG-treated* | 1205.134 | 13885 | 7.686 | 16.207 | 0.47424 | |
| *t-BG+NGF-treated* | 3794.255 | 14608.49 | 24.199 | 17.052 | 1.41913 | |
| *(-) t-BG* | 1520.426 | 15829.34 | 9.697 | 18.477 | 0.524815 | |
| *Wild type* | 10819.15 | 29821.02 | 15.562 | 15.464 | 1.006337 | Replicate 3 |
| *Control* | 11140.32 | 30351.45 | 16.024 | 15.739 | 1.018108 | |
| *NGF-treated* | 15726.51 | 33891.4 | 22.621 | 17.575 | 1.287112 | |
| *t-BG-treated* | 4599.711 | 31509.35 | 6.616 | 16.34 | 0.404896 | |
| *t-BG+NGF-treated* | 18791.84 | 36074.99 | 27.03 | 18.707 | 1.444914 | |
| *(-) t-BG* | 8444.803 | 31192.09 | 12.147 | 16.175 | 0.750974 | |

*Cfos1 western blot replica 48-hour treatment*

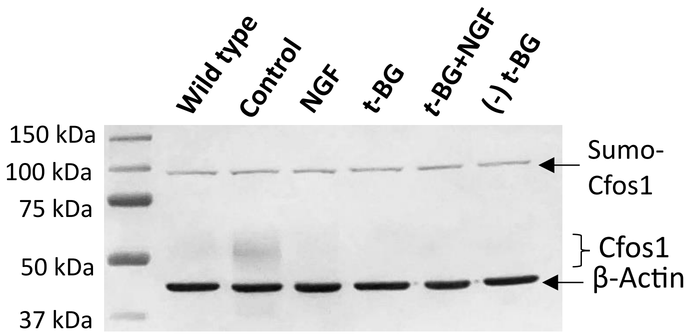


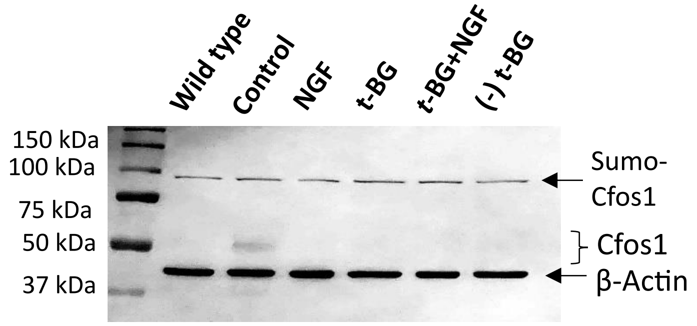


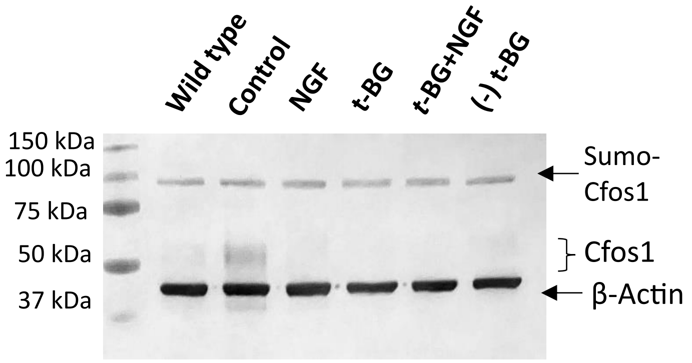


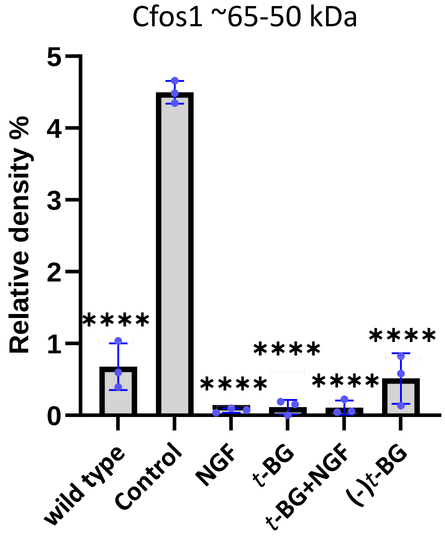


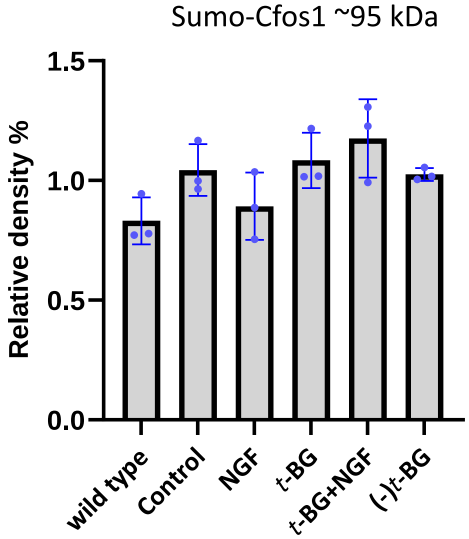

**Table 5: Sumo-Cfos 1 ~95 kDa calculations for the replica *without* wild type for 48-hour treatment**

| *Sample name* | *Cfos 1 band area* | *Actin band area* | *Cfos 1 area%* | *Actin area%* | *Relative density% (Cfos 1 area%/actin area%)* | |
|---|---|---|---|---|---|---|
| *Control* | 1215.749 | 9443.69 | 19.384 | 20.259 | 0.956809 | Replicate 1 |
| *NGF-treated* | 1185.335 | 10019.71 | 18.899 | 21.495 | 0.879228 | |
| *t-BG-treated* | 1323.456 | 9764.912 | 21.101 | 20.948 | 1.007304 | |
| *t-BG+NGF-treated* | 1325.163 | 8076.518 | 21.128 | 17.326 | 1.219439 | |
| *(-) t-BG* | 1222.255 | 9309.205 | 19.488 | 19.971 | 0.975815 | |
| *Control* | 1674.627 | 11590.61 | 19.687 | 20.637 | 0.953966 | Replicate 2 |
| *NGF-treated* | 1366.113 | 12519.53 | 16.06 | 22.291 | 0.72047 | |
| *t-BG-treated* | 2039.749 | 11581.05 | 23.979 | 20.62 | 1.1629 | |
| *t-BG+NGF-treated* | 1861.456 | 9840.296 | 21.883 | 17.52 | 1.24903 | |
| *(-) t-BG* | 1564.284 | 10633.72 | 18.39 | 18.933 | 0.97132 | |
| *Control* | 2272.426 | 12569.28 | 22.029 | 19.913 | 1.106262 | Replicate 3 |
| *NGF-treated* | 2175.477 | 13545.56 | 21.089 | 21.46 | 0.982712 | |
| *t-BG-treated* | 1836.891 | 11641.93 | 17.807 | 18.444 | 0.965463 | |
| *t-BG+NGF-treated* | 1901.648 | 12355.25 | 18.435 | 19.574 | 0.941811 | |
| *(-) t-BG* | 2129.012 | 13008.2 | 20.639 | 20.609 | 1.001456 | |

**Table 6: Sumo-Cfos 1 ~95 kDa calculations for the replica *without* wild type for 48-hour treatment**

| *Sample name* | *Cfos 1 band area* | *Actin band area* | *Cfos 1 area%* | *Actin area%* | *Relative density% (Cfos 1 area%/actin area%)* | |
|---|---|---|---|---|---|---|
| *Wild type* | 1279.577 | 10152.4 | 16.889 | 17.878 | 0.944681 | Replicate 1 |
| *Control* | 1215.749 | 9443.69 | 16.047 | 16.63 | 0.964943 | |
| *NGF-treated* | 1185.335 | 10019.71 | 15.645 | 17.644 | 0.886704 | |
| *t-BG-treated* | 1323.456 | 9764.912 | 17.468 | 17.195 | 1.015877 | |
| *t-BG+NGF-treated* | 1325.163 | 8096.569 | 17.491 | 14.258 | 1.22675 | |
| *(-) t-BG* | 1246.991 | 9310.376 | 16.459 | 16.395 | 1.003904 | |
| *Wild type* | 1331.213 | 11826.46 | 13.532 | 17.394 | 0.777969 | Replicate 2 |
| *Control* | 1674.627 | 11590.61 | 17.023 | 17.047 | 0.998592 | |
| *NGF-treated* | 1366.113 | 12519.53 | 13.887 | 18.413 | 0.754195 | |
| *t-BG-treated* | 2039.749 | 11581.05 | 20.735 | 17.033 | 1.217343 | |
| *t-BG+NGF-treated* | 1861.456 | 9840.296 | 18.922 | 14.473 | 1.3074 | |
| *(-) t-BG* | 1564.284 | 10633.72 | 15.901 | 15.64 | 1.016688 | |
| *Wild type* | 1791.527 | 14959.75 | 14.79 | 19.16 | 0.771921 | Replicate 3 |
| *Control* | 2276.012 | 12569.28 | 18.79 | 16.098 | 1.167226 | |
| *NGF-treated* | 2175.477 | 13545.56 | 17.96 | 17.348 | 1.035278 | |
| *t-BG-treated* | 1839.477 | 11641.93 | 15.186 | 14.91 | 1.01851 | |
| *t-BG+NGF-treated* | 1901.648 | 12355.25 | 15.699 | 15.824 | 0.992101 | |
| *(-) t-BG* | 2129.012 | 13008.2 | 17.576 | 16.66 | 1.054982 | |

**Table 7: Cfos 1 (~65-50 kDa) calculations for the replica *without* wild type for 48-hour treatment**

| *Sample name* | *Cfos 1 band area* | *Actin band area* | *Cfos 1 area%* | *Actin area%* | *Relative density% (Cfos 1 area%/actin area%)* | |
|---|---|---|---|---|---|---|
| *Control* | 4137.267 | 9443.69 | 88.699 | 20.259 | 4.378252 | Replicate 1 |
| *NGF-treated* | 34.536 | 10019.71 | 0.74 | 21.495 | 0.034427 | |
| *t-BG-treated* | 186.778 | 9764.912 | 4.004 | 20.948 | 0.19114 | |
| *t-BG+NGF-treated* | 180.607 | 8076.518 | 3.872 | 17.326 | 0.223479 | |
| *(-) t-BG* | 125.192 | 9309.205 | 2.684 | 19.971 | 0.134395 | |
| *Control* | 1750.861 | 11590.61 | 85.301 | 20.637 | 4.13340 | Replicate 2 |
| *NGF-treated* | 29.243 | 12519.53 | 1.425 | 22.291 | 0.063927 | |
| *t-BG-treated* | 57.778 | 11581.05 | 2.815 | 20.62 | 0.136518 | |
| *t-BG+NGF-treated* | 14.536 | 9840.296 | 0.708 | 17.52 | 0.040411 | |
| *(-) t-BG* | 200.142 | 10633.72 | 9.751 | 18.933 | 0.515027 | |
| *Control* | 2780.66 | 12569.28 | 81.96 | 19.913 | 4.115904 | Replicate 3 |
| *NGF-treated* | 65.536 | 13545.56 | 1.932 | 21.46 | 0.090028 | |
| *t-BG-treated* | Not Detected | 11641.93 | Not Detected | 18.444 | Not Detected | |
| *t-BG+NGF-treated* | 23.95 | 12355.25 | 0.706 | 19.574 | 0.036068 | |
| *(-) t-BG* | 522.577 | 13008.2 | 15.403 | 20.609 | 0.747392 | |

**Table 8: Cfos 1 (~65-50 kDa) calculations for the replica *with* wild type for 48-hour treatment**

| *Sample name* | *Cfos 1 band area* | *Actin band area* | *Cfos 1 area%* | *Actin area%* | *Relative density% (Cfos 1 area%/actin area%)* | |
|---|---|---|---|---|---|---|
| *Wild type* | 1058.497 | 10152.4 | 18.497 | 17.878 | 1.034624 | Replicate 1 |
| *Control* | 4137.267 | 9443.69 | 72.297 | 16.63 | 4.347384 | |
| *NGF-treated* | 34.536 | 10019.71 | 0.603 | 17.644 | 0.034176 | |
| *t-BG-treated* | 186.485 | 9764.912 | 3.259 | 17.195 | 0.189532 | |
| *t-BG+NGF-treated* | 180.607 | 8096.569 | 3.156 | 14.258 | 0.221349 | |
| *(-) t-BG* | 125.192 | 9310.376 | 2.188 | 16.395 | 0.133455 | |
| *Wild type* | 151.607 | 11826.46 | 6.878 | 17.394 | 0.395424 | Replicate 2 |
| *Control* | 1750.861 | 11590.61 | 79.434 | 17.047 | 4.659706 | |
| *NGF-treated* | 29.243 | 12519.53 | 1.327 | 18.413 | 0.072069 | |
| *t-BG-treated* | 57.778 | 11581.05 | 2.621 | 17.033 | 0.153878 | |
| *t-BG+NGF-treated* | 14.536 | 9840.296 | 0.659 | 14.473 | 0.045533 | |
| *(-) t-BG* | 200.142 | 10633.72 | 9.08 | 15.64 | 0.580563 | |
| *Wild type* | 446.577 | 14959.75 | 11.491 | 19.16 | 0.599739 | Replicate 3 |
| *Control* | 2806.711 | 12569.28 | 72.218 | 16.098 | 4.486147 | |
| *NGF-treated* | 65.536 | 13545.56 | 1.686 | 17.348 | 0.097187 | |

| | | | | | |
|---|---|---|---|---|---|
| *t-BG-treated* | Not Detected | 11641.93 | Not Detected | 14.91 | Not Detected |
| *t-BG+NGF-treated* | 33.536 | 12355.25 | 0.863 | 15.824 | 0.054537 |
| *(-) t-BG* | 534.092 | 13008.2 | 13.742 | 16.66 | 0.82485 |

*Erk1/2 western blot replica 48-hour treatment*

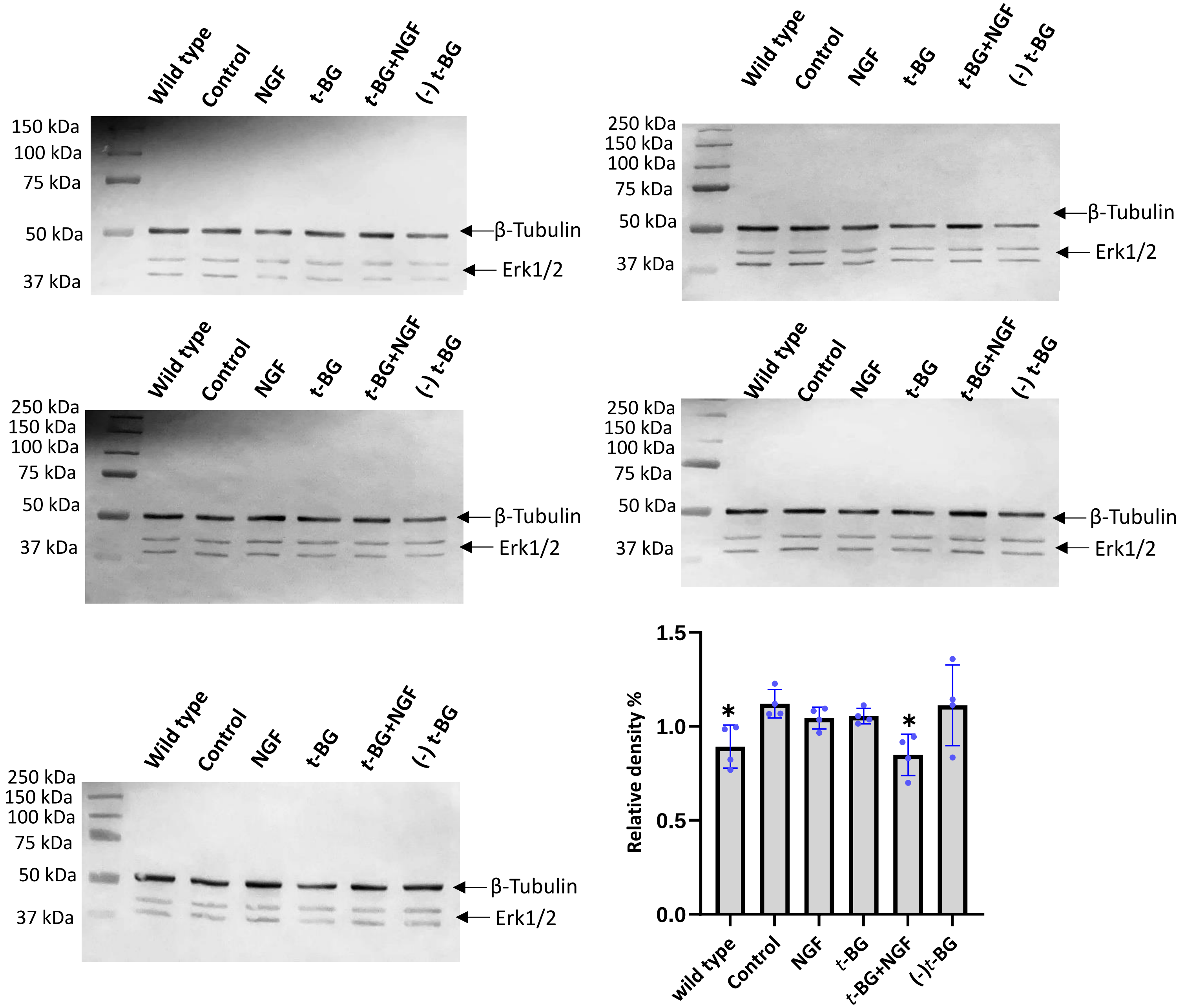

**Table 9: Erk1/2 calculations for the replica *without* wild type for 48-hour treatment**

| *Sample name* | *Erk1/2 band area* | *Tubulin band area* | *Erk1/2 area%* | *Tubulin area%* | *Relative density% (Erk1/2 area%/tubulin area%)* | |
|---|---|---|---|---|---|---|
| *Control* | 7107.681 | 11295.25 | 25.047 | 24.552 | 1.020161 | Replicate 1 |
| *NGF-treated* | 5477.531 | 8987.225 | 19.303 | 19.535 | 0.988124 | |
| *t-BG-treated* | 5734.146 | 8360.033 | 20.207 | 18.172 | 1.111985 | |
| *t-BG+NGF-treated* | 4829.903 | 11205.52 | 17.021 | 24.357 | 0.698813 | |
| *(-) t-BG* | 5227.903 | 6157.719 | 18.423 | 13.385 | 1.376391 | |
| *Control* | 5869.945 | 11611.2 | 20.175 | 18.615 | 1.083803 | Replicate 2 |
| *NGF-treated* | 6793.188 | 15031.8 | 23.348 | 24.099 | 0.968837 | |
| *t-BG-treated* | 6193.409 | 13283.68 | 21.287 | 21.296 | 0.999577 | |
| *t-BG+NGF-treated* | 5323.995 | 12720.15 | 18.299 | 20.393 | 0.897318 | |
| *(-) t-BG* | 4914.359 | 9728.66 | 16.89 | 15.597 | 1.082901 | |
| *Control* | 6382.752 | 13146.9 | 21.13 | 21.168 | 0.998205 | Replicate 3 |
| *NGF-treated* | 5235.995 | 10414.73 | 17.333 | 16.769 | 1.033633 | |
| *t-BG-treated* | 5817.359 | 12320.17 | 19.258 | 19.837 | 0.970812 | |
| *t-BG+NGF-treated* | 6673.845 | 14786.41 | 22.094 | 23.808 | 0.928007 | |
| *(-) t-BG* | 6097.48 | 11438.39 | 20.185 | 18.417 | 1.095998 | |
| *Control* | 5033.317 | 12865.42 | 26.529 | 21.522 | 1.232646 | Replicate 4 |
| *NGF-treated* | 3364.145 | 9653.64 | 17.731 | 16.149 | 1.097963 | |
| *t-BG-treated* | 4384.046 | 13623.78 | 23.107 | 22.791 | 1.013865 | |
| *t-BG+NGF-treated* | 3691.439 | 14048.83 | 19.456 | 23.502 | 0.827844 | |
| *(-) t-BG* | 2500.246 | 9585.811 | 13.177 | 16.036 | 0.821714 | |
| *Control* | 4246.823 | 13595.58 | 19.723 | 19.672 | 1.002593 | Replicate 5 |
| *NGF-treated* | 4739.702 | 17628.72 | 22.012 | 25.508 | 0.862945 | |
| *t-BG-treated* | 3133.611 | 10537.49 | 14.553 | 15.247 | 0.954483 | |
| *t-BG+NGF-treated* | 4198.581 | 13169.56 | 19.499 | 19.056 | 1.023247 | |
| *(-) t-BG* | 5213.702 | 14179.02 | 24.213 | 20.516 | 1.180201 | |

**Table 10: Erk1/2 calculations for the replica *with* wild type for 48-hour treatment**

| *Sample name* | *Erk1/2 band area* | *Tubulin band area* | *Erk1/2 area%* | *Tubulin area%* | *Relative density% (Erk1/2 area%/tubulin area%)* | |
|---|---|---|---|---|---|---|
| *Wild type* | 8199.258 | 13264.08 | 22.063 | 22.427 | 0.98377 | Replicate 1 |
| *Control* | 7567.995 | 11272.25 | 20.365 | 19.059 | 1.068524 | |
| *NGF-treated* | 5478.945 | 9030.518 | 14.743 | 15.269 | 0.965551 | |
| *t-BG-treated* | 5813.974 | 8325.033 | 15.645 | 14.076 | 1.111466 | |
| *t-BG+NGF-treated* | 4904.146 | 11158.81 | 13.196 | 18.867 | 0.699422 | |
| *(-) t-BG* | 5198.024 | 6093.719 | 13.987 | 10.303 | 1.357566 | |
| *Wild type* | 4891.51 | 14376.8 | 14.403 | 18.761 | 0.76771 | Replicate 2 |
| *Control* | 5852.358 | 11796.32 | 17.232 | 15.393 | 1.11947 | |
| *NGF-treated* | 6841.944 | 14900.85 | 20.146 | 19.444 | 1.036104 | |
| *t-BG-treated* | 6177.995 | 13214.73 | 18.191 | 17.244 | 1.054918 | |
| *t-BG+NGF-treated* | 5337.995 | 12752.85 | 15.718 | 16.641 | 0.944535 | |
| *(-) t-BG* | 4859.116 | 9591.539 | 14.308 | 12.516 | 1.143177 | |
| *Wild type* | 4708.509 | 12101.66 | 13.664 | 16.592 | 0.823529 | Replicate 3 |
| *Control* | 6575.995 | 13062.49 | 19.084 | 17.91 | 1.06555 | |
| *NGF-treated* | 5279.409 | 10330.32 | 15.321 | 14.164 | 1.081686 | |
| *t-BG-treated* | 5623.823 | 11483.56 | 16.321 | 15.745 | 1.036583 | |
| *t-BG+NGF-treated* | 6398.601 | 14788 | 18.569 | 20.276 | 0.915812 | |
| *(-) t-BG* | 5872.409 | 11169.15 | 17.042 | 15.314 | 1.112838 | |
| *Wild type* | 4381.954 | 13983.66 | 18.933 | 19.033 | 0.994746 | Replicate 4 |
| *Control* | 4942.196 | 12791.13 | 21.354 | 17.41 | 1.226536 | |
| *NGF-treated* | 3290.732 | 9550.225 | 14.218 | 12.999 | 1.093776 | |
| *t-BG-treated* | 4411.167 | 13807.32 | 19.059 | 18.793 | 1.014154 | |
| *t-BG+NGF-treated* | 3662.56 | 13991.54 | 15.824 | 19.044 | 0.830918 | |
| *(-) t-BG* | 2456.367 | 9345.983 | 10.613 | 12.721 | 0.83429 | |
| *Wild type* | 4059.066 | 16166.65 | 15.684 | 19.04 | 0.823739 | Replicate 5 |
| *Control* | 4158.995 | 13551.87 | 16.071 | 15.961 | 1.006892 | |
| *NGF-treated* | 4842.53 | 17694.43 | 18.712 | 20.84 | 0.897889 | |
| *t-BG-treated* | 3387.267 | 10474.49 | 13.089 | 12.336 | 1.061041 | |
| *t-BG+NGF-treated* | 4233.288 | 13074.39 | 16.358 | 15.398 | 1.06234 | |
| *(-) t-BG* | 5198.288 | 13945.08 | 20.087 | 16.424 | 1.223027 | |

*Ferritin Western blot replica for 48-hour treatment*

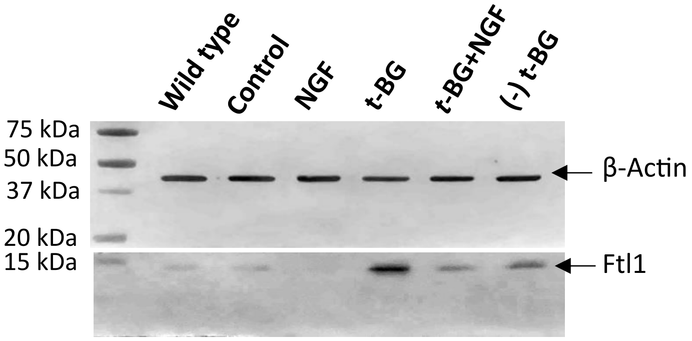


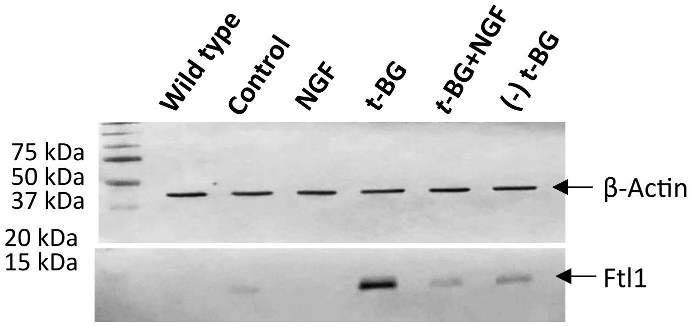


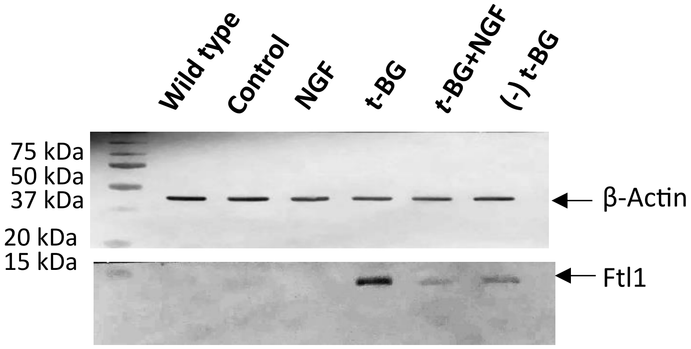


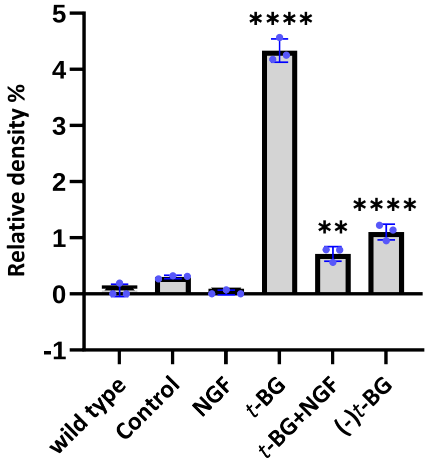

**Table 11: Ferritin calculations for the replica *without* wild type for 48-hour treatment**

| *Sample name* | *Ftl1 band area* | *Actin band area* | *Ftl1 area%* | *Actin area%* | *Relative density% (Ftl1 area%/ Actin area%)* | |
|---|---|---|---|---|---|---|
| *Control* | 3036.083 | 24135.39 | 5.003 | 21.587 | 0.23176 | Replicate 1 |
| *NGF-treated* | 689.506 | 24385.82 | 1.136 | 21.811 | 0.052084 | |
| *t-BG-treated* | 36504.58 | 18541.39 | 60.156 | 16.583 | 3.62757 | |
| *t-BG+NGF-treated* | 7606.995 | 21304.97 | 12.535 | 19.055 | 0.657833 | |
| *(-) t-BG* | 12846.52 | 23439.73 | 21.17 | 20.964 | 1.009826 | |
| *Control* | 2413.154 | 14789.51 | 5.896 | 19.824 | 0.297417 | Replicate 2 |
| *NGF-treated* | Not Detected | 14328.37 | Not Detected | 19.206 | Not Detected | |
| *t-BG-treated* | 28167.25 | 14374.44 | 68.822 | 19.267 | 3.572014 | |
| *t-BG+NGF-treated* | 3710.175 | 15219.49 | 9.065 | 20.4 | 0.444363 | |
| *(-) t-BG* | 6637.338 | 15893.32 | 16.217 | 21.303 | 0.761254 | |
| *Control* | 3431.255 | 22487.49 | 7.426 | 25.15 | 0.295268 | Replicate 3 |
| *NGF-treated* | Not Detected | 17238.22 | Not Detected | 19.28 | Not Detected | |
| *t-BG-treated* | 28295.42 | 15127.13 | 61.236 | 16.918 | 3.619577 | |
| *t-BG+NGF-treated* | 5191.225 | 16530.38 | 11.235 | 18.488 | 0.607691 | |
| *(-) t-BG* | 9289.51 | 18028.68 | 20.104 | 20.164 | 0.997024 | |

**Table 12: Ferritin calculations for the replica *with* wild type for 48-hour treatment**

| *Sample name* | *Ftl1 band area* | *Actin band area* | *Ftl1 area%* | *Actin area%* | *Relative density% (Ftl1 area%/ Actin area%)* | |
|---|---|---|---|---|---|---|
| *Wild type* | 1913.648 | 21690.39 | 3.081 | 16.121 | 0.191117 | Replicate 1 |
| *Control* | 2982.326 | 24176.1 | 4.802 | 17.968 | 0.267253 | |
| *NGF-treated* | 838.163 | 24777.36 | 1.349 | 18.415 | 0.073255 | |
| *t-BG-treated* | 36201.21 | 18772.92 | 58.285 | 13.952 | 4.177537 | |
| *t-BG+NGF treated* | 7788.581 | 21546.8 | 12.54 | 16.014 | 0.783065 | |
| *(-) t-BG* | 12386.5 | 23587.15 | 19.943 | 17.53 | 1.13765 | |
| *Wild type* | NaN | 15254.73 | NaN | 17.23 | NaN | Replicate 2 |
| *Control* | 2175.74 | 14291.61 | 5.184 | 16.142 | 0.32115 | |
| *NGF-treated* | NaN | 14201.54 | NaN | 16.04 | NaN | |
| *t-BG-treated* | 28797.74 | 14281.44 | 68.612 | 16.13 | 4.253689 | |
| *t-BG+NGF treated* | 3942.539 | 14839.49 | 9.393 | 16.761 | 0.560408 | |
| *(-) t-BG* | 7055.995 | 15668.9 | 16.811 | 17.697 | 0.949935 | |
| *Wild type* | NaN | 21595.68 | NaN | 19.883 | NaN | Replicate 3 |
| *Control* | 2946.012 | 21775.61 | 6.309 | 20.049 | 0.314679 | |
| *NGF-treated* | NaN | 17047.22 | NaN | 15.695 | NaN | |
| *t-BG-treated* | 29184.66 | 14856.95 | 62.502 | 13.679 | 4.569194 | |
| *t-BG+NGF treated* | 5396.296 | 15923.72 | 11.557 | 14.661 | 0.788282 | |
| *(-) t-BG* | 9167.095 | 17413.08 | 19.632 | 16.032 | 1.224551 | |

*Mef2c Western blot replica for 48-hour treatment*

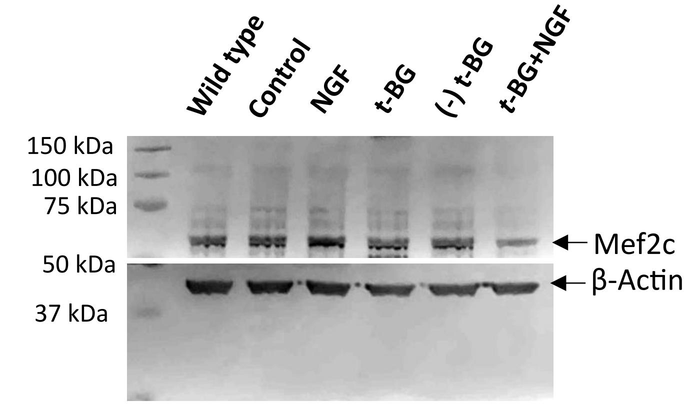


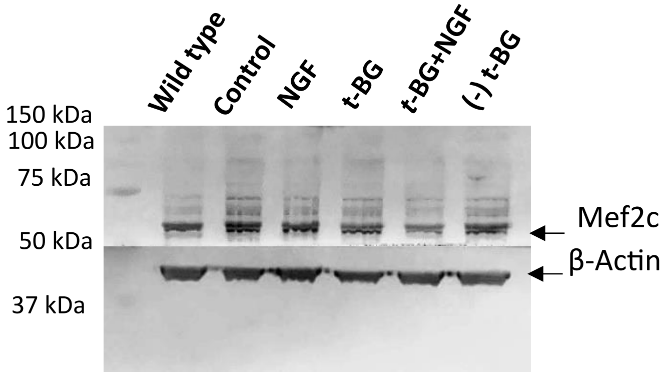


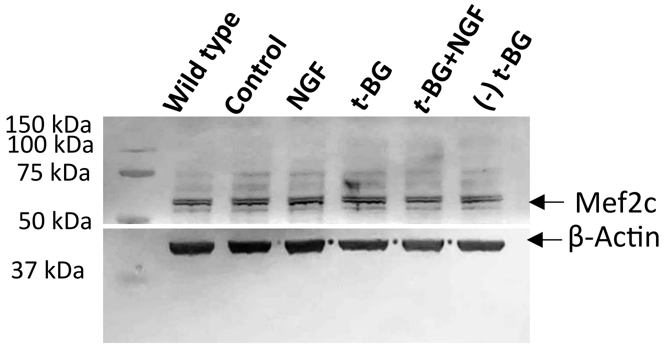


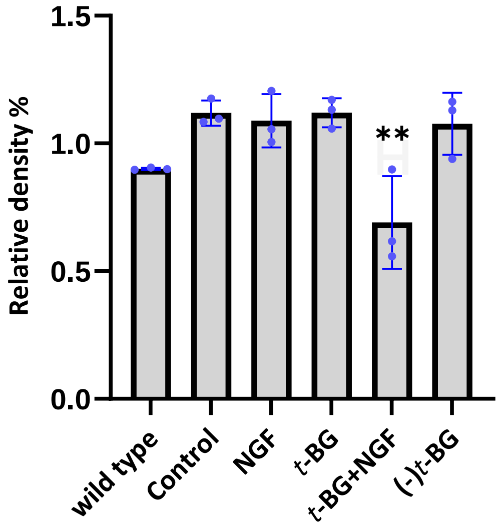

**Table 13: Mef2c calculations for the replica *without* wild type for 48-hour treatment**

| *Sample name* | *Mef2c band area* | *Actin band area* | *Mef2c area%* | *Actin area%* | *Relative density% (Mef2c area%/ Actin area%)* | |
|---|---|---|---|---|---|---|
| *Control* | 19746.25 | 16995.18 | 22.001 | 20.09 | 1.095122 | Replicate 1 |
| *NGF-treated* | 20332.44 | 17814.74 | 22.655 | 21.059 | 1.075787 | |
| *t-BG-treated* | 18148.15 | 15050.92 | 20.221 | 17.792 | 1.136522 | |
| *t-BG+NGF-treated* | 21134.03 | 17549.46 | 23.548 | 20.745 | 0.569853 | |
| *(-) t-BG* | 10389.07 | 17184.92 | 11.576 | 20.314 | 1.135117 | |
| *Control* | 25081.81 | 21606.89 | 21.153 | 18.401 | 1.149557 | Replicate 2 |
| *NGF-treated* | 25730.43 | 25707.08 | 21.7 | 21.893 | 0.991184 | |
| *t-BG-treated* | 25550.26 | 21463.45 | 21.549 | 18.279 | 1.178894 | |
| *t-BG+NGF-treated* | 14202.92 | 23241.62 | 11.978 | 19.793 | 0.605163 | |
| *(-) t-BG* | 28005.51 | 25402.77 | 23.619 | 21.634 | 1.091754 | |
| *Control* | 14099.77 | 22244.48 | 19.837 | 18.896 | 1.049799 | Replicate 3 |
| *NGF-treated* | 14948.66 | 24450.09 | 21.032 | 20.77 | 1.012614 | |
| *t-BG-treated* | 16425.35 | 24269.17 | 23.109 | 20.616 | 1.120925 | |
| *t-BG+NGF-treated* | 11721.79 | 21760.65 | 16.492 | 18.485 | 0.892183 | |
| *(-) t-BG* | 13881.69 | 24993.12 | 19.53 | 21.231 | 0.919881 | |

**Table 14: Mef2c calculations for the replica *with* wild type for 48-hour treatment**

| *Sample name* | *Mef2c band area* | *Actin band area* | *Mef2c area%* | *Actin area%* | *Relative density% (Mef2c area%/ Actin area%)* | |
|---|---|---|---|---|---|---|
| *Wild type* | 16541.99 | 21197.82 | 15.057 | 16.631 | 0.905357 | Replicate 1 |
| *Control* | 19556.13 | 20681.14 | 17.801 | 16.225 | 1.097134 | |
| *NGF-treated* | 24040.51 | 23125.38 | 21.883 | 18.143 | 1.20614 | |
| *t-BG-treated* | 18051.03 | 19786.72 | 16.431 | 15.524 | 1.058426 | |
| *t-BG+NGF-treated* | 10250.36 | 21293.36 | 9.33 | 16.706 | 0.558482 | |
| *(-) t-BG* | 21419.03 | 21376.31 | 19.497 | 16.771 | 1.162542 | |
| *Wild type* | 18999.21 | 21978.5 | 14.353 | 15.966 | 0.898973 | Replicate 2 |
| *Control* | 23564.86 | 20845.23 | 17.802 | 15.142 | 1.17567 | |
| *NGF-treated* | 25892.8 | 25498.54 | 19.56 | 18.523 | 1.055984 | |
| *t-BG-treated* | 22930.93 | 21071.74 | 17.323 | 15.307 | 1.131704 | |
| *t-BG+NGF-treated* | 13728.92 | 23164.62 | 10.371 | 16.827 | 0.616331 | |
| *(-) t-BG* | 27258.29 | 25103.47 | 20.592 | 18.236 | 1.129195 | |
| *Wild type* | 12127.45 | 22978.4 | 14.687 | 16.364 | 0.897519 | Replicate 3 |
| *Control* | 14259.35 | 22021.07 | 17.269 | 15.682 | 1.101199 | |
| *NGF-treated* | 14829.84 | 24956.16 | 17.96 | 17.772 | 1.010578 | |
| *t-BG-treated* | 16476.23 | 24113.34 | 19.953 | 17.172 | 1.16195 | |
| *t-BG+NGF-treated* | 11295.14 | 22003.89 | 13.679 | 15.67 | 0.872942 | |
| *(-) t-BG* | 13585.45 | 24348.58 | 16.453 | 17.34 | 0.948847 | |

*Ndufs1 Western blot replica for 48-hour treatment*

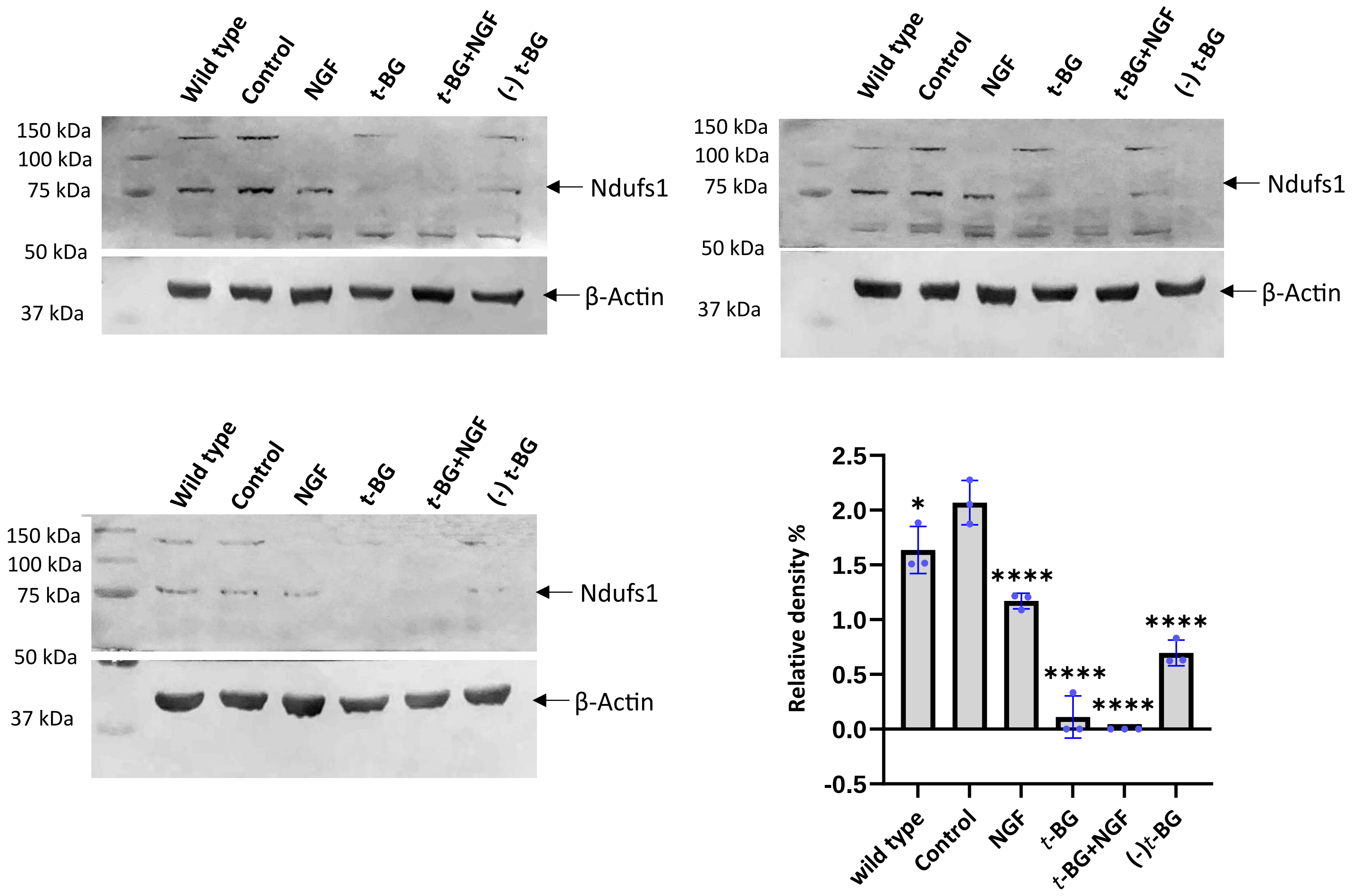

**Table 15: Ndufs1 calculations for the replica *without* type for 48-hour treatment**

| *Sample name* | *Ndufs1 band area* | *Actin band area* | *Ndufs1 area%* | *Actin area%* | *Relative density% (Ndufs1 area%/ Actin area%)* | |
|---|---|---|---|---|---|---|
| *Control* | 19753.19 | 44189.58 | 52.936 | 20.789 | 2.546347 | Replicate 1 |
| *NGF-treated* | 11650.55 | 48585.35 | 31.222 | 22.857 | 1.365971 | |
| *t-BG-treated* | Not Detected | 36814.58 | Not Detected | 17.32 | Not Detected | |
| *t-BG+NGF-treated* | Not Detected | 45691.3 | Not Detected | 21.496 | Not Detected | |
| *(-) t-BG* | 5911.803 | 37279.66 | 15.843 | 17.538 | 0.903353 | |
| *Control* | 21497.46 | 36470.66 | 50.083 | 19.513 | 2.566648 | Replicate 2 |
| *NGF-treated* | 14708.2 | 41646.1 | 29.209 | 22.282 | 1.310879 | |
| *t-BG-treated* | Not Detected | 36272.35 | 7.593 | 19.407 | 0.391251 | |
| *t-BG+NGF-treated* | Not Detected | 37727.01 | Not Detected | 20.185 | Not Detected | |
| *(-) t-BG* | 6455.459 | 34791.98 | 13.115 | 18.614 | 0.704577 | |
| *Control* | 21497.46 | 43727.81 | 50.391 | 21.315 | 2.36411 | Replicate 3 |
| *NGF-treated* | 14708.2 | 50608.76 | 34.477 | 24.669 | 1.397584 | |
| *t-BG-treated* | Not Detected | 37114.56 | Not Detected | 18.091 | Not Detected | |
| *t-BG+NGF-treated* | Not Detected | 32445.52 | Not Detected | 15.815 | Not Detected | |
| *(-) t-BG* | 6455.459 | 41256.56 | 15.132 | 20.11 | 0.752461 | |

**Table 16: Ndufs1 calculations for the replica *with* type for 48-hour treatment**

| *Sample name* | *Ndufs1 band area* | *Actin band area* | *Ndufs1 area%* | *Actin area%* | *Relative density% (Ndufs1 area%/ Actin area%)* | |
|---|---|---|---|---|---|---|
| *Wild type* | 12577.68 | 43255.97 | 25.27 | 16.757 | 1.508026 | Replicate 1 |
| *Control* | 19502.89 | 44421.58 | 39.184 | 17.208 | 2.27708 | |
| *NGF-treated* | 11651.55 | 49698.65 | 23.41 | 19.253 | 1.215914 | |
| *t-BG-treated* | Not Detected | 37359.11 | Not Detected | 14.472 | Not Detected | |
| *t-BG+NGF-treated* | Not Detected | 45710.01 | Not Detected | 17.707 | Not Detected | |
| *(-) t-BG* | 6040.167 | 37694.36 | 12.136 | 14.602 | 0.831119 | |
| *Wild type* | 13735.48 | 44141.98 | 30.63 | 20.216 | 1.515137 | Replicate 2 |
| *Control* | 15331.87 | 36340.1 | 34.19 | 16.643 | 2.054317 | |
| *NGF-treated* | 9307.995 | 37561.74 | 20.757 | 17.202 | 1.206662 | |
| *t-BG-treated* | 2299.548 | 33578.3 | 5.128 | 15.378 | 0.333463 | |

| | | | | | | |
|---|---|---|---|---|---|---|
| *t-BG+NGF-treated* | Not Detected | 34612.47 | Not Detected | 15.851 | Not Detected | |
| *(-) t-BG* | 4167.66 | 32120.44 | 9.294 | 14.71 | 0.631815 | |
| *Wild type* | 21914.64 | 45886.39 | 35.349 | 18.759 | 1.884375 | Replicate 3 |
| *Control* | 20131.81 | 42383.98 | 32.473 | 17.327 | 1.874127 | |
| *NGF-treated* | 13552.84 | 49158.88 | 21.861 | 20.097 | 1.087774 | |
| *t-BG-treated* | Not Detected | 34959.37 | Not Detected | 14.292 | Not Detected | |
| *t-BG+NGF-treated* | Not Detected | 31820.06 | Not Detected | 13.009 | Not Detected | |
| *(-) t-BG* | 6396.459 | 40399.27 | 10.318 | 16.516 | 0.624728 | |

*NFκB western blot replica 48-hour treatment*

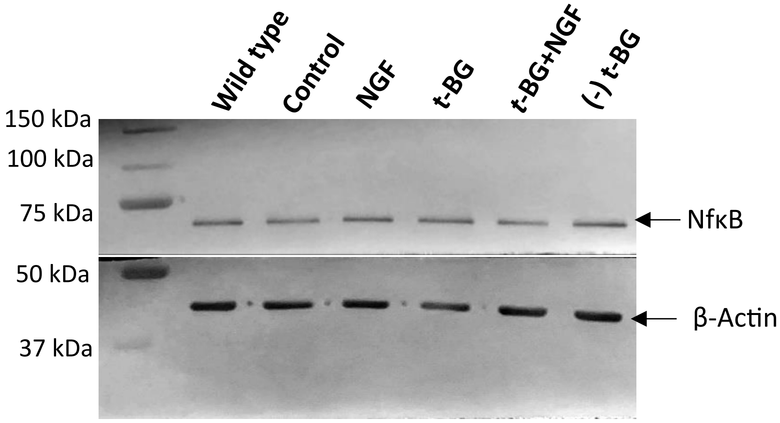


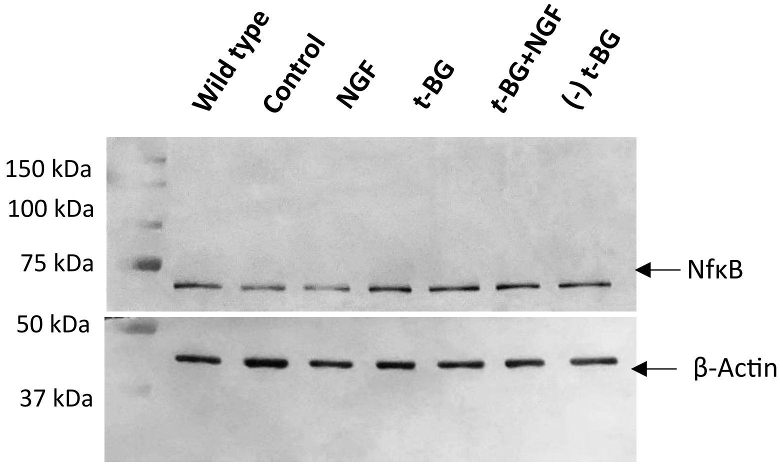


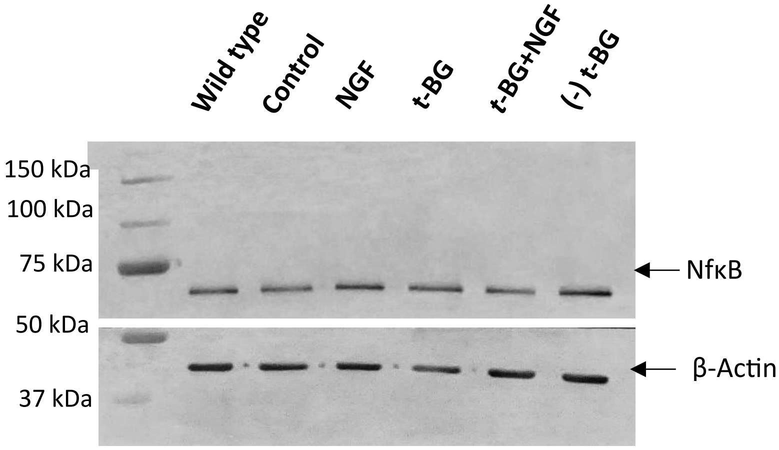


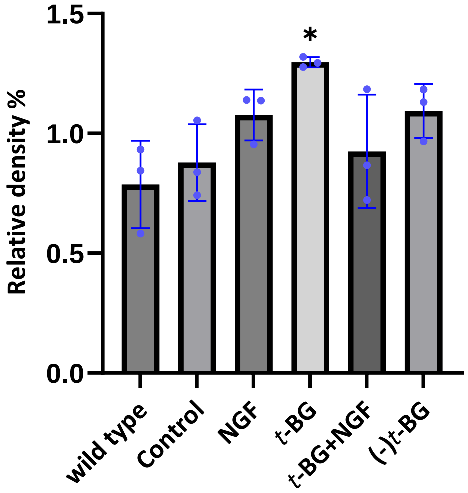

**Table 17: NfκB calculations for the replica *without* type for 48-hour treatment**

| *Sample name* | *NfκB band area* | *Actin band area* | *NfκB area%* | *Actin area%* | *Relative density% (NfκB area%/ Actin area%)* | |
|---|---|---|---|---|---|---|
| *Control* | 20132.18 | 16938.34 | 20.131 | 20.718 | 0.971667 | Replicate 1 |
| *NGF-treated* | 19040.13 | 16361.46 | 19.039 | 20.013 | 0.951332 | |
| *t-BG-treated* | 29079.83 | 19198.68 | 29.077 | 23.483 | 1.238215 | |
| *t-BG+NGF-treated* | 15623.16 | 15223.34 | 15.622 | 18.62 | 0.83899 | |
| *(-) t-BG* | 16132.99 | 14034.25 | 16.132 | 17.166 | 0.939765 | |
| *Control* | 7327.459 | 18835.12 | 12.361 | 17.669 | 0.699587 | Replicate 2 |
| *NGF-treated* | 12583.02 | 21197.74 | 21.227 | 19.886 | 1.067434 | |
| *t-BG-treated* | 14511.65 | 22406.77 | 24.481 | 21.02 | 1.164653 | |
| *t-BG+NGF-treated* | 12360.68 | 22263.62 | 20.852 | 20.885 | 0.99842 | |
| *(-) t-BG* | 12494.87 | 21895.41 | 21.079 | 20.54 | 1.026241 | |
| *Control* | 14042.64 | 28702.77 | 17.336 | 20.569 | 0.842822 | Replicate 3 |
| *NGF-treated* | 17596.89 | 27447.33 | 21.724 | 19.67 | 1.104423 | |
| *t-BG-treated* | 15936.47 | 21129.79 | 19.674 | 15.142 | 1.2993 | |
| *t-BG+NGF-treated* | 12440.74 | 30083.84 | 15.358 | 21.559 | 0.712371 | |
| *(-) t-BG* | 20985.66 | 32178.28 | 25.907 | 23.06 | 1.123461 | |

**Table 18: NfκB calculations for the replica *with* type for 48-hour treatment**

| *Sample name* | *NfκB band area* | *Actin band area* | *NfκB area%* | *Actin area%* | *Relative density% (NfκB area%/ Actin area%)* | |
|---|---|---|---|---|---|---|
| *Wild type* | 18277.89 | 18576.73 | 15.651 | 18.527 | 0.844767 | Replicate 1 |
| *Control* | 20045.84 | 16332.1 | 17.165 | 16.288 | 1.053843 | |
| *NGF-treated* | 18055.23 | 16248.17 | 15.461 | 16.204 | 0.954147 | |
| *t-BG-treated* | 28515.22 | 19183.1 | 24.418 | 19.132 | 1.276291 | |
| *t-BG+NGF-treated* | 15796.16 | 15639 | 13.526 | 15.597 | 0.867218 | |
| *(-) t-BG* | 16090.87 | 14290.49 | 13.779 | 14.252 | 0.966812 | |
| *Wild type* | 8722.338 | 29875.23 | 12.608 | 21.654 | 0.582248 | Replicate 2 |
| *Control* | 7477.581 | 20115.84 | 10.809 | 14.58 | 0.741358 | |
| *NGF-treated* | 12165.29 | 21334.74 | 17.585 | 15.464 | 1.137157 | |
| *t-BG-treated* | 14669.77 | 22599.02 | 21.205 | 16.38 | 1.294567 | |
| *t-BG+NGF-treated* | 13276.95 | 22346.16 | 19.192 | 16.197 | 1.184911 | |
| *(-) t-BG* | 12867.87 | 21693.87 | 18.601 | 15.724 | 1.182969 | |
| *Wild type* | 15913.77 | 30488.13 | 16.341 | 17.501 | 0.933718 | Replicate 3 |
| *Control* | 13959.94 | 29790.66 | 14.335 | 17.101 | 0.838255 | |
| *NGF-treated* | 17787.3 | 27913.57 | 18.265 | 16.023 | 1.139924 | |
| *t-BG-treated* | 16153.94 | 21901.62 | 16.587 | 12.572 | 1.31936 | |
| *t-BG+NGF-treated* | 12267.92 | 30413.97 | 12.597 | 17.459 | 0.721519 | |
| *(-) t-BG* | 21303.49 | 33697.54 | 21.875 | 19.344 | 1.130842 | |

*Pkc western blot replica 48-hour treatment*

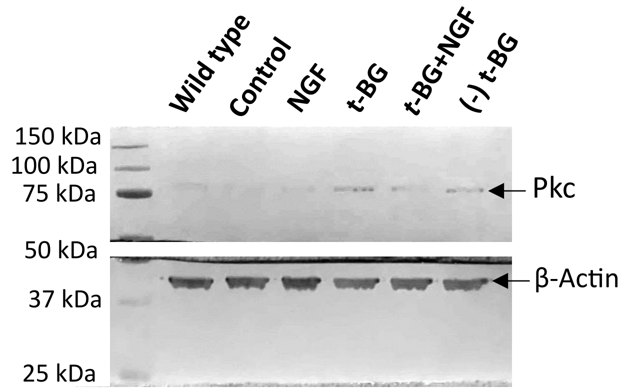


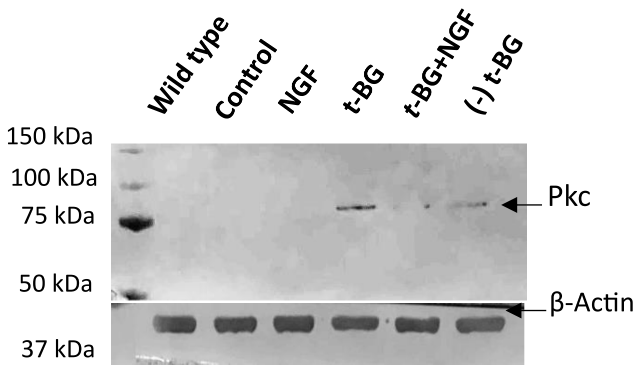


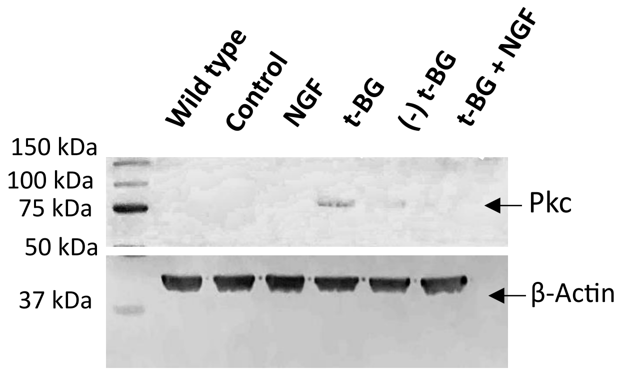


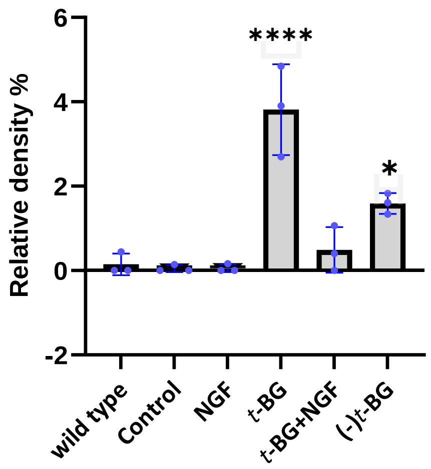

**Table 19: Pkc calculations for the replica *without* wild type for 48-hour treatment**

| *Sample name* | *Pkc band area* | *Actin band area* | *Pkc area%* | *Actin area%* | *Relative density% (Pkc area%/ Actin area%)* | |
|---|---|---|---|---|---|---|
| *Control* | 22.828 | 33546.31 | 0.171 | 21.139 | 0.0080893 | Replicate 1 |
| *NGF-treated* | 70.95 | 33636.36 | 0.532 | 21.196 | 0.025099075 | |
| *t-BG-treated* | 9323.296 | 29282.87 | 69.92 | 18.452 | 3.789291 | |
| *t-BG+NGF-treated* | 1011.184 | 33330.82 | 7.583 | 21.003 | 0.361044 | |
| *(-) t-BG* | 2905.882 | 28899.02 | 21.793 | 18.21 | 1.19676 | |
| *Control* | 263.849 | 27167.77 | 0.171 | 20.526 | 0.0445776 | Replicate 2 |
| *NGF-treated* | 623.991 | 28484.35 | 0.532 | 21.521 | 0.100506482 | |
| *t-BG-treated* | 20219.99 | 26579.87 | 69.92 | 20.082 | 3.49019 | |
| *t-BG+NGF-treated* | 804.891 | 26938.49 | 2.79 | 20.353 | 0.137081 | |
| *(-) t-BG* | 6935.782 | 23185.92 | 24.042 | 17.518 | 1.372417 | |
| *Control* | 818.113 | 23933.28 | 2.986 | 20.301 | 0.1470864 | Replicate 3 |
| *NGF-treated* | 1162.527 | 26948.37 | 4.243 | 22.859 | 0.185616169 | |
| *t-BG-treated* | 12071.85 | 21413.45 | 44.062 | 18.164 | 2.425787 | |
| *t-BG+NGF-treated* | 4884.317 | 22986.97 | 17.828 | 19.498 | 0.91435 | |
| *(-) t-BG* | 8460.61 | 22609.23 | 30.881 | 19.178 | 1.61023 | |

**Table 20: Pkc calculations for the replica *with* wild type for 48-hour treatment**

| *Sample name* | *Pkc band area* | *Actin band area* | *Pkc area%* | *Actin area%* | *Relative density% (Pkc area%/ Actin area%)* | |
|---|---|---|---|---|---|---|
| *Wild type* | 80.536 | 35021.723 | 0.6 | 18.1 | 0.03314 | Replicate 1 |
| *Control* | 22.828 | 33695.551 | 0.17 | 17.415 | 0.00976 | |
| *NGF-treated* | 70.95 | 33352.823 | 0.529 | 17.238 | 0.03068 | |
| *t-BG-treated* | 9323.296 | 29168.045 | 69.501 | 15.075 | 4.61034 | |
| *t-BG+NGF-treated* | 1011.184 | 33138.116 | 7.538 | 17.127 | 0.44012 | |
| *(-) t-BG* | 2905.882 | 29110.258 | 21.662 | 15.045 | 1.43981 | |
| *Wild type* | 262.971 | 24796.401 | 0.903 | 15.702 | 0.05750 | Replicate 2 |
| *Control* | 263.849 | 27346.472 | 0.906 | 17.317 | 0.05231 | |
| *NGF-treated* | 627.163 | 28863.472 | 2.153 | 18.278 | 0.11779 | |
| *t-BG-treated* | 20219.986 | 26513.744 | 69.428 | 16.79 | 4.13508 | |
| *t-BG+NGF-treated* | 813.82 | 27090.492 | 2.794 | 17.155 | 0.16286 | |
| *(-) t-BG* | 6935.782 | 23305.622 | 23.815 | 14.758 | 1.61370 | |
| *Wild type* | 2292.598 | 23790.057 | 7.769 | 17.563 | 0.44235 | Replicate 3 |
| *Control* | 693.82 | 22554.744 | 2.351 | 16.651 | 0.14119 | |
| *NGF-treated* | 903.627 | 25325.836 | 3.062 | 18.697 | 0.16376 | |
| *t-BG-treated* | 11971.853 | 20390.037 | 40.57 | 15.053 | 2.69514 | |
| *t-BG+NGF-treated* | 5086.439 | 21903.137 | 17.237 | 16.17 | 1.06598 | |
| *(-) t-BG* | 8560.581 | 21493.522 | 29.01 | 15.867 | 1.82832 | |

### 6. Western blots for 2-hour treatments

*Akt Western blot replica for 2-hour treatment*

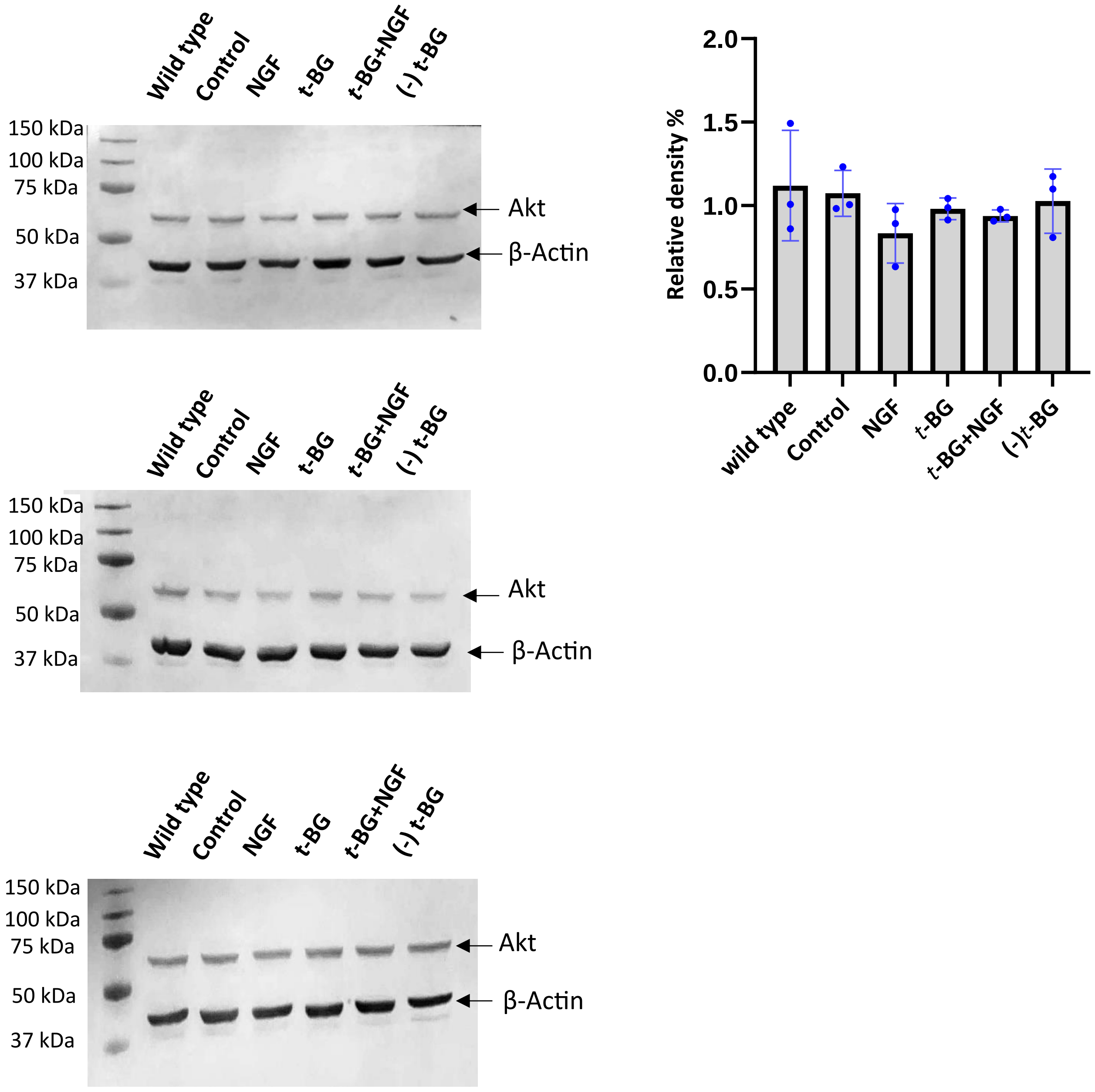

**Tabe 21: Akt calculations for the replica *without* type for 2-hour treatment**

| *Sample name* | *Akt band area* | *Actin band area* | *Akt area%* | *Actin area%* | *Relative density% (Akt area%/ Actin area%)* | |
|---|---|---|---|---|---|---|
| *Control* | 7911.338 | 18835.87 | 23.825 | 20.047 | 1.188457 | Replicate 1 |
| *NGF-treated* | 5210.418 | 17911.84 | 15.691 | 19.064 | 0.82307 | |
| *t-BG-treated* | 7078.489 | 21406.07 | 21.317 | 22.783 | 0.935654 | |
| *t-BG+NGF-treated* | 6120.782 | 18541.29 | 18.433 | 19.734 | 0.934073 | |
| *(-) t-BG* | 6884.681 | 17261.68 | 20.733 | 18.372 | 1.128511 | |
| *Control* | 3635.296 | 20000.02 | 22.362 | 20.452 | 1.093389 | Replicate 2 |
| *NGF-treated* | 2633.811 | 20345.05 | 16.201 | 20.804 | 0.778744 | |
| *t-BG-treated* | 4280.832 | 21444.17 | 26.333 | 21.928 | 1.200885 | |
| *t-BG+NGF-treated* | 3222.64 | 19287.73 | 19.823 | 19.723 | 1.00507 | |
| *(-) t-BG* | 2484.104 | 16714.73 | 15.281 | 17.092 | 0.894044 | |
| *Control* | 6556.66 | 15845 | 17.76 | 17.795 | 0.998033 | Replicate 3 |
| *NGF-treated* | 6679.075 | 16438.63 | 18.092 | 18.461 | 0.980012 | |
| *t-BG-treated* | 8000.731 | 18631.41 | 21.672 | 20.924 | 1.035748 | |
| *t-BG+NGF-treated* | 7676.246 | 19661.87 | 20.793 | 22.081 | 0.941669 | |
| *(-) t-BG* | 8004.539 | 18467.27 | 21.682 | 20.739 | 1.04547 | |

**Table 22: Akt calculations for the replica *with* type for 2-hour treatment**

| *Sample name* | *Akt band area* | *Actin band area* | *Akt area%* | *Actin area%* | *Relative density% (Akt area%/ Actin area%)* | |
|---|---|---|---|---|---|---|
| *Wild type* | 6629.317 | 22757.7 | 16.776 | 19.505 | 0.860087 | Replicate 1 |
| *Control* | 7874.167 | 18865.87 | 19.926 | 16.17 | 1.232282 | |
| *NGF-treated* | 5407.66 | 17877.02 | 13.685 | 15.322 | 0.89316 | |
| *t-BG-treated* | 6664.832 | 21534.55 | 16.866 | 18.457 | 0.9138 | |
| *t-BG+NGF-treated* | 6100.196 | 18429.7 | 15.437 | 15.796 | 0.977273 | |
| *(-) t-BG* | 6840.146 | 17209.97 | 17.31 | 14.75 | 1.173559 | |
| *Wild type* | 6868.288 | 23655.41 | 29.479 | 19.743 | 1.493137 | Replicate 2 |
| *Control* | 3790.711 | 19826.84 | 16.27 | 16.548 | 0.9832 | |
| *NGF-treated* | 2473.569 | 20059.34 | 10.617 | 16.742 | 0.634154 | |
| *t-BG-treated* | 4243.711 | 20940.63 | 18.214 | 17.477 | 1.04217 | |
| *t-BG+NGF-treated* | 3360.761 | 19050.97 | 14.425 | 15.9 | 0.907233 | |
| *(-) t-BG* | 2561.933 | 16283.02 | 10.996 | 13.59 | 0.809124 | |
| *Wild type* | 6364.296 | 15739.77 | 15.172 | 15.07 | 1.006768 | Replicate 3 |
| *Control* | 6391.711 | 15818 | 15.237 | 15.145 | 1.006075 | |
| *NGF-treated* | 6420.539 | 16394.22 | 15.306 | 15.697 | 0.975091 | |
| *t-BG-treated* | 7405.903 | 18687.12 | 17.655 | 17.892 | 0.986754 | |
| *t-BG+NGF-treated* | 7275.711 | 19473.05 | 17.345 | 18.645 | 0.930276 | |
| *(-) t-BG* | 8089.246 | 18331.15 | 19.284 | 17.551 | 1.098741 | |

*Catalase Western blot replica for 2-hour treatment*

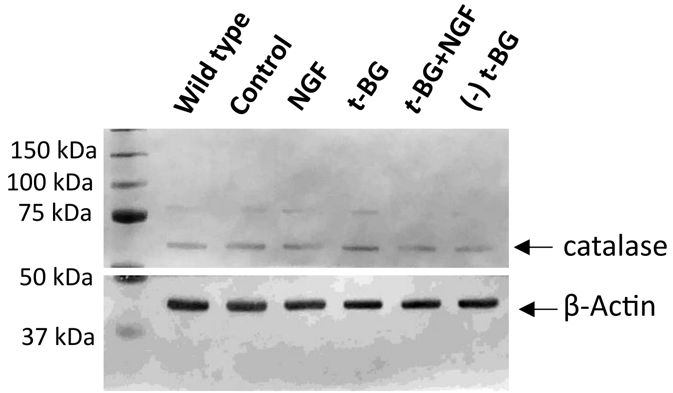


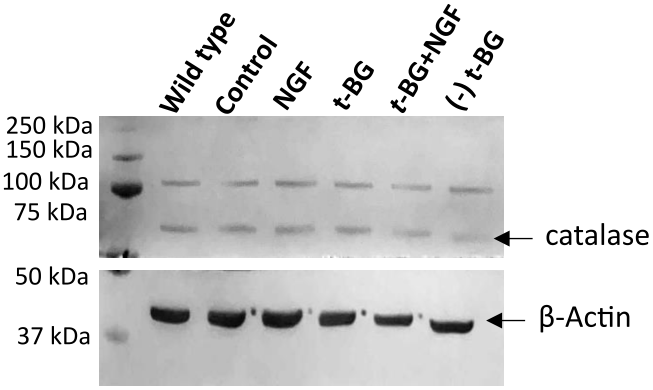


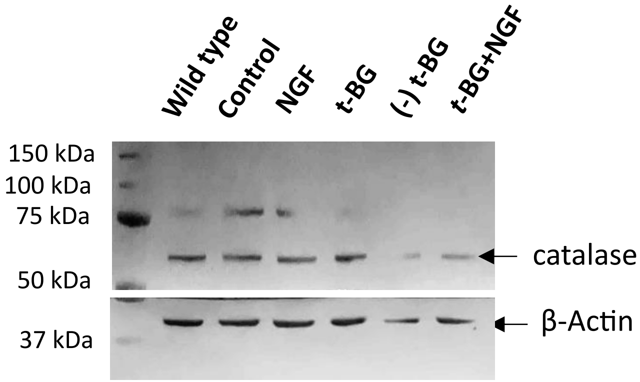


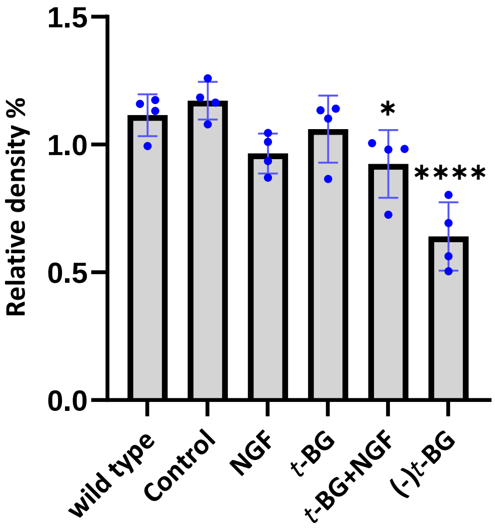


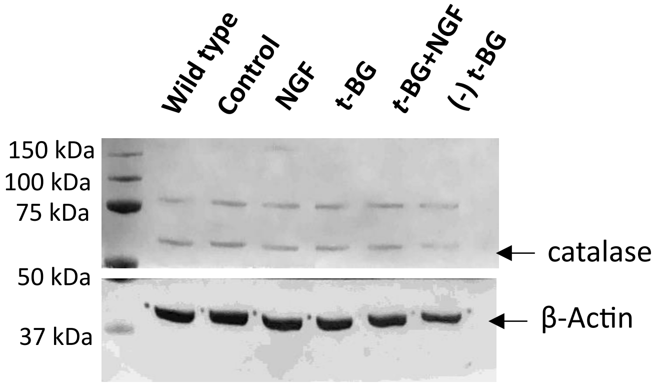

**Table 23: Catalase calculations for the replica *without* type for 2-hour treatment**

| *Sample name* | *Catalase band area* | *Actin band area* | *Catalase area%* | *Actin area%* | *Relative density% (Catalase area%/ Actin area%)* | |
|---|---|---|---|---|---|---|
| *Control* | 9419.673 | 18345.43 | 27.058 | 25.156 | 1.075608 | Replicate 1 |
| *NGF-treated* | 8967.238 | 18239.75 | 25.759 | 25.011 | 1.029907 | |
| *t-BG-treated* | 9032.48 | 17224.36 | 25.946 | 23.618 | 1.098569 | |
| *t-BG+NGF-treated* | 5351.459 | 11784.46 | 15.372 | 16.159 | 0.951296 | |
| *(-) t-BG* | 2041.619 | 7333.66 | 5.865 | 10.056 | 0.583234 | |
| *Control* | 19005.25 | 23994.94 | 28.001 | 21.225 | 1.319246 | Replicate 2 |
| *NGF-treated* | 14313.28 | 23232.21 | 21.088 | 20.551 | 1.02613 | |
| *t-BG-treated* | 15410.24 | 21879.7 | 22.704 | 19.354 | 1.173091 | |
| *t-BG+NGF-treated* | 10136.7 | 21913.84 | 14.935 | 19.384 | 0.770481 | |
| *(-) t-BG* | 9007.652 | 22027.97 | 13.271 | 19.485 | 0.681088 | |
| *Control* | 12103.31 | 44652.51 | 30.06 | 25.289 | 1.188659 | Replicate 3 |
| *NGF-treated* | 7800.731 | 37782.56 | 19.374 | 21.398 | 0.905412 | |
| *t-BG-treated* | 6854.539 | 33537.47 | 17.024 | 18.994 | 0.896283 | |
| *t-BG+NGF-treated* | 8459.024 | 34686.81 | 21.009 | 19.645 | 1.069432 | |
| *(-) t-BG* | 5046.731 | 25911.94 | 12.534 | 14.675 | 0.854106 | |
| *Control* | 19416.44 | 32098.58 | 24.739 | 20.905 | 1.183401 | Replicate 4 |
| *NGF-treated* | 20475.06 | 37058.71 | 26.087 | 24.135 | 1.080878 | |
| *t-BG-treated* | 16779.04 | 28953.25 | 21.378 | 18.856 | 1.133751 | |
| *t-BG+NGF-treated* | 12193.74 | 24623.4 | 15.536 | 16.036 | 0.96882 | |
| *(-) t-BG* | 9622.016 | 30813.28 | 12.259 | 20.068 | 0.610873 | |

**Table 24: Catalase calculations for the replica *with* type for 2-hour treatment**

| *Sample name* | *Catalase band area* | *Actin band area* | *Catalase area%* | *Actin area%* | *Relative density% (Catalase area%/ Actin area%)* | |
|---|---|---|---|---|---|---|
| *Wild type* | 7946.187 | 18423.55 | 20.113 | 20.216 | 0.994905 | Replicate 1 |
| *Control* | 8516.602 | 18198.14 | 21.557 | 19.969 | 1.079523 | |
| *NGF-treated* | 8303.53 | 18323.17 | 21.018 | 20.106 | 1.04536 | |
| *t-BG-treated* | 8151.53 | 17061.36 | 20.633 | 18.722 | 1.102072 | |
| *t-BG+NGF-treated* | 4952.045 | 11646.87 | 12.535 | 12.78 | 0.980829 | |
| *(-) t-BG* | 1637.154 | 7479.196 | 4.144 | 8.207 | 0.504935 | |
| *Wild type* | 21185.59 | 28382.54 | 23.533 | 20.041 | 1.174243 | Replicate 2 |
| *Control* | 19351.03 | 24180.35 | 21.495 | 17.074 | 1.258932 | |
| *NGF-treated* | 13797.57 | 23212.09 | 15.326 | 16.39 | 0.935082 | |
| *t-BG-treated* | 15817.84 | 21808 | 17.57 | 15.399 | 1.140983 | |
| *t-BG+NGF-treated* | 10217.53 | 22136.21 | 11.35 | 15.631 | 0.726121 | |
| *(-) t-BG* | 9655.844 | 21900.26 | 10.726 | 15.464 | 0.693611 | |
| *Wild type* | 12426.55 | 40148.64 | 22.048 | 18.411 | 1.197545 | Replicate 3 |
| *Control* | 12949.14 | 44312.56 | 22.975 | 20.32 | 1.130659 | |
| *NGF-treated* | 8260.731 | 38223.93 | 14.657 | 17.528 | 0.836205 | |
| *t-BG-treated* | 7380.53 | 33875.25 | 13.095 | 15.534 | 0.84299 | |
| *t-BG+NGF-treated* | 9661.945 | 34941.22 | 17.143 | 16.023 | 1.0699 | |
| *(-) t-BG* | 5681.974 | 26571.35 | 10.081 | 12.185 | 0.827329 | |
| *Wild type* | 17745.74 | 30997.32 | 19.098 | 16.867 | 1.13227 | Replicate 4 |
| *Control* | 18947.71 | 32185.58 | 20.392 | 17.514 | 1.164326 | |
| *NGF-treated* | 18795.79 | 36794.58 | 20.228 | 20.022 | 1.010289 | |
| *t-BG-treated* | 16775.04 | 29236.25 | 18.053 | 15.909 | 1.134766 | |
| *t-BG+NGF-treated* | 11994.5 | 24126.45 | 12.909 | 13.128 | 0.983318 | |
| *(-) t-BG* | 8660.409 | 30432.04 | 9.32 | 16.56 | 0.562802 | |

*Cfos1 Western blot replica for 2-hour treatment*

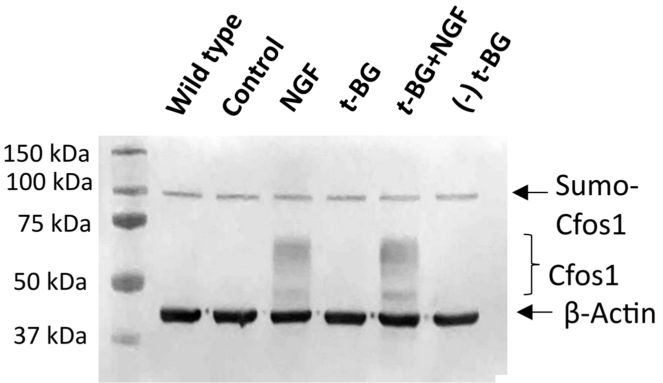


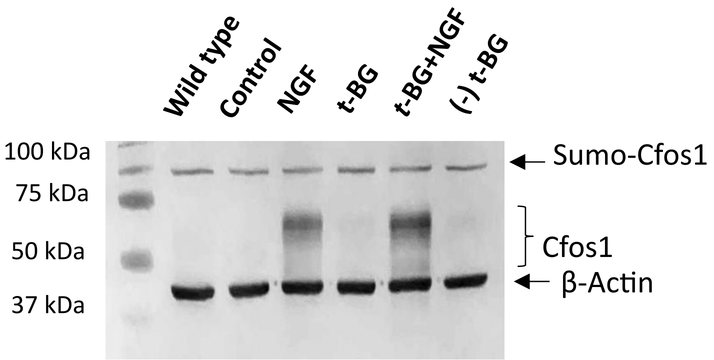


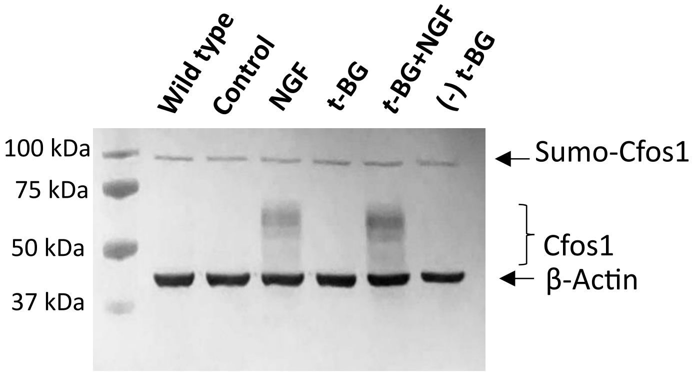


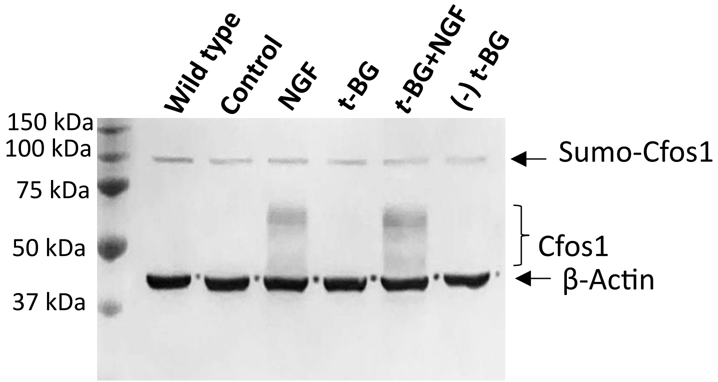


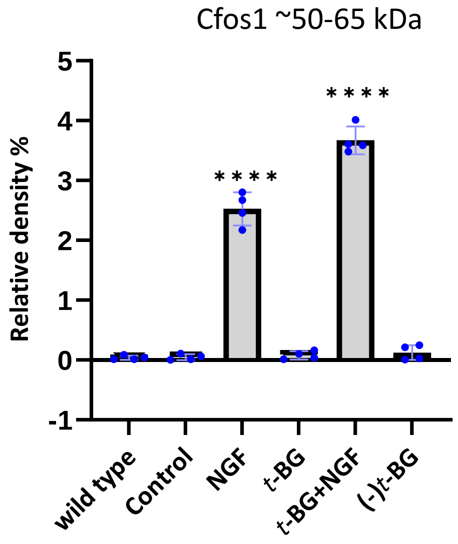


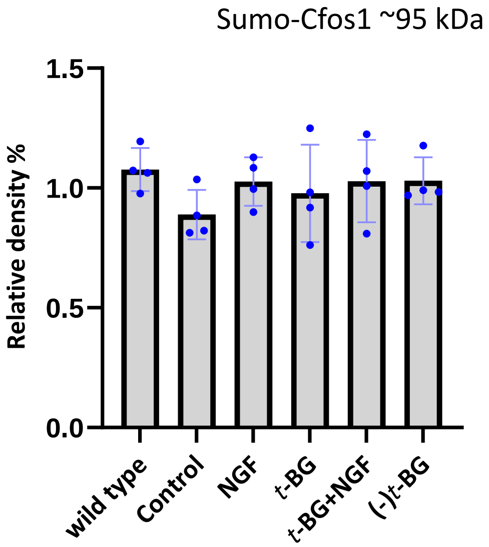

**Table 25: Sumo-Cfos 1 (~95 kDa ) calculations for the replica *without* type for 2-hour treatment**

| *Sample name* | *Sumo-Cfos 1 band area* | *Actin band area* | *Sumo-Cfos 1 area%* | *Actin area%* | *Relative density% (Sumo-Cfos 1 area%/ Actin area%)* | |
|---|---|---|---|---|---|---|
| *Control* | 1499.406 | 13926.2 | 23.769 | 22.643 | 1.049728 | Replicate 1 |
| *NGF-treated* | 1277.82 | 10896.4 | 20.257 | 17.716 | 1.14343 | |
| *t-BG-treated* | 1311.991 | 12848.76 | 20.798 | 20.891 | 0.995548 | |
| *t-BG+NGF-treated* | 1017.527 | 12083.52 | 16.13 | 19.647 | 0.82099 | |
| *(-) t-BG* | 1201.406 | 11749.59 | 19.045 | 19.104 | 0.996912 | |
| *Control* | 988.113 | 12926.71 | 18.868 | 22.733 | 0.829983 | Replicate 2 |
| *NGF-treated* | 1038.113 | 11091.57 | 19.823 | 19.505 | 1.016304 | |
| *t-BG-treated* | 1078.577 | 12379.4 | 20.595 | 21.77 | 0.946027 | |
| *t-BG+NGF-treated* | 1113.527 | 11138.69 | 21.263 | 19.588 | 1.085512 | |
| *(-) t-BG* | 1018.648 | 9327.619 | 19.451 | 16.403 | 1.18582 | |
| *Control* | 1737.012 | 11422.95 | 17.918 | 20.487 | 0.874603 | Replicate 3 |
| *NGF-treated* | 1776.184 | 11398.8 | 18.323 | 20.444 | 0.896253 | |
| *t-BG-treated* | 2436.598 | 11245.78 | 25.135 | 20.169 | 1.246219 | |
| *t-BG+NGF-treated* | 1739.841 | 10023.08 | 17.948 | 17.976 | 0.998442 | |
| *(-) t-BG* | 2004.355 | 11666.61 | 20.676 | 20.924 | 0.988148 | |
| *Control* | 822.749 | 12844.51 | 19.798 | 23.297 | 0.849809 | Replicate 4 |
| *NGF-treated* | 993.941 | 11616.44 | 23.918 | 21.07 | 1.135168 | |
| *t-BG-treated* | 689.335 | 11496.9 | 16.588 | 20.853 | 0.795473 | |
| *t-BG+NGF-treated* | 889.062 | 9215.861 | 21.394 | 16.716 | 1.279852 | |
| *(-) t-BG* | 760.577 | 9959.518 | 18.302 | 18.064 | 1.013175 | |

**Table 26: Sumo-Cfos 1 (~95 kDa ) calculations for the replica *with* type for 2-hour treatment**

| *Sample name* | *Sumo-Cfos 1 band area* | *Actin band area* | *Sumo-Cfos 1 area%* | *Actin area%* | *Relative density% (Sumo-Cfos 1 area%/ Actin area%)* | |
|---|---|---|---|---|---|---|
| *Wild type* | 1486.234 | 13445.54 | 19.068 | 17.939 | 1.062936 | Replicate 1 |
| *Control* | 1499.406 | 13926.2 | 19.237 | 18.581 | 1.035305 | |
| *NGF-treated* | 1277.82 | 10896.4 | 16.394 | 14.538 | 1.127665 | |
| *t-BG-treated* | 1311.991 | 12848.76 | 16.832 | 17.143 | 0.981858 | |
| *t-BG+NGF-treated* | 1017.698 | 12083.52 | 13.057 | 16.122 | 0.809887 | |
| *(-) t-BG* | 1201.406 | 11749.59 | 15.413 | 15.677 | 0.98316 | |
| *Wild type* | 1290.406 | 12671.83 | 19.564 | 18.223 | 1.073588 | Replicate 2 |
| *Control* | 1007.335 | 12926.71 | 15.272 | 18.59 | 0.821517 | |
| *NGF-treated* | 1047.527 | 11091.57 | 15.882 | 15.951 | 0.995674 | |
| *t-BG-treated* | 1078.577 | 12379.4 | 16.352 | 17.803 | 0.918497 | |
| *t-BG+NGF-treated* | 1130.749 | 11138.69 | 17.143 | 16.019 | 1.070167 | |
| *(-) t-BG* | 1041.234 | 9327.619 | 15.786 | 13.414 | 1.17683 | |
| *Wild type* | 2464.062 | 14541.24 | 20.209 | 20.685 | 0.976988 | Replicate 3 |
| *Control* | 1754.941 | 11422.95 | 14.393 | 16.249 | 0.885778 | |
| *NGF-treated* | 1779.184 | 11398.8 | 14.592 | 16.215 | 0.899907 | |
| *t-BG-treated* | 2436.598 | 11245.78 | 19.984 | 15.997 | 1.249234 | |
| *t-BG+NGF-treated* | 1753.527 | 10023.08 | 14.382 | 14.258 | 1.008697 | |
| *(-) t-BG* | 2004.355 | 11666.61 | 16.439 | 16.596 | 0.99054 | |
| *Wild type* | 1151.627 | 12238.97 | 21.705 | 18.166 | 1.194814 | Replicate 4 |
| *Control* | 822.749 | 12844.51 | 15.506 | 19.065 | 0.813323 | |
| *NGF-treated* | 992.527 | 11616.44 | 18.706 | 17.242 | 1.084909 | |
| *t-BG-treated* | 689.335 | 11496.9 | 12.992 | 17.065 | 0.761324 | |
| *t-BG+NGF-treated* | 889.062 | 9215.861 | 16.756 | 13.679 | 1.224943 | |
| *(-) t-BG* | 760.577 | 9959.518 | 14.335 | 14.783 | 0.969695 | |

**Table 27: Cfos1 (~50-65 kDa) calculations for the replica *without* type for 2-hour treatment**

| *Sample name* | *Cfos 1 band area* | *Actin band area* | *Cfos 1 area%* | *Actin area%* | *Relative density% (Cfos 1 area%/ Actin area%)* | |
|---|---|---|---|---|---|---|
| *Control* | 6.121 | 13926.2 | 0.043 | 22.643 | 0.001899 | Replicate 1 |
| *NGF-treated* | 5757.815 | 10896.4 | 40.912 | 17.716 | 2.309325 | |
| *t-BG-treated* | 90.364 | 12848.76 | 0.642 | 20.891 | 0.030731 | |
| *t-BG+NGF-treated* | 8199.614 | 12083.52 | 58.263 | 19.647 | 2.96549 | |
| *(-) t-BG* | 19.536 | 11749.59 | 0.139 | 19.104 | 0.007276 | |
| *Control* | 40.121 | 12926.71 | 0.25 | 22.733 | 0.010997 | Replicate 2 |
| *NGF-treated* | 6910.53 | 11091.57 | 42.995 | 19.505 | 2.204307 | |
| *t-BG-treated* | 30.121 | 12379.4 | 0.187 | 21.77 | 0.00859 | |
| *t-BG+NGF-treated* | 9033.451 | 11138.69 | 56.203 | 19.588 | 2.869257 | |
| *(-) t-BG* | 58.778 | 9327.619 | 0.366 | 16.403 | 0.022313 | |
| *Control* | 263.192 | 11422.95 | 1.057 | 20.487 | 0.051594 | Replicate 3 |
| *NGF-treated* | 10112.43 | 11398.8 | 40.605 | 20.444 | 1.986157 | |
| *t-BG-treated* | 661.506 | 11245.78 | 2.656 | 20.169 | 0.131687 | |
| *t-BG+NGF-treated* | 12972.69 | 10023.08 | 52.09 | 17.976 | 2.897753 | |
| *(-) t-BG* | 894.406 | 11666.61 | 3.591 | 20.924 | 0.171621 | |
| *Control* | 212.192 | 12877.56 | 2.062 | 23.11 | 0.089225 | Replicate 4 |
| *NGF-treated* | 3872.66 | 12383.1 | 37.632 | 22.223 | 1.693381 | |
| *t-BG-treated* | 183.364 | 11271.95 | 1.782 | 20.229 | 0.088091 | |
| *t-BG+NGF-treated* | 5641.593 | 9314.861 | 54.821 | 16.717 | 3.279356 | |
| *(-) t-BG* | 381.092 | 9874.861 | 3.703 | 17.722 | 0.208949 | |

**Table 28: Cfos1 (~50-65 kDa) calculations for the replica *with* type for 2-hour treatment**

| *Sample name* | *Cfos 1 band area* | *Actin band area* | *Cfos 1 area%* | *Actin area%* | *Relative density% (Cfos 1 area%/ Actin area%)* | |
|---|---|---|---|---|---|---|
| *Wild type* | 37.95 | 13445.54 | 0.268 | 17.939 | 0.01494 | Replicate 1 |
| *Control* | 6.121 | 13926.2 | 0.043 | 18.581 | 0.002314 | |
| *NGF-treated* | 5761.187 | 10896.4 | 40.757 | 14.538 | 2.803481 | |
| *t-BG-treated* | 90.364 | 12848.76 | 0.639 | 17.143 | 0.037275 | |
| *t-BG+NGF-treated* | 8219.886 | 12083.52 | 58.151 | 16.122 | 3.606935 | |
| *(-) t-BG* | 19.828 | 11749.59 | 0.14 | 15.677 | 0.00893 | |
| *Wild type* | 130.95 | 12671.83 | 0.808151 | 18.223 | 0.044348 | Replicate 2 |
| *Control* | 40.121 | 12926.71 | 0.247605 | 18.59 | 0.013319 | |
| *NGF-treated* | 6910.53 | 11091.57 | 42.64796 | 15.951 | 2.673686 | |
| *t-BG-treated* | 30.121 | 12379.4 | 0.18589 | 17.803 | 0.010442 | |
| *t-BG+NGF-treated* | 9033.158 | 11138.69 | 55.74765 | 16.019 | 3.480095 | |
| *(-) t-BG* | 58.778 | 9327.619 | 0.362745 | 13.414 | 0.02704 | |
| *Wild type* | 458.021 | 14541.24 | 1.805 | 20.685 | 0.087261 | Replicate 3 |
| *Control* | 263.192 | 11422.95 | 1.037 | 16.249 | 0.063819 | |
| *NGF-treated* | 10112.43 | 11398.8 | 39.852 | 16.215 | 2.457724 | |
| *t-BG-treated* | 666.678 | 11245.78 | 2.627 | 15.997 | 0.164218 | |
| *t-BG+NGF-treated* | 12972.69 | 10023.08 | 51.124 | 14.258 | 3.585636 | |
| *(-) t-BG* | 901.92 | 11666.61 | 3.554 | 16.596 | 0.214148 | |
| *Wild type* | 22.121 | 12238.97 | 0.214 | 18.166 | 0.01178 | Replicate 4 |
| *Control* | 212.192 | 12844.51 | 2.05 | 19.065 | 0.107527 | |
| *NGF-treated* | 3872.368 | 11616.44 | 37.418 | 17.242 | 2.170166 | |
| *t-BG-treated* | 183.364 | 11496.9 | 1.772 | 17.065 | 0.103838 | |
| *t-BG+NGF-treated* | 5677.844 | 9215.861 | 54.864 | 13.679 | 4.01082 | |
| *(-) t-BG* | 381.092 | 9959.518 | 3.682 | 14.783 | 0.24907 | |

*Erk1/2 Western blot replica for 2-hour treatment*

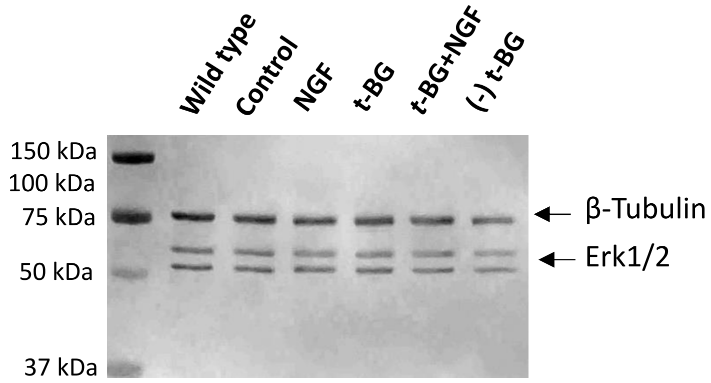


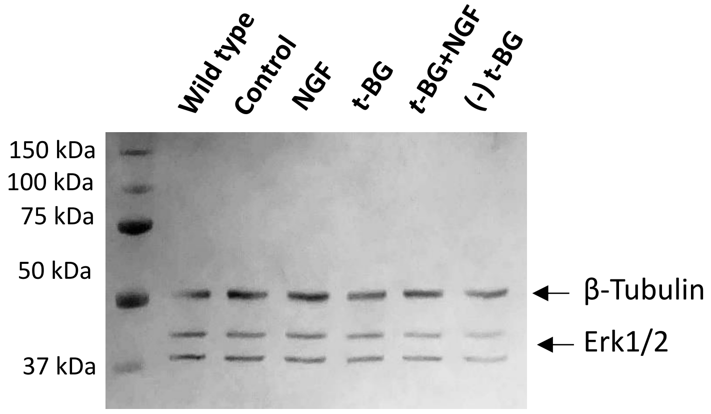


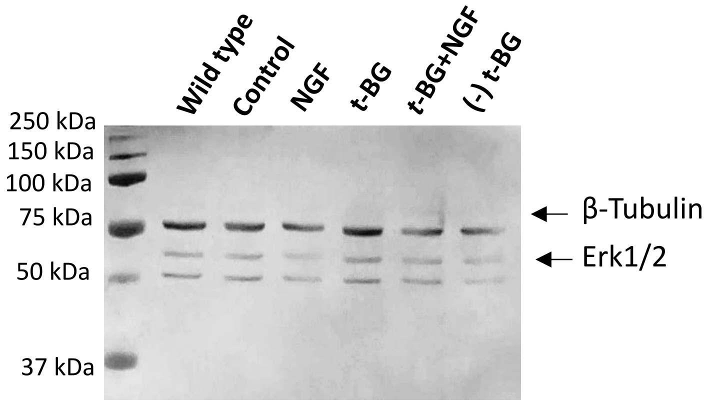


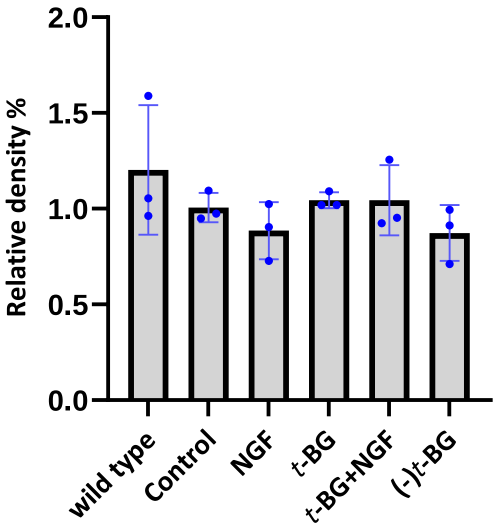

**Table 29: Erk1/2 calculations for the replica *without* type for 2-hour treatment**

| *Sample name* | *Erk1/2 band area* | *Tubulin band area* | *Erk1/2 area%* | *Tubulin area%* | *Relative density% (Erk1/2 area%/ Tubulin area%)* | |
|---|---|---|---|---|---|---|
| *Control* | 9447.287 | 11932.9 | 24.135 | 22.644 | 1.065845 | Replicate 1 |
| *NGF-treated* | 10096.12 | 13710.92 | 25.793 | 26.018 | 0.991352 | |
| *t-BG-treated* | 8309.602 | 9384.146 | 21.229 | 17.807 | 1.192172 | |
| *t-BG+NGF-treated* | 6675.388 | 9680.004 | 17.053 | 18.369 | 0.928358 | |
| *(-) t-BG* | 4615.459 | 7990.418 | 11.791 | 15.163 | 0.777617 | |
| *Control* | 11392.26 | 14796.05 | 24.457 | 25.775 | 0.948865 | Replicate 2 |
| *NGF-treated* | 10281.6 | 12330.61 | 22.073 | 21.48 | 1.027607 | |
| *t-BG-treated* | 9721.773 | 11654.37 | 20.871 | 20.302 | 1.028027 | |
| *t-BG+NGF-treated* | 8681.652 | 10723.66 | 18.638 | 18.681 | 0.997698 | |
| *(-) t-BG* | 6503.309 | 7899.66 | 13.961 | 13.761 | 1.014534 | |
| *Control* | 4370.317 | 9667.518 | 23.821 | 23.976 | 0.993535 | Replicate 3 |
| *NGF-treated* | 2394.539 | 6822.326 | 13.052 | 16.92 | 0.771395 | |
| *t-BG-treated* | 5323.61 | 11440.28 | 29.017 | 28.372 | 1.022734 | |
| *t-BG+NGF-treated* | 3650.246 | 6751.69 | 19.896 | 16.744 | 1.188247 | |
| *(-) t-BG* | 2608.025 | 5640.397 | 14.215 | 13.988 | 1.016228 | |

**Table 30: Erk1/2 calculations for the replica *with* type for 2-hour treatment**

| *Sample name* | *Erk1/2 band area* | *Tubulin band area* | *Erk1/2 area%* | *Tubulin area%* | *Relative density% (Erk1/2 area%/ Tubulin area%)* | |
|---|---|---|---|---|---|---|
| *Wild type* | 7817.803 | 5971.761 | 17.321 | 10.899 | 1.589228 | Replicate 1 |
| *Control* | 9020.288 | 11243.2 | 19.984 | 20.519 | 0.973927 | |
| *NGF-treated* | 9489.459 | 12737.56 | 21.024 | 23.246 | 0.904414 | |
| *t-BG-treated* | 7772.045 | 8644.024 | 17.219 | 15.776 | 1.091468 | |
| *t-BG+NGF-treated* | 6711.238 | 8815.74 | 14.868 | 16.089 | 0.92411 | |
| *(-) t-BG* | 4325.995 | 7381.225 | 9.584 | 13.471 | 0.711454 | |
| *Wild type* | 11545.82 | 13538.83 | 21.046 | 19.959 | 1.054462 | Replicate 2 |
| *Control* | 10629.02 | 13838.8 | 19.375 | 20.401 | 0.949708 | |
| *NGF-treated* | 9512.187 | 11488.78 | 17.339 | 16.937 | 1.023735 | |
| *t-BG-treated* | 9100.673 | 11034.66 | 16.588 | 16.267 | 1.019733 | |
| *t-BG+NGF-treated* | 7999.531 | 10378.08 | 14.582 | 15.299 | 0.953134 | |
| *(-) t-BG* | 6073.601 | 7554.246 | 11.071 | 11.136 | 0.994163 | |
| *Wild type* | 4464.61 | 10964.52 | 22.301 | 23.175 | 0.962287 | Replicate 3 |
| *Control* | 4031.904 | 8714.104 | 20.14 | 18.418 | 1.093495 | |
| *NGF-treated* | 1920.518 | 6244.326 | 9.593 | 13.198 | 0.726853 | |
| *t-BG-treated* | 4390.954 | 10184.33 | 21.933 | 21.526 | 1.018907 | |
| *t-BG+NGF-treated* | 3232.883 | 6082.569 | 16.148 | 12.856 | 1.256067 | |
| *(-) t-BG* | 1978.883 | 5122.104 | 9.884 | 10.826 | 0.912987 | |

*Ferritin Western blot replica for 2-hour treatment*

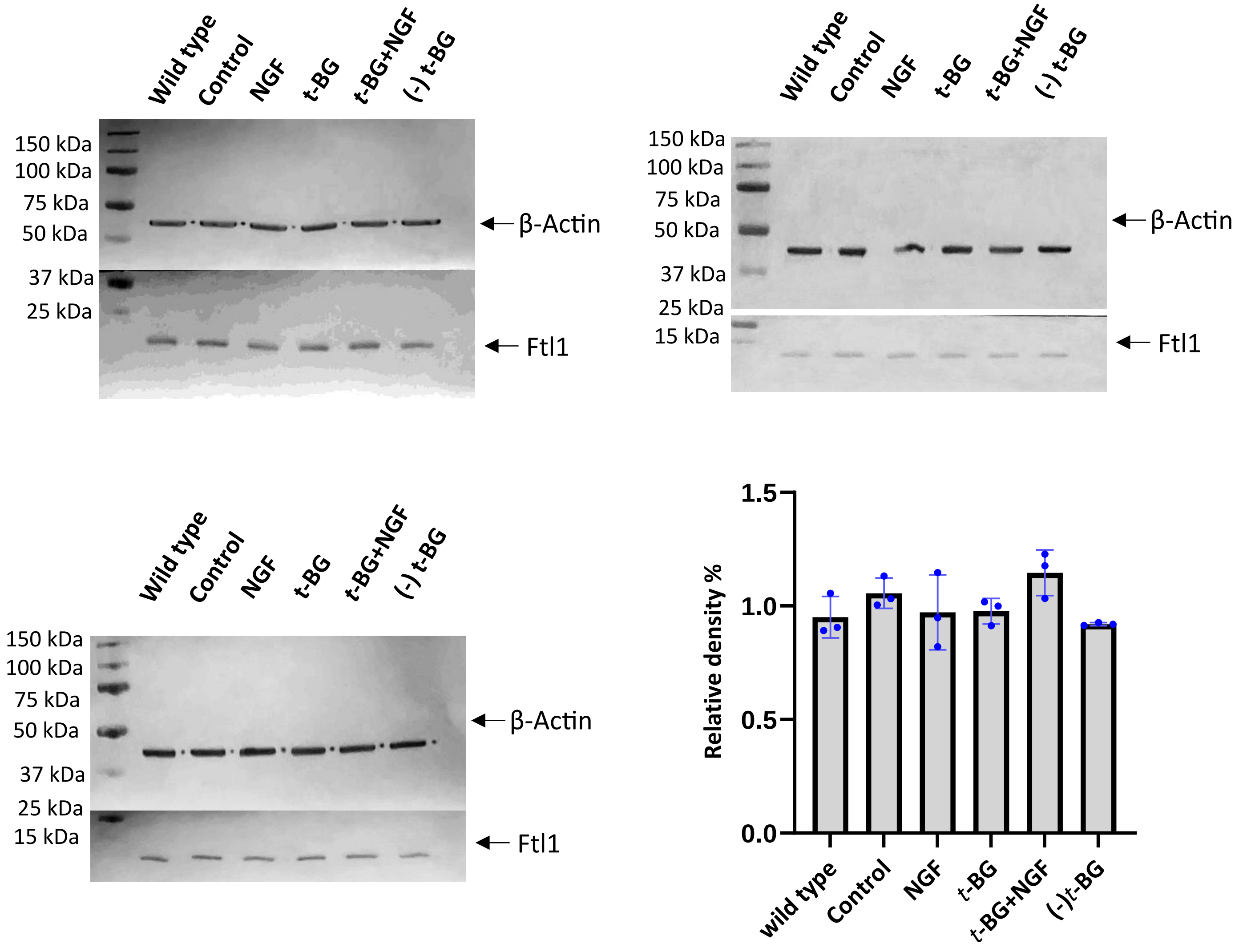

**Table 31: Ferritin calculations for the replica *without* type for 2-hour treatment**

| *Sample name* | *Ftl1 band area* | *Actin band area* | *Ftl1 area%* | *Actin area%* | *Relative density% (Ftl1 area%/ Actin area%)* | |
|---|---|---|---|---|---|---|
| *Control* | 22700.38 | 14565.17 | 18.748 | 18.246 | 1.027513 | Replicate 1 |
| *NGF-treated* | 21140.71 | 16750.65 | 17.46 | 20.983 | 0.832102 | |
| *t-BG-treated* | 27010.18 | 17493.36 | 22.308 | 21.914 | 1.017979 | |
| *t-BG+NGF-treated* | 29030.44 | 16195.05 | 23.976 | 20.287 | 1.181841 | |
| *(-) t-BG* | 21198.18 | 14824.27 | 17.508 | 18.57 | 0.942811 | |
| *Control* | 24809.99 | 14825.46 | 23.437 | 21.705 | 1.079797 | Replicate 2 |
| *NGF-treated* | 19683.25 | 11393.07 | 18.594 | 16.68 | 1.114748 | |
| *t-BG-treated* | 19889.22 | 14557.22 | 18.789 | 21.313 | 0.881575 | |
| *t-BG+NGF-treated* | 19672.84 | 12232.85 | 18.584 | 17.91 | 1.037633 | |
| *(-) t-BG* | 21801.59 | 15294.68 | 20.595 | 22.392 | 0.919748 | |
| *Control* | 21244.09 | 22566.48 | 20.462 | 20.413 | 1.0024 | Replicate 3 |
| *NGF-treated* | 21707.06 | 25096.67 | 20.908 | 22.702 | 0.920976 | |
| *t-BG-treated* | 21758.11 | 23358.62 | 20.957 | 21.13 | 0.991813 | |
| *t-BG+NGF-treated* | 22297.4 | 20502.67 | 21.477 | 18.547 | 1.157977 | |
| *(-) t-BG* | 16813.82 | 19022.95 | 16.195 | 17.208 | 0.941132 | |

**Table 32: Ferritin calculations for the replica *with* type for 2-hour treatment**

| *Sample name* | *Ftl1 band area* | *Actin band area* | *Ftl1 area%* | *Actin area%* | *Relative density% (Ftl1 area%/ Actin area%)* | |
|---|---|---|---|---|---|---|
| *Wild type* | 22968.25 | 14045.85 | 15.905 | 15.068 | 1.055548 | Replicate 1 |
| *Control* | 22717.97 | 14213.51 | 15.732 | 15.248 | 1.031742 | |
| *NGF-treated* | 21046.01 | 16565.24 | 14.574 | 17.77 | 0.820146 | |
| *t-BG-treated* | 27062.89 | 17481.19 | 18.74 | 18.753 | 0.999307 | |
| *t-BG+NGF-treated* | 29370.86 | 16111.75 | 20.339 | 17.284 | 1.176753 | |
| *(-) t-BG* | 21243.47 | 14800.27 | 14.711 | 15.877 | 0.92656 | |
| *Wild type* | 20861.25 | 15417.15 | 16.482 | 18.211 | 0.905057 | Replicate 2 |
| *Control* | 25141.99 | 14859.34 | 19.864 | 17.552 | 1.131723 | |
| *NGF-treated* | 19784.13 | 11538.95 | 15.631 | 13.63 | 1.146809 | |
| *t-BG-treated* | 20006.69 | 14655.68 | 15.807 | 17.311 | 0.913119 | |
| *t-BG+NGF-treated* | 19493.47 | 12613.92 | 15.401 | 14.9 | 1.033624 | |
| *(-) t-BG* | 21282.06 | 15574.22 | 16.815 | 18.396 | 0.914057 | |
| *Wild type* | 18450.89 | 22213.33 | 14.908 | 16.722 | 0.89152 | Replicate 3 |
| *Control* | 21231.55 | 22690.89 | 17.155 | 17.081 | 1.004332 | |
| *NGF-treated* | 21946.06 | 24842.38 | 17.732 | 18.701 | 0.948185 | |
| *t-BG-treated* | 21859.82 | 23036.33 | 17.663 | 17.341 | 1.018569 | |
| *t-BG+NGF-treated* | 23734.06 | 20747.04 | 19.177 | 15.618 | 1.227878 | |
| *(-) t-BG* | 16541.28 | 19311.89 | 13.365 | 14.538 | 0.919315 | |

*Mef2c Western blot replica for 2-hour treatment*

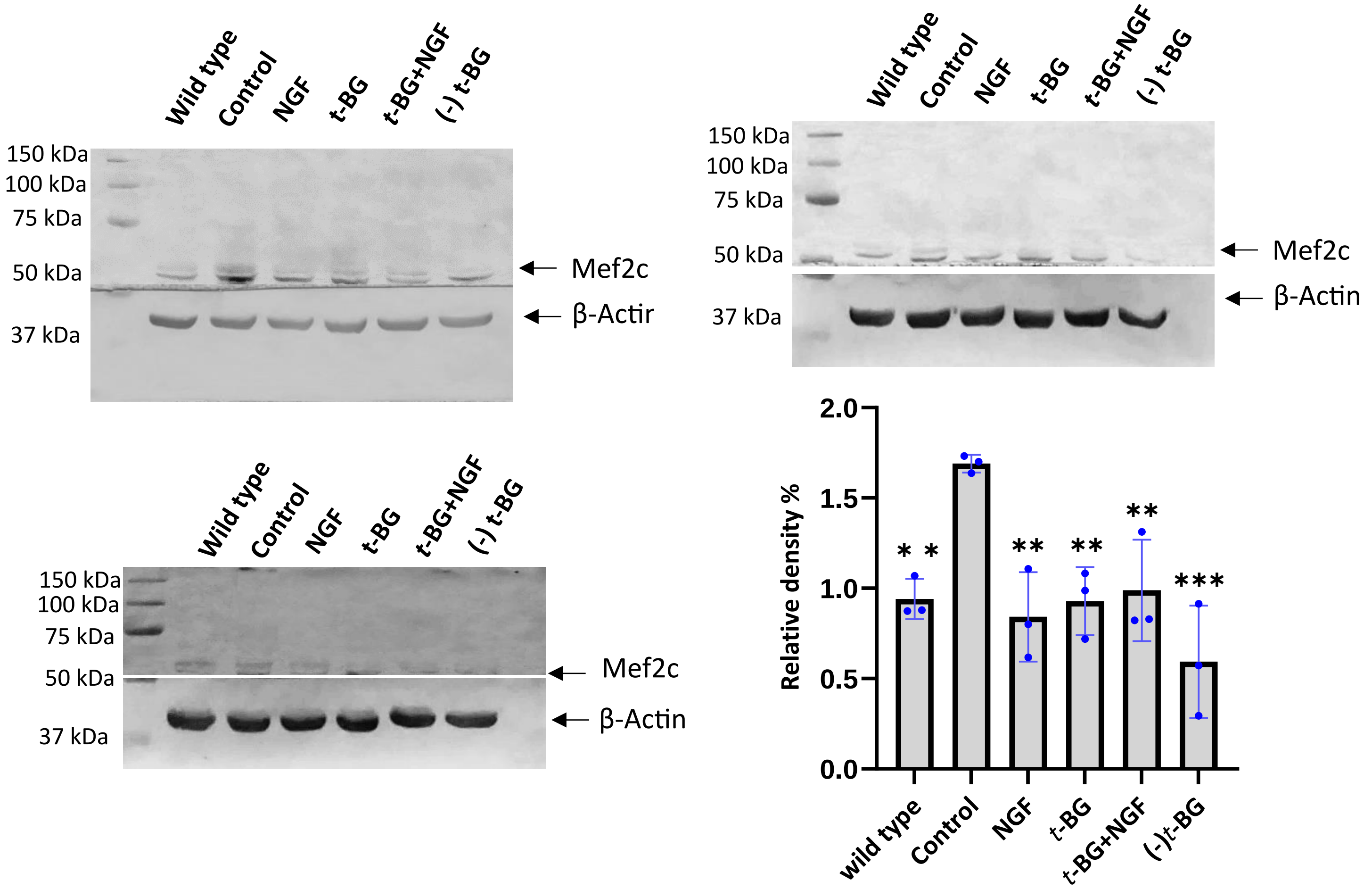

**Table 33: Mef2c calculations for the replica *without* type for 2-hour treatment**

| *Sample name* | *Mef2c band area* | *Actin band area* | *Mef2c area%* | *Actin area%* | *Relative density% (Mef2c area%/ Actin area%)* | |
|---|---|---|---|---|---|---|
| *Control* | 6036.468 | 49602.64 | 32.362 | 20.082 | 1.611493 | Replicate 1 |
| *NGF-treated* | 3736.61 | 49746.43 | 20.032 | 20.14 | 0.994638 | |
| *t-BG-treated* | 3100.903 | 43130.46 | 16.624 | 17.462 | 0.95201 | |
| *t-BG+NGF-treated* | 3241.894 | 50562.1 | 17.38 | 20.47 | 0.849047 | |
| *(-) t-BG* | 2537.154 | 53959.61 | 13.602 | 21.846 | 0.622631 | |
| *Control* | 6855.693 | 42349.39 | 35.898 | 22.706 | 1.580992 | Replicate 2 |
| *NGF-treated* | 2345.832 | 41856.34 | 12.283 | 22.441 | 0.547346 | |
| *t-BG-treated* | 3820.51 | 36663.08 | 20.005 | 19.657 | 1.017704 | |
| *t-BG+NGF-treated* | 5024.501 | 36458.56 | 26.309 | 19.547 | 1.345935 | |
| *(-) t-BG* | 1051.276 | 29185.78 | 5.505 | 15.648 | 0.351802 | |
| *Control* | 4379.46 | 35060.45 | 28.095 | 18.799 | 1.494494 | Replicate 3 |
| *NGF-treated* | 2418.702 | 36217.16 | 15.516 | 19.419 | 0.799011 | |
| *t-BG-treated* | 2681.61 | 40419.96 | 17.203 | 21.672 | 0.793789 | |
| *t-BG+NGF-treated* | 3414.3 | 40125.35 | 21.903 | 21.514 | 1.018081 | |
| *(-) t-BG* | 2694.158 | 34682.98 | 17.284 | 18.596 | 0.929447 | |

**Table 34: Mef2c calculations for the replica *with* type for 2-hour treatment**

| *Sample name* | *Mef2c band area* | *Actin band area* | *Mef2c area%* | *Actin area%* | *Relative density% (Mef2c area%/ Actin area%)* | |
|---|---|---|---|---|---|---|
| *Wild type* | 3350.459 | 47006.26 | 15.295 | 17.385 | 0.879781 | Replicate 1 |
| *Control* | 6099.296 | 44261.86 | 27.843 | 16.37 | 1.700855 | |
| *NGF-treated* | 4015.924 | 44760.28 | 18.333 | 16.554 | 1.107466 | |
| *t-BG-treated* | 3085.146 | 38577.1 | 14.084 | 14.268 | 0.987104 | |
| *t-BG+NGF-treated* | 2999.066 | 44988.03 | 13.691 | 16.639 | 0.822826 | |
| *(-) t-BG* | 2356.033 | 50788.9 | 10.755 | 18.784 | 0.572562 | |
| *Wild type* | 3646.045 | 37755.94 | 14.794 | 16.932 | 0.87373 | Replicate 2 |
| *Control* | 7510.936 | 41509.74 | 30.476 | 18.615 | 1.637174 | |
| *NGF-treated* | 2829.56 | 41488.64 | 11.481 | 18.605 | 0.617092 | |
| *t-BG-treated* | 4398.409 | 36756.37 | 17.846 | 16.483 | 1.082691 | |
| *t-BG+NGF-treated* | 5327.744 | 36716.69 | 21.617 | 16.466 | 1.312826 | |
| *(-) t-BG* | 933.104 | 28763.95 | 3.786 | 12.899 | 0.293511 | |
| *Wild type* | 2603.095 | 40026.58 | 18.883 | 17.654 | 1.069616 | Replicate 3 |
| *Control* | 3683.217 | 34965.92 | 26.717 | 15.422 | 1.732395 | |
| *NGF-treated* | 1762.933 | 36201.73 | 12.789 | 15.967 | 0.800964 | |
| *t-BG-treated* | 1769.225 | 40483.72 | 12.834 | 17.856 | 0.71875 | |
| *t-BG+NGF-treated* | 2022.581 | 40082.99 | 14.672 | 17.679 | 0.829911 | |
| *(-) t-BG* | 1944.631 | 34962.98 | 14.106 | 15.421 | 0.914727 | |

*Ndufs1 western blot replica 2-hour treatment*

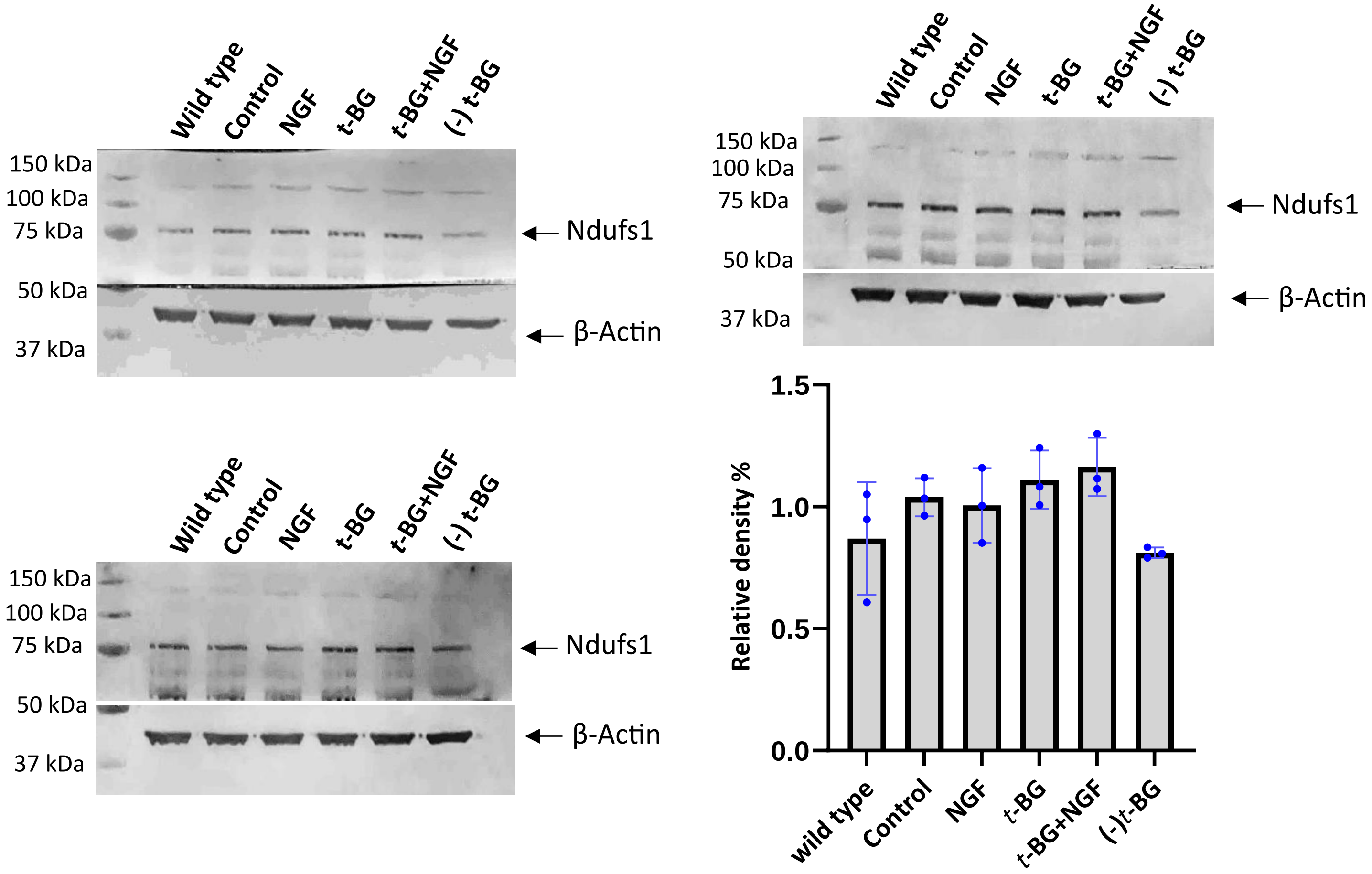

**Table 34: Ndufs1 calculations for the replica *without* type for 2-hour treatment**

| *Sample name* | *Ndufs1 band area* | *Actin band area* | *Ndufs1 area%* | *Actin area%* | *Relative density% (Ndufs1 area%/ Actin area%)* | |
|---|---|---|---|---|---|---|
| *Control* | 12772.97 | 29551.62 | 22.271 | 20.405 | 1.091448 | Replicate 1 |
| *NGF-treated* | 12353.56 | 31153.45 | 21.54 | 21.511 | 1.001348 | |
| *t-BG-treated* | 13055.6 | 32662.89 | 22.764 | 22.554 | 1.009311 | |
| *t-BG+NGF-treated* | 12153.51 | 29067.64 | 21.191 | 20.071 | 1.055802 | |
| *(-) t-BG* | 7015.731 | 22388.38 | 12.233 | 15.459 | 0.791319 | |
| *Control* | 9057.125 | 24947.97 | 19.728 | 21.033 | 0.937955 | Replicate 2 |
| *NGF-treated* | 10434.9 | 25344.04 | 22.729 | 21.367 | 1.063743 | |
| *t-BG-treated* | 9421.075 | 24702.87 | 20.521 | 20.826 | 0.985355 | |
| *t-BG+NGF-treated* | 10347.08 | 22162.6 | 22.538 | 18.685 | 1.206208 | |
| *(-) t-BG* | 6648.903 | 21455.53 | 14.483 | 18.089 | 0.800652 | |
| *Control* | 7059.731 | 14664.41 | 17.876 | 18.168 | 0.983928 | Replicate 3 |
| *NGF-treated* | 6630.338 | 15618.36 | 16.789 | 19.35 | 0.867649 | |
| *t-BG-treated* | 9211.267 | 15762.12 | 23.324 | 19.528 | 1.194388 | |
| *t-BG+NGF-treated* | 9472.468 | 17178 | 23.985 | 21.283 | 1.126956 | |
| *(-) t-BG* | 7119.317 | 17490.61 | 18.027 | 21.67 | 0.831887 | |

**Table 35: Ndufs1 calculations for the replica *with* type for 2-hour treatment**

| *Sample name* | *Ndufs1 band area* | *Actin band area* | *Ndufs1 area%* | *Actin area%* | *Relative density% (Ndufs1 area%/ Actin area%)* | |
|---|---|---|---|---|---|---|
| *Wild type* | 11497.63 | 30722.57 | 16.546 | 17.45 | 0.948195 | Replicate 1 |
| *Control* | 12995.97 | 29425.62 | 18.702 | 16.713 | 1.119009 | |
| *NGF-treated* | 12370.68 | 31220.04 | 17.802 | 17.733 | 1.003891 | |
| *t-BG-treated* | 13062.65 | 32866.54 | 18.798 | 18.668 | 1.006964 | |
| *t-BG+NGF-treated* | 12247.1 | 28897.64 | 17.624 | 16.414 | 1.073718 | |
| *(-) t-BG* | 7316.267 | 22926.79 | 10.528 | 13.022 | 0.808478 | |
| *Wild type* | 8244.409 | 25094.48 | 10.653 | 17.508 | 0.608465 | Replicate 2 |
| *Control* | 6946.61 | 24629.14 | 17.772 | 17.184 | 1.034218 | |
| *NGF-treated* | 6578.51 | 25371.74 | 20.521 | 17.702 | 1.159248 | |
| *t-BG-treated* | 9494.752 | 24640.45 | 18.606 | 17.192 | 1.082248 | |
| *t-BG+NGF-treated* | 9443.054 | 21816.02 | 19.778 | 15.221 | 1.299389 | |
| *(-) t-BG* | 6876.075 | 21776.95 | 12.669 | 15.194 | 0.833816 | |
| *Wild type* | 8244.409 | 15828.7 | 17.326 | 16.485 | 1.051016 | Replicate 3 |
| *Control* | 6946.61 | 14550.87 | 14.599 | 15.154 | 0.963376 | |
| *NGF-treated* | 6578.51 | 15577.36 | 13.825 | 16.223 | 0.852185 | |
| *t-BG-treated* | 9494.752 | 15422.29 | 19.954 | 16.061 | 1.242388 | |

| | | | | | |
|---|---|---|---|---|---|
| *t-BG+NGF-treated* | 9443.054 | 17086.17 | 19.845 | 17.794 | 1.115264 |
| *(-) t-BG* | 6876.075 | 17556.32 | 14.451 | 18.284 | 0.790363 |

*NfκB Western blot replica for 2-hour treatment*

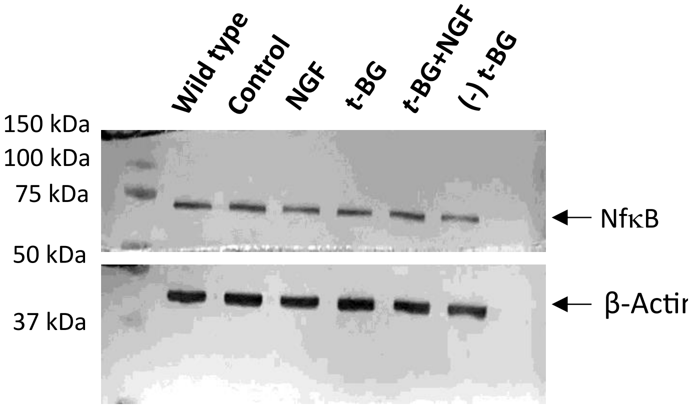


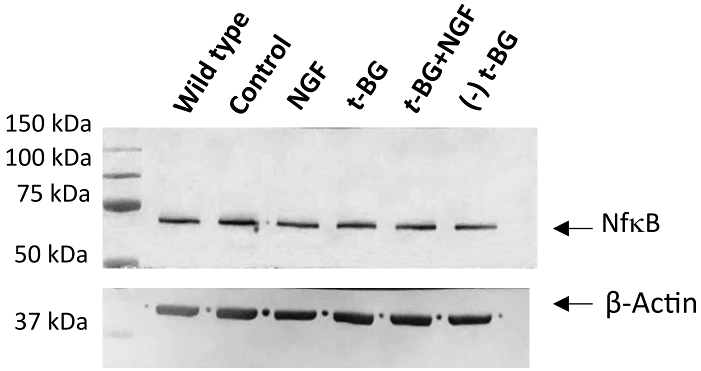


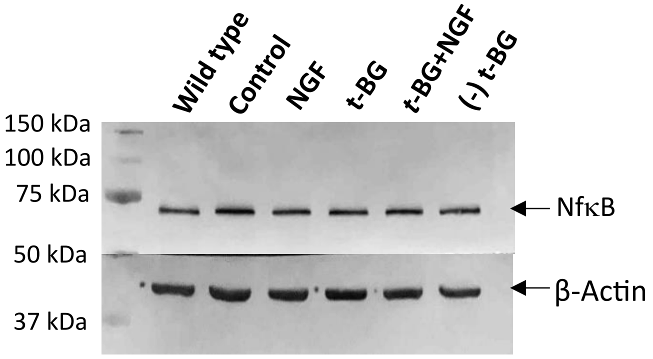


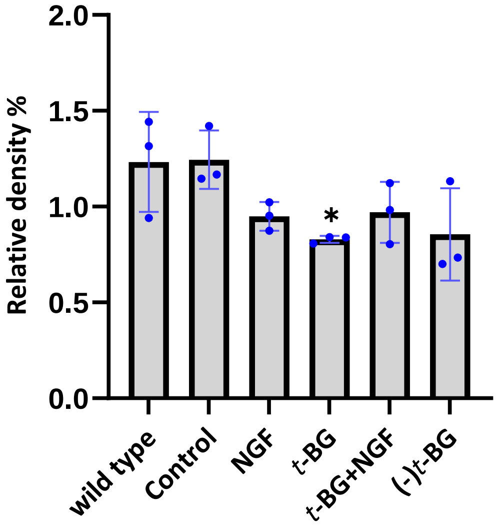

**Table 36: NfκB calculations for the replica *without* type for 2-hour treatment**

| *Sample name* | *NfκB band area* | *Actin band area* | *NfκB area%* | *Actin area%* | *Relative density% (NfκB area%/ Actin area%)* | |
|---|---|---|---|---|---|---|
| *Control* | 12438.97 | 25021.82 | 25.849 | 21.08 | 1.226233 | Replicate 1 |
| *NGF-treated* | 8863.489 | 22993.39 | 18.419 | 19.371 | 0.950854 | |
| *t-BG-treated* | 9429.489 | 25886.38 | 19.595 | 21.808 | 0.898523 | |
| *t-BG+NGF-treated* | 10668.29 | 22852.67 | 22.169 | 19.252 | 1.151517 | |
| *(-) t-BG* | 6722.296 | 21947.04 | 13.969 | 18.489 | 0.75553 | |
| *Control* | 19754.91 | 23523.74 | 28.367 | 19.271 | 1.472005 | Replicate 2 |
| *NGF-treated* | 13934.33 | 22452.75 | 20.009 | 18.393 | 1.08786 | |
| *t-BG-treated* | 12636.95 | 26103.35 | 18.146 | 21.384 | 0.848578 | |
| *t-BG+NGF-treated* | 13377.77 | 26796.04 | 19.21 | 21.951 | 0.875131 | |
| *(-) t-BG* | 9936.439 | 23194.02 | 14.268 | 19.001 | 0.750908 | |
| *Control* | 18707.42 | 25784.71 | 25.457 | 22.252 | 1.144032 | Replicate 3 |
| *NGF-treated* | 13259.55 | 22724.09 | 18.043 | 19.61 | 0.920092 | |
| *t-BG-treated* | 13756.14 | 25479.77 | 18.719 | 21.989 | 0.851289 | |
| *t-BG+NGF-treated* | 14163.09 | 22811.45 | 19.273 | 19.686 | 0.979021 | |
| *(-) t-BG* | 13600.43 | 19077.6 | 18.507 | 16.464 | 1.124089 | |

**Table 37: NfκB calculations for the replica *with* type for 2-hour treatment**

| *Sample name* | *NfκB band area* | *Actin band area* | *NfκB area%* | *Actin area%* | *Relative density% (NfκB area%/ Actin area%)* | |
|---|---|---|---|---|---|---|
| *Wild type* | 11067.9 | 20133.87 | 19.309 | 14.675 | 1.315775 | Replicate 1 |
| *Control* | 11884.27 | 24809.7 | 20.733 | 18.083 | 1.146546 | |
| *NGF-treated* | 8426.368 | 23091.51 | 14.7 | 16.83 | 0.87344 | |
| *t-BG-treated* | 9065.075 | 25815.67 | 15.815 | 18.816 | 0.840508 | |
| *t-BG+NGF-treated* | 10355.87 | 22082.72 | 18.067 | 16.095 | 1.122523 | |
| *(-) t-BG* | 6521.004 | 21269.09 | 11.376 | 15.502 | 0.733841 | |
| *Wild type* | 13705.14 | 15849.24 | 16.64 | 11.537 | 1.442316 | Replicate 2 |
| *Control* | 19685.61 | 23121.38 | 23.901 | 16.831 | 1.420058 | |
| *NGF-treated* | 13654.38 | 22260.46 | 16.578 | 16.204 | 1.023081 | |
| *t-BG-treated* | 12664.95 | 26166.77 | 15.377 | 19.048 | 0.807276 | |
| *t-BG+NGF-treated* | 12848.24 | 26621.62 | 15.599 | 19.379 | 0.804943 | |
| *(-) t-BG* | 9805.146 | 23355.31 | 11.905 | 17.001 | 0.700253 | |
| *Wild type* | 12307.36 | 20658.5 | 14.305 | 15.195 | 0.941428 | Replicate 3 |
| *Control* | 18790.71 | 25427.59 | 21.84 | 18.703 | 1.167727 | |
| *NGF-treated* | 13522.21 | 22446.79 | 15.717 | 16.511 | 0.951911 | |
| *t-BG-treated* | 13555.6 | 25546.89 | 15.755 | 18.791 | 0.838433 | |
| *t-BG+NGF-treated* | 14227.62 | 22863.16 | 16.536 | 16.817 | 0.983291 | |
| *(-) t-BG* | 13634.43 | 19011.19 | 15.847 | 13.984 | 1.133224 | |

*Pkc Western blot replica for 2-hour treatment*

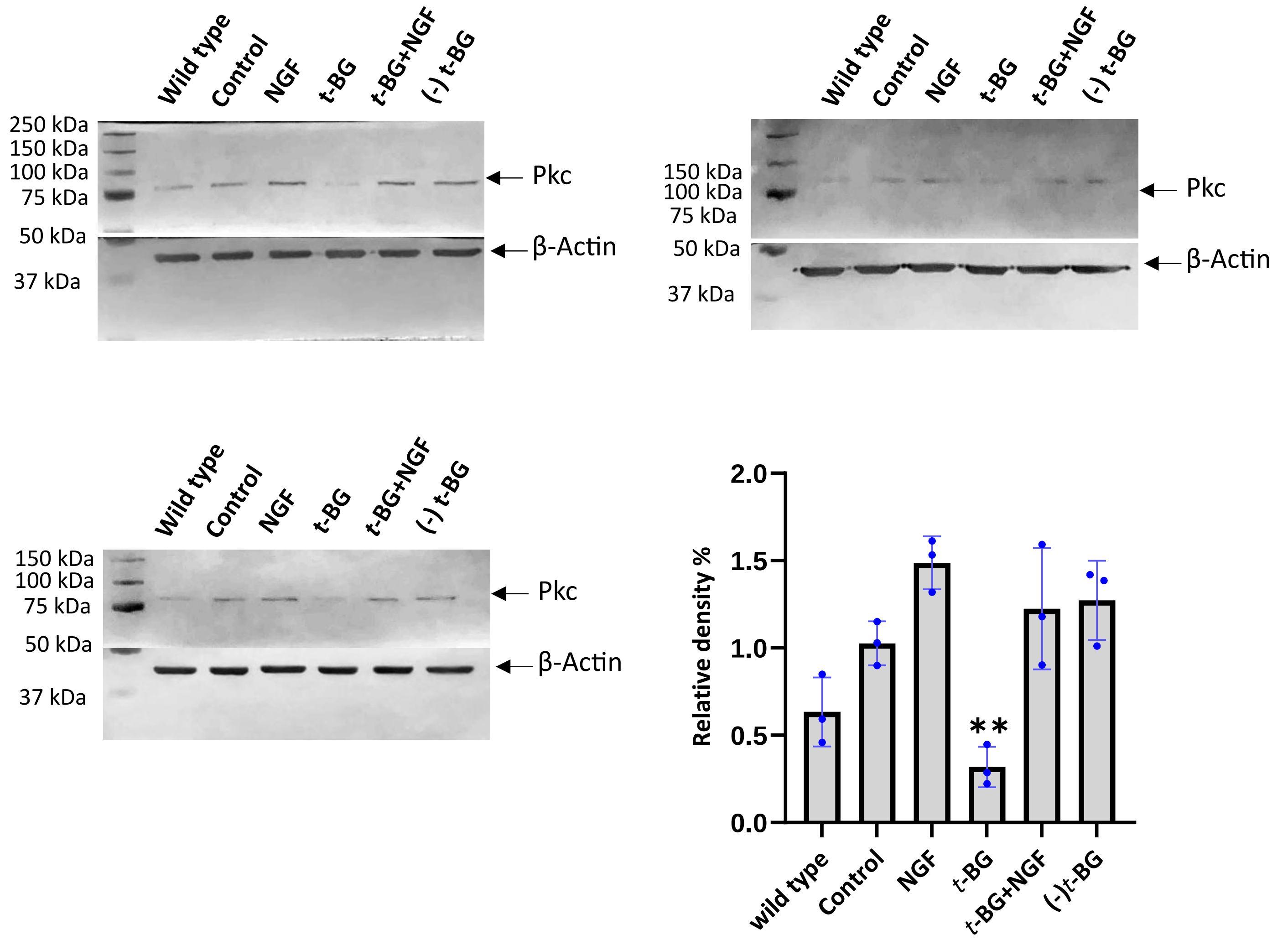

**Table 38: Pkc calculations for the replica *without* type for 2-hour treatment**

| *Sample name* | *Pkc band area* | *Actin band area* | *Pkc area%* | *Actin area%* | *Relative density% (Pkc area%/ Actin area%)* | |
|---|---|---|---|---|---|---|
| *Control* | 7469.146 | 33621.26 | 19.567 | 20.387 | 0.959778 | Replicate 1 |
| *NGF-treated* | 12459.33 | 34480.87 | 32.64 | 20.908 | 1.561125 | |
| *t-BG-treated* | 3287.347 | 34702.6 | 8.612 | 21.042 | 0.409277 | |
| *t-BG+NGF-treated* | 6482.51 | 29896.53 | 16.982 | 18.128 | 0.936783 | |
| *(-) t-BG* | 8473.853 | 32215.95 | 22.199 | 19.535 | 1.136371 | |
| *Control* | 7195.167 | 37437.54 | 19.078 | 19.275 | 0.98978 | Replicate 2 |
| *NGF-treated* | 10603.7 | 40052.89 | 28.116 | 20.621 | 1.363464 | |
| *t-BG-treated* | 1889.589 | 37376.03 | 5.01 | 19.243 | 0.260354 | |
| *t-BG+NGF-treated* | 8268.48 | 39974.46 | 21.924 | 20.581 | 1.065254 | |
| *(-) t-BG* | 9757.702 | 39391.95 | 25.872 | 20.281 | 1.275677 | |
| *Control* | 6334.782 | 19146.6 | 16.564 | 21.155 | 0.782983 | Replicate 3 |
| *NGF-treated* | 9531.338 | 18362.41 | 24.923 | 20.288 | 1.22846 | |
| *t-BG-treated* | 1433.598 | 16801.68 | 3.749 | 18.564 | 0.20195 | |
| *t-BG+NGF-treated* | 10224.22 | 16479.41 | 26.734 | 18.208 | 1.468256 | |
| *(-) t-BG* | 10719.68 | 19716.87 | 28.03 | 21.785 | 1.286665 | |

**Table 39: Pkc calculations for the replica *with* type for 2-hour treatment**

| *Sample name* | *Pkc band area* | *Actin band area* | *Pkc area%* | *Actin area%* | *Relative density% (Pkc area%/ Actin area%)* | |
|---|---|---|---|---|---|---|
| *Wild type* | 5033.731 | 16380.16 | 11.662 | 13.734 | 0.849134 | Replicate 1 |
| *Control* | 7449.439 | 17895.74 | 17.259 | 15.005 | 1.150217 | |
| *NGF-treated* | 12209.62 | 20926.19 | 28.287 | 17.546 | 1.612162 | |
| *t-BG-treated* | 3307.589 | 20414.5 | 7.663 | 17.117 | 0.447684 | |
| *t-BG+NGF-treated* | 6717.51 | 20573.26 | 15.563 | 17.25 | 0.902203 | |
| *(-) t-BG* | 8445.439 | 23073.33 | 19.566 | 19.347 | 1.01132 | |
| *Wild type* | 2873.004 | 35535.77 | 7.042 | 15.327 | 0.459451 | Replicate 2 |
| *Control* | 6847.974 | 37794.96 | 16.786 | 16.301 | 1.029753 | |
| *NGF-treated* | 10724.29 | 39790.06 | 26.288 | 17.162 | 1.531756 | |
| *t-BG-treated* | 1908.468 | 37980.81 | 4.678 | 16.381 | 0.285575 | |
| *t-BG+NGF-treated* | 8511.945 | 41005.17 | 20.865 | 17.686 | 1.179747 | |
| *(-) t-BG* | 9929.995 | 39747.36 | 24.341 | 17.143 | 1.41988 | |
| *Wild type* | 4703.974 | 19962.65 | 11.065 | 18.077 | 0.612104 | Replicate 3 |
| *Control* | 6405.489 | 18859.48 | 15.068 | 17.078 | 0.882305 | |
| *NGF-treated* | 9589.631 | 18737.53 | 22.558 | 16.967 | 1.329522 | |
| *t-BG-treated* | 1350.598 | 16906.39 | 3.177 | 15.309 | 0.207525 | |
| *t-BG+NGF-treated* | 9758.681 | 16168.58 | 22.955 | 14.641 | 1.567857 | |
| *(-) t-BG* | 10703.56 | 19797.29 | 25.178 | 17.927 | 1.404474 | |

### 7. FerroOrange assay

| Sample | Absorbance at 534 nm/580 nm | | | | | |
|---|---|---|---|---|---|---|
| | Wild | Control | NGF-treated | *t*-BG-treated | *t*-BG+NGF-treated | (-) *t*-BG |
| Replicate 1 | 2681 | 2903 | 2153 | 1101 | 1836 | 1549 |
| Replicate 2 | 1643 | 3197 | 3317 | 1282 | 1978 | 1960 |
| Replicate 3 | 2167 | 2349 | 3613 | 1006 | 1937 | 1525 |

Collagen reading was deducted from each sample.

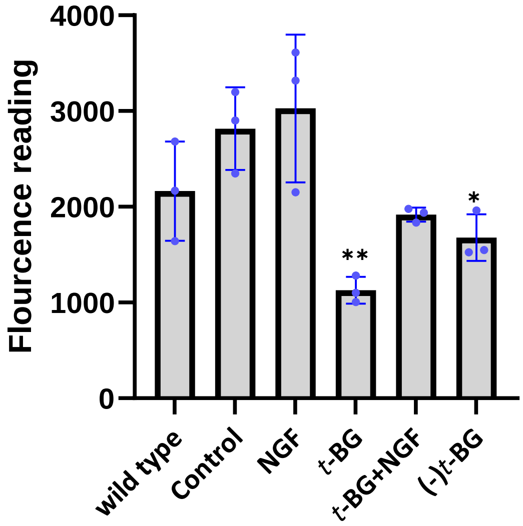

### 8. NMR spectra

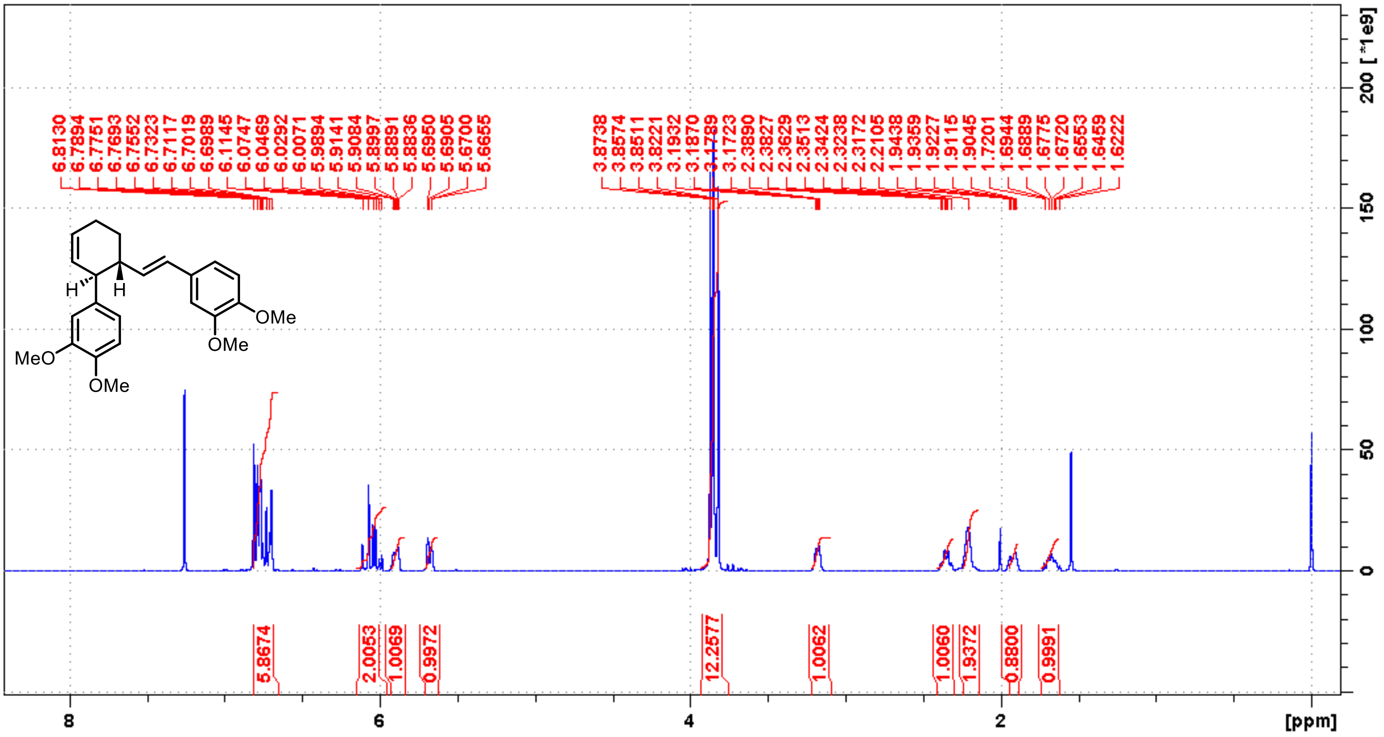


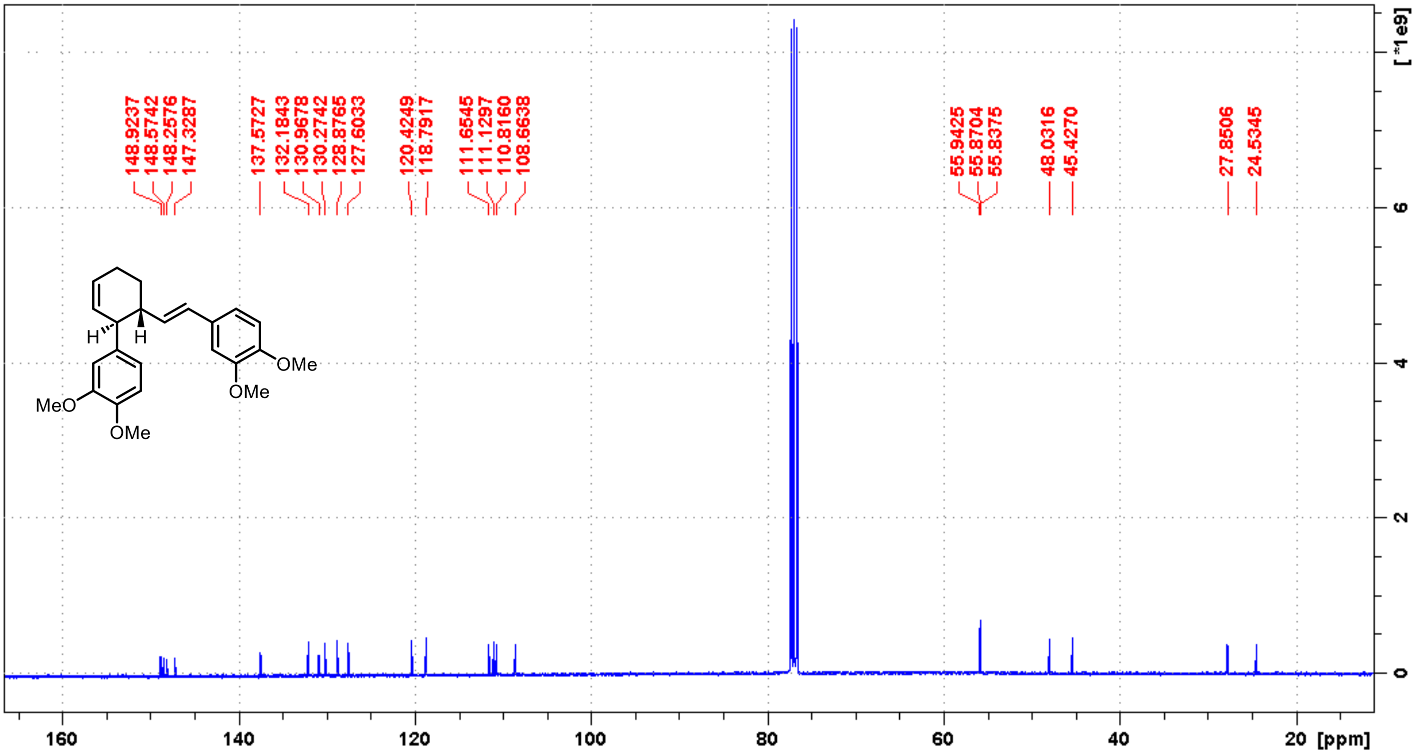

***t*–BG- 3(S/R)-(3,4-Dimethoxyphenyl)-4(R/S)-[(*E*)-3,4-dimethoxystyryl]cyclohex-1-ene:** light yellow oil, HPLC injection amount: 300 mg, yield after HPLC: 129 mg, **43%**. Spectral data are consistent with Gohil *et al* (2022) work .[2] **$^{1}$H NMR** ($CDCl_3$, 400MHz) δ: 6.82-6.7 (m, 6H), 6.09 (d, J = 16.1 Hz, 1H), 6.02 (dd, J = 7.1 Hz, 16.1 Hz), 5.90 (tdd, J = 2.3 Hz, 4.6 Hz, 10.1 Hz, 1H), 5.69-5.67 (dq, J = 10.3 Hz, 2.0 Hz, 1H), 3.87 (s, 3H), 3.86 (s, 3H), 3.85 (s, 3H), 3.82 (s, 3H), 3.19-3.17 (m, 1H), 2.39-2.32 (qd, J = 9.4 Hz, 2.8 Hz 1H), 2.25–2.18 (m, 2H), 1.94-1.90 (m, 1H), 1.72–1.62 (m, 1H). **$^{13}$C NMR** ($CDCl_3$, 176 MHz) δ: 148.9, 148.6, 148.3, 147.3, 137.6, 132.2, 131.0, 130.3, 128.9, 127.6, 120.4, 118.8, 111.7, 111.1, 110.8, 108.7, 56.0, 55.9, 55.9, 55.8, 48.0, 45.4, 27.9, 24.5

**References**

1. Stael, S.; Miller, L. P.; Fernández-Fernández Á, D.; Van Breusegem, F., Detection of Damage-Activated Metacaspase Activity by Western Blot in Plants. *Methods Mol Biol* **2022,** *2447*, 127-137.
2. Gohil, K.; Kazmi, M. Z. H.; Williams, F. J., Structure-activity relationship and bioactivity studies of neurotrophic trans-banglene. *Organic & Biomolecular Chemistry* **2022,** *20* (11), 2187-2193.
